# Haplotype-level metabarcoding revealed the relationship between species occurrence and population genetic structure at the catchment scale in aquatic insects

**DOI:** 10.64898/2026.01.13.699187

**Authors:** Dan Joseph C. Logronio, Ming-Chih Chiu, Arnelyn D. Doloiras-Laraño, Levente-Péter Kolcsár, Joeselle M. Serrana, Kohei Hamamoto, Kozo Watanabe

## Abstract

A longstanding question in ecology and evolutionary biology is understanding the evolutionary mechanisms whereby some species appear frequently while others do not. Current research examined this phenomenon through computer simulations and empirical studies involving a few species, while community-wide investigations of this relationship in natural settings remain largely unexplored. To address this, we employed haplotype-level bulk DNA metabarcoding to infer haplotypes from many aquatic insect species with varying occurrence in a catchment. We identified 891 haplotypes belonging to 200 taxonomically identified Operational Taxonomic Units (OTUs). Results showed that species with higher occurrence in the catchment had greater ɣ-genetic diversity, a wider range of environmental conditions, and an increased probability of β-species-genetic diversity correlations (SGDC). We found negative α-SGDC in species with lower occurrence in the catchment. Dispersal abilities and environmental factors may influence the relationship among species occurrence, population genetic structure, and SGDC. Our findings present a novel approach to address the association of species occurrence and population genetic structure, as well as SGDC, from a community-level perspective. This has important implications for addressing biodiversity loss driven by rapid environmental changes from human activities and climate change.

## Introduction

The evolutionary mechanisms that determine why some species are widely distributed while others are not have long been debated among ecologists and evolutionary biologists (Loveless and Hamrick, 1984; Gaston and Kunin, 1997; Holt, 2003). Species occurrence is defined as the presence or absence of a species at a specific location (Tingley and Beissinger, 2009; Gaston and He, 2011). It is commonly used to estimate species’ ranges and distributions (Fourcade, 2016; Guillera-Arroita, 2017). Linking species occurrence with population genetic data can provide more realistic insights into changes in species range (Gotelli and Shanton-Geddes, 2015). Previous theoretical and empirical studies have examined the evolutionary mechanisms underlying species’ ranges and distributions. However, these studies were based on computational simulations and a small number of test species (Gitzendanner and Soltis, 2000; Pierce et al., 2017; Polechová, 2018). These studies do not provide a comprehensive understanding of species’ ranges and distributions due to limitations in mathematical models and the small number of species examined. Therefore, to achieve a more robust and generalizable conclusion about the evolutionary mechanisms underlying species range and distribution, it is imperative to examine many species with varying occurrences and population genetic structures.

Species with higher occurrence are more likely to occupy a broader range of environmental conditions than those with lower occurrence. Previous research has shown that environmental heterogeneity increases with area (Darlington, 1943; Gentile et al., 2021); thus, species occurring at many sites likely experience a wider range of environmental conditions. Consequently, high environmental heterogeneity increases genetic differentiation among populations (β-genetic diversity) and total genetic diversity (ɣ-genetic diversity) within a landscape (Vellend and Geber 2005; Fortune et al. 2016; Bonte and Bafort 2019) through adaptive divergence. Therefore, species with higher occurrence plausibly have higher β- and ɣ-genetic diversity than those with lower occurrence. However, these links have yet to be examined across many species with different occurrences and population genetic structures.

To fully understand these relationships, population genetic analyses must be conducted across many taxa with different occurrences. Traditional methods based on Sanger sequencing are often limited by cost, labor, and time, making large-scale, multi-species population genetic analysis challenging. High-throughput sequencing techniques, such as haplotype-level DNA metabarcoding, are now attracting interest. These methods allow both species identification and the measurement of intraspecific genetic variation. As a result, population genetic analysis can be conducted across numerous taxa (Elbrecht et al. 2018; Turon et al. 2020; Serrana and Watanabe 2023). An added advantage is that this approach compares population genetic structures across multiple species and provides information about the entire community. This enables simultaneous testing of ecological and evolutionary concepts, such as Species-Genetic Diversity Correlations (SGDCs), across many species. SGDC has two types: α- and β-SGDC. The α-SGDC is the correlation between local genetic and species diversities. The β-SGDC refers to the correlation between population genetic differentiation and community divergence (Kahilainen et al. 2014). Species with high occurrence are more likely to show positive α- and β-SGDC, compared to those with low occurrence, due to a wider range of environmental conditions that promote adaptive divergence. Vellend and Geber (2005) and Fortune et al. (2016) found that environmental heterogeneity promotes diversifying selection and adaptive divergence at both the population and community levels. This drives positive β-SGDC. However, recent studies show that environmental heterogeneity can have opposite effects on these two levels of biodiversity (Martin et al. 2021). More empirical research is needed to clarify the relationships among species occurrence, environmental heterogeneity, and SGDC. Addressing this issue with a multi-species approach can aid SGDC-based conservation, since species’ ecological differences strongly impact SGDC (Seymour et al. 2016).

We conducted a case study using haplotype-level bulk DNA metabarcoding on aquatic insects to address key questions in ecology and evolution, focusing on the relationships between species occurrence, population genetic structure, and SGDC at the community level. Aquatic insects were chosen for their species diversity and ubiquity (Dijkstra et al. 2014; Starr and Wallace 2021). We tested whether species with higher occurrence in the catchment (1) occupy a broader range of environments and show greater ɣ-genetic diversity, (2) possess higher β-genetic diversity, and (3) are more likely to exhibit positive α- and β-SGDC.

## Methods

### Study site and sample collection

Sampling was conducted from May 6 to 12, 2021, in the Hiji River, Shikoku Island, Japan. The river originates from Torisaka Pass in Seiyo City, Ehime Prefecture, and flows into the Iyo Sea. The watershed area is 1,210 km². Sixty sites across different environmental conditions were selected for sampling (Fig. 1). We collected three samples (upstream, middle, downstream) per site, which we processed separately for sorting, DNA extraction, and 1st PCR, but pooled for the 2nd PCR. Macroinvertebrates were collected using a kick-net method (0.5 mm mesh). The collected macroinvertebrates were preserved in 99.5% ethanol and transported to the laboratory for sorting. Sorted macroinvertebrates were stored in 99.5% ethanol at 4°C until DNA extraction. Environmental parameters, including water temperature, pH, electrical conductivity, and dissolved oxygen, were measured in situ using a portable meter (LAQUA, Horiba Corporation, Japan). Surface water was also collected for the measurement of ammonium nitrogen (NH_4_-N), nitrate-nitrogen (NO_3_-N), nitrite-nitrogen (NO_2_-N), and phosphate-phosphorus (PO_4_-P) using the QuAAtro 2-HR system (BeLtech).

**Figure 1.**
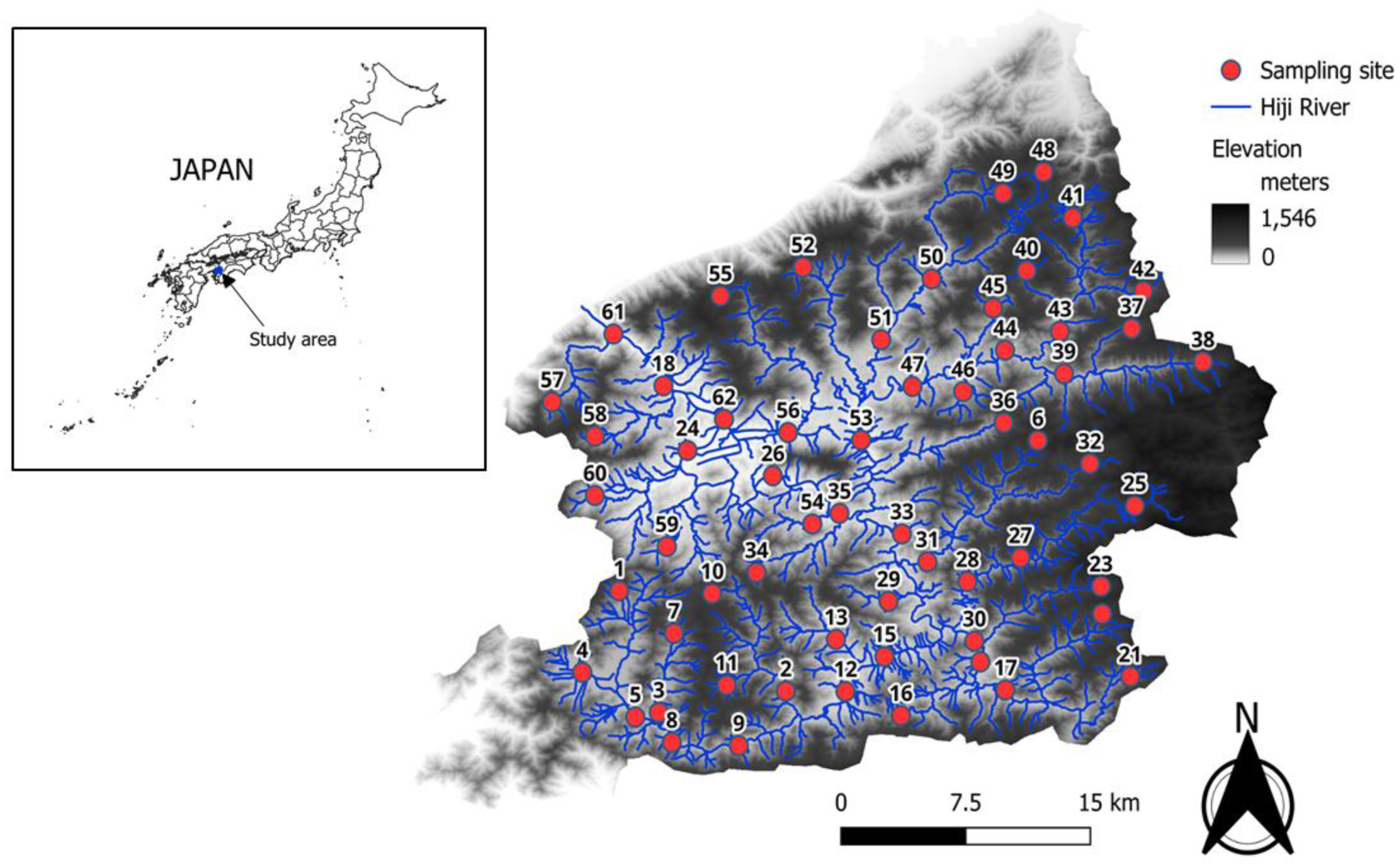
Location of sampling sites in Hiji River (Ehime, Japan).

### DNA extraction, library preparation, and sequencing

We extracted DNA from ethanol-dried macroinvertebrate bulk community samples using the Phenol-Chloroform-Isoamyl Alcohol (PCI) method (supplementary protocol 1). DNA was quantified using a NanoDrop spectrophotometer (NanoDrop 2000, Thermo Scientific, USA), quality-checked by agarose gel electrophoresis, and diluted to 20 ng/uL. We performed PCR amplification of the mitochondrial cytochrome oxidase subunit 1 (COI) gene using the primer set BF2 (5’-GCHCCHGAYATRGCHTTYCC-3’) and BR2 (5’-TCDGGRTGNCCRAARAAYCA-3’) (Elbrecht and Leese 2017). A two-step library preparation method was employed following Elbrecht and Steinke (2019) with modifications (supplementary protocol 2). We included negative (molecular-grade water) and positive controls (macroinvertebrate bulk-community DNA) in all PCR assays. We quantified the libraries using the KAPA Library Quantification Kit (Kapa Biosystems, USA) in a C1000 Touch Real-Time System Thermal cycler (Bio-Rad Laboratories, CA, USA). Then, we pooled each sample in equimolar amounts (4 nM) and cleaned the pooled libraries using SPRIselect (Beckman Coulter Inc., USA) with a left-sided selection ratio of 0.85X and checked the quality using the Agilent Bioanalyzer 2100 system (Agilent Biotechnologies). High-throughput sequencing was performed on an Illumina MiSeq system with a 300 bp paired-end read kit v3 (Illumina, USA, MS-102-3003) at a final concentration of 8 pM with a 10% Phi-X spike-in.

### Read processing and generation of haplotypes

Raw paired-end reads were quality-checked using FastQC (Andrews 2010) and then demultiplexed via JAMP (Elbrecht et al. 2018) in R v.4.3.2 (http://github.com/vascoElbrecht/JAMP). After demultiplexing, the reads were processed through the DADA2 pipeline (Callahan et al. 2016), which included primer removal (trimleft = 20, 20), trimming of forward and reverse reads (trunclen = 260, 220), filtering of erroneous reads (maxEE = 1, 3), denoising, merging of forward and reverse reads, and removal of chimeras. Amplicon Sequence Variants (ASVs) were further filtered to extract haplotypes. ASVs were filtered for the expected amplicon size (421 bp) using Cutadapt version 4.9 (Martin 2011), and sequences with fewer than 10 reads per sample were removed. We used Multiple Alignment of Coding Sequences (MACSE) version 2.07 (Ranwez et al. 2011; Ranwez et al. 2020) to assess the presence of internal frame shift, stop codons, and deletions. ASVs were then translated into protein sequences using Geneious Prime version 2023.0.1 (https://www.geneious.com). We performed pseudogene filtering by comparing the open reading frame (ORF) to the arthropod amino acid COI gene HMM profile using a hidden Markov model (HMM) scan analysis (http://hmmer.org/) (Porter and Hajibabaei 2021). We discarded sequences with a bit score below the first quartile (Q1) – 1.5 × interquartile range (IQR) as putative pseudogenes (Porter and Hajibabaei 2021). ASVs that passed these filtering steps were considered haplotypes.

### OTU Clustering for Species Delineation

In this study, Operational Taxonomic Units (OTUs) were used to define species boundaries. Haplotype sequences were clustered into OTUs with 97% identity using the cluster_smallmem command in VSEARCH v2.21.1 (Rognes et al. 2016). Because OTUs can group haplotype sequences from multiple species (Martin et al., 2021), we screened all OTUs for signals of mixed-species ancestry. Initial screening was conducted based on the maximum pairwise p-distance. OTUs with a maximum pairwise p-distance of < 0.030 were considered single species, whereas those with a maximum pairwise p-distance of > 0.030 were potentially mixed species. OTUs with a maximum pairwise p-distance > 0.030 were further confirmed for mixed-species signals based on two criteria: genetic distance and phylogeny. The genetic distance method involved testing the modality of pairwise p-distance distribution of all haplotypes within the OTU using a histogram and excess mass-based multimodality testing according to Ameijeiras-Alonso et al. (2019), implemented in the multimode package v1.5 in R (Ameijeiras-Alonso et al., 2021). The phylogeny method was based on Poisson Tree Process (PTP) (https://species.h-its.org/ptp/) (Zhang et al., 2013) (supplementary protocol 3). We considered OTUs with a maximum pairwise distance of > 0.03 as single species if the pairwise distance of all haplotypes within the OTU showed a unimodal distribution based on a histogram and excess mass-based multimodality testing and demonstrated a monophyletic clade with strong Maximum Likelihood and Bayesian support values (≥ 75) based on PTP. Otherwise, those that did not meet both criteria were classified as mixed species.

### Taxonomic Assignment of Haplotypes

For taxonomic assignment, haplotype sequences were first queried against the Barcode of Life Database (BOLDv5, http://www.boldsystems.org/; accessed October 2025) using a ≥97% identity cutoff. Sequences that did not return taxonomic assignment using the BOLD database were then queried against the in-house database using the –usearch global command in VSEARCH v2.21.1 (Rognes et al., 2016), with a ≥97% identity threshold. The in-house database consisted of Sanger sequences of morphologically identified adult crane flies and was deposited in the DNA Data Bank of Japan (accession numbers LC898117-LC898137). Sequences that did not have taxonomic identification using BOLD and an in-house database were further blasted against the National Center for Biotechnology Information (NCBI) nonredundant database (accessed October 2025) using the megablast option in Geneious Prime version 2023.0.1 (https://www.geneious.com) at a ≥97% similarity threshold and a minimum query coverage of ≥97%. Final identification in each database was based on the highest percent identity. If multiple hits were obtained with equal percent identity but with different species identification, we downgraded to genus-level identification (Arulandhu et al., 2017). Haplotype sequences without significant hits, non-insect species, and terrestrial insect species were excluded from subsequent analysis. When two or more OTUs share the same species identification, which may indicate cryptic species, we distinguish them by appending a number to the scientific name for the purpose of our analysis (e.g., *Cladotanytarsus vanderwulpi* 1, *Cladotanytarsus vanderwulpi* 2, etc.).

### Genetic diversity analysis

OTUs detected at least five sites were used for population genetic analysis to ensure a minimum of three data points for the correlation analysis. More frequently and less frequently detected OTUs in the catchment were determined based on the number of sites. We classified OTUs identified at ≥ 17 sites (75th percentile) as more frequently detected and those at ≤ 6 sites (25th percentile) as less frequently detected. Then, the haplotype read counts were transformed into presence/absence data. Haplotype richness was based on the total number of unique haplotypes of an OTU in a sample. Global ɸ-statistics (GɸST) among populations within an OTU were calculated using Analysis of Molecular Variance (AMOVA) (Weir and Cockerham, 1984; Excoffier et al., 1992; Weir, 1996) according to the differences in the allelic content of haplotypes (pairwise difference) within and among populations (Excoffier et al., 1992). Pairwise ɸST was also computed using the dissimilarity of allelic content of the haplotypes (pairwise difference) between populations. We also determined the average pairwise ɸST per OTU. Both GɸST and pairwise ɸST were computed using the Arlequin version 3.5.2.2 (Excoffier and Lischer 2010). We classified average pairwise ɸST values as high (≥ 0.3145; 75th percentile) or low (≤ −0.1895; 25th percentile).

To understand the relation and location of haplotypes within an OTU in the catchment, we analyzed their phylogeography and haplotype network. We used PopART (Bandelt et al. 1999) to create the Minimum Spanning Network. We classified haplotype network structures as “star-like,” “complex star,” “reciprocally monophyletic,” or “complex mutational” (Jenkins et al. 2018). Populations with few haplotypes and networks that did not fit these types were described as showing “no pattern.” We defined ancestral haplotypes as those at the network center and found more often in the catchment. For phylogeographic analysis, we mapped all haplotypes within an OTU using QGIS version 3.32.0-Lima (https://qgis.org/en/site).

We used the ggscatter function (ggpubr v0.6.0, Kassambara 2023) to run a Kendall correlation test on the number of sites, ɣ-genetic diversity, average α-genetic diversity, GɸST, and average pairwise ɸST. We made correlation plots with ggplot2 (Wickham 2016) and ggpubr. In the average pairwise ɸST analysis, we excluded OTUs that were genetically homogeneous (one haplotype per species). For GɸST, we excluded OTUs that were either genetically homogeneous or monomorphic (one haplotype per site).

### Species-Genetic Diversity Correlations Analysis

α- and β-SGDCs were analyzed using presence/absence data for OTUs and haplotypes. Specifically, for the α-SGDC, haplotype richness was correlated with community OTU richness. OTU richness was measured as the number of unique OTUs in the community. Kendall correlation tests were performed using the ggscatter function of the ggpubr package (version 0.6.0; Kassambara 2023). For the β-SGDC, population pairwise ɸST was correlated with community Sørensen dissimilarity using Mantel tests with the Kendall method and 999 permutations in the vegan version 2.6-4 (Oksanen et al., 2022). For OTUs with a small number of occurrence sites, the vegan version 2.6-4 (Oksanen et al., 2022) conducted an exact permutation test using all possible data arrangements (Berry et al., 2011). In all SGDC analyses, we included only sites in which the species of interest was detected. Multiple-testing corrections were not performed, and interpretations were based on the complete set of results (Watanabe and Monaghan, 2017).

### Relationship between female wing length with number of sites and genetic diversity

We used female wing length as an estimate of the insect’s dispersal ability. Female wing length was categorized from 1 to 8 (1=shortest or <5 mm; 8 = longest or ≥ 50 mm) according to the DISPERSE database (Sarramejane et al., 2020). We included in the analysis OTUs identified at the genus level, along with the female wing-length data, in DISPERSE. However, for OTUs within Diptera, the data were based on subfamilies and tribes available in DISPERSE. The longest female wing length category for each OTU was correlated with the number of sites, average α-genetic diversity, average pairwise ɸST, and GɸST using the Kendall method implemented in the ggscatter function of the ggpubr package version 0.6.0 (Kassambara 2023).

### Correlation with Environmental Parameters

We used Mantel tests to determine whether geographic distance (river and straight-line distances) significantly affected the pairwise ɸ-statistics for each OTU. River distance is defined as the waterline distance along river corridors. In contrast, straight-line distance is the air distance across the landscape. Mantel statistics were implemented in the vegan package version 2.6-4 (Oksanen et al., 2022) using the Kendall method, with 999 permutations. For OTUs with a small number of occurrence sites, vegan conducted an exact permutation test using all possible data arrangements (Berry et al., 2011). We used the same analysis to test the correlation between geographic distance and community Sørensen dissimilarity. We excluded genetically homogeneous populations from the analysis.

We employed univariate Distance-based Redundancy Analysis (dbRDA) in the vegan package 2.6-4 (Oksanen et al. 2022) with 999 permutations to examine the correlation of eight environmental parameters, namely, pH, conductivity, dissolved oxygen, NH_4_-N, NO_3_-N, NO_2_-N, PO_4_-P, and elevation, with population pairwise ɸST in each OTU. The significance of the capscale model was tested using anova.cca function. The p-values were adjusted using Benjamini-Hochberg false discovery rate (FDR) correction. Univariate dbRDA was used because the less frequently detected OTUs in the catchment had fewer sites than environmental parameters. However, for 16 OTUs with a substantially higher number of sites (≥ 16 sites) than environmental parameters (8 parameters), an additional analysis using ordiR2step with forward selection and 999 permutations in the vegan package 2.6-4 (Oksanen et al. 2022) was conducted to verify results. The same analysis was performed to correlate the environmental parameters with the community Sǿrensen dissimilarity.

We correlated the mean and maximum pairwise differences of environmental parameters with the number of sites, ɣ-genetic diversity, average pairwise ɸST, and GɸST for all 59 OTUs using the Kendall method in the ggscatter function of ggpubr (v0.6.0; Kassambara 2023). We adjusted p-values with the Benjamini-Hochberg FDR correction. For each OTU, the mean and maximum pairwise differences for each environmental parameter were calculated from the environmental distance matrix using all sites where the OTU was detected. We excluded genetically homogeneous OTUs from average pairwise ɸST analysis, and both genetically homogeneous and monomorphic OTUs from GɸST analysis.

## Results

### Metabarcoding data

The sequencing run yielded 6,932,659 demultiplexed reads. We obtained 586,062 reads, comprising 1,879 ASVs, after primer removal, quality filtering, denoising, merging, and chimera removal. After length and abundance filtering, the number of reads was reduced to 526,468, yielding 1,423 ASVs. None of the ASVs contained internal frame shift, stop codons, or deletions according to MACSE analysis. After pseudogene filtering based on HMM scan analysis, 1,297 ASVs (hereafter referred to as haplotypes) were retained. Of these, 918 haplotypes had taxonomic identification at least at the family level. After excluding the non-insect and terrestrial insect sequences, we obtained 891 haplotypes and clustered them into 200 OTUs. Six out of 200 OTUs (3%) were verified to be a mixture of haplotypes from two or more species. We excluded them from downstream population genetic and community analyses because they may affect the biological interpretation of the results. Furthermore, removal of the 6 OTUs containing haplotypes from mixed species did not significantly change the overall community diversity, as we found a strong positive correlation of OTU richness (*R* = 0.91, *p* < 2.2 x 10^−16^) and Sǿrensen dissimilarity (Mantel statistic *R* = 0.98, *p* = 0.001) between datasets with and without the 6 OTUs containing haplotypes from mixed species (supplementary fig. S1). Out of 60 sites, 59 were deemed successful. Site 9 was excluded from downstream analysis due to the limited number of reads that failed to match public and in-house databases.

### Genetic diversity

We considered 59 OTUs for population genetics analysis. These OTUs belong to six orders, namely Coleoptera (4 OTUs), Diptera (17 OTUs), Ephemeroptera (20 OTUs), Megaloptera (1 OTU), Odonata (5 OTUs), Plecoptera (4 OTUs), and Trichoptera (8 OTUs). We found 15 more frequently and 19 less frequently detected OTUs in the catchment (Table 1). Haplotype richness ranged from 1 to 31. GɸST ranged from −6 to 0.637 while average pairwise ɸST ranged from −0.716 to 1(Table 1).

**Table 1.**
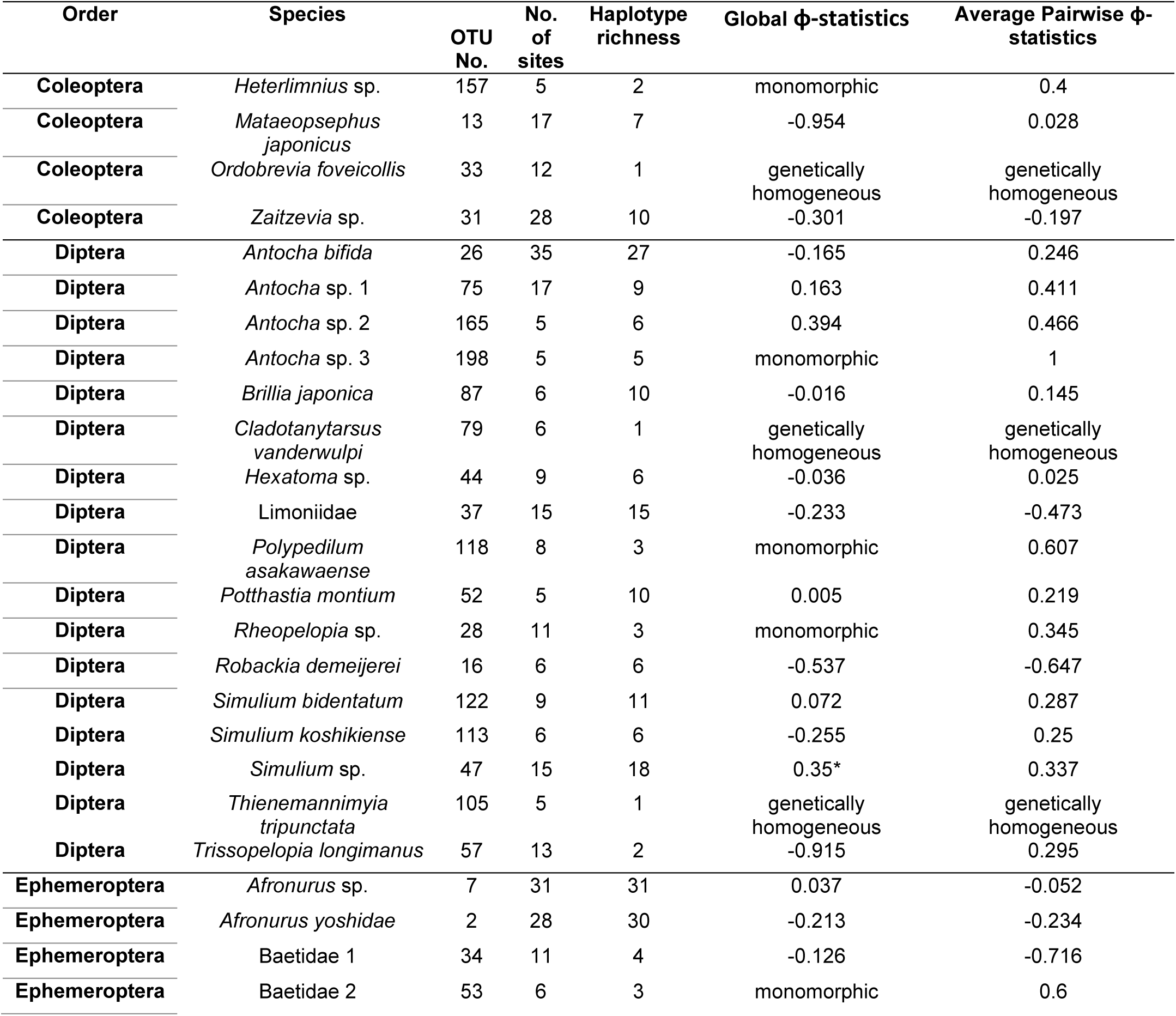
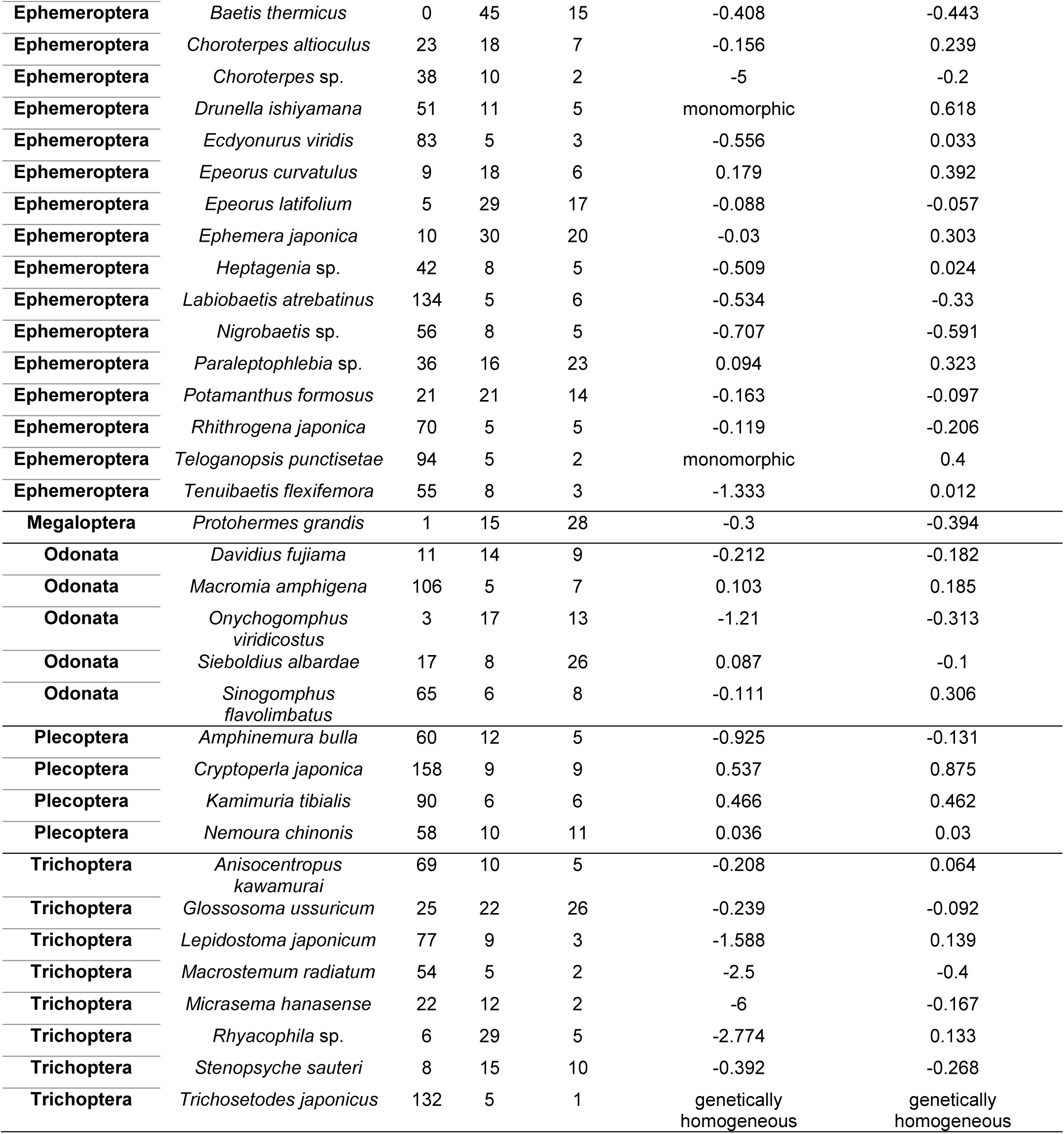
Summary of geographic distribution and genetic diversity of the 59 OTUs used for population genetics analysis.

We found a positive correlation between the number of sites with haplotype richness using 59 OTUs detected in more than 5 sites (*R* = 0.42, *p* = 1 x 10^−5^) and all 194 OTUs (*R* = 0.76, *p* < 2.2 x 10^−16^) (Figure 2, supplementary figure S2). A positive correlation was also observed between the number of sites and average α-genetic diversity using 59 OTUs with more than 5 sites (*R* = 0.22, *p* = 0.018) and 117 OTUs present in at least two sites (*R* = 0.42, *p* = 1.7 x 10^−9^). However, we did not find a correlation between the number of sites and the average pairwise ɸST and GɸST (Fig. 2).

**Figure 2.**
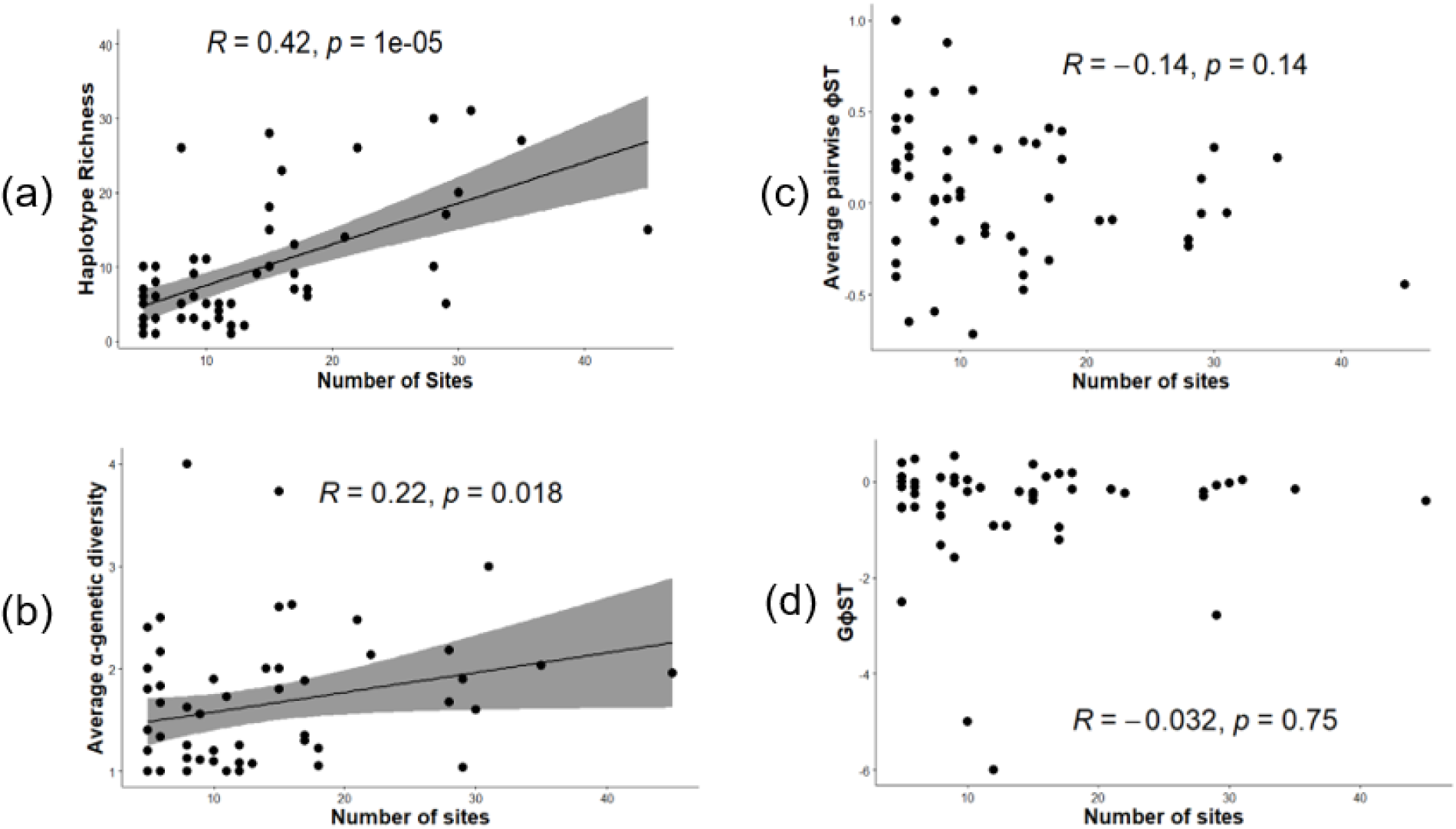
Correlation between number of sites and haplotype richness (a), average α-genetic diversity (b), average pairwise ɸST (c), and GɸST (d)

Haplotype network analysis revealed that 17 OTUs displayed star-like networks, 13 had complex stars, 3 had complex mutational patterns, 1 had reciprocally monophyletic patterns, 4 had genetically homogeneous patterns, and 21 had no pattern (Fig. 3, supplementary fig. S3, and supplementary table S1). In addition to network structures, phylogeographic analysis revealed that 16 OTUs exhibited one high-frequency haplotype in the catchment, 6 OTUs showed 2 high-frequency haplotypes in the catchment, and 14 had an ancestral haplotype that was frequently detected in the catchment (Fig. 3, supplementary fig. S3, and supplementary table S1).

**Figure 3.**
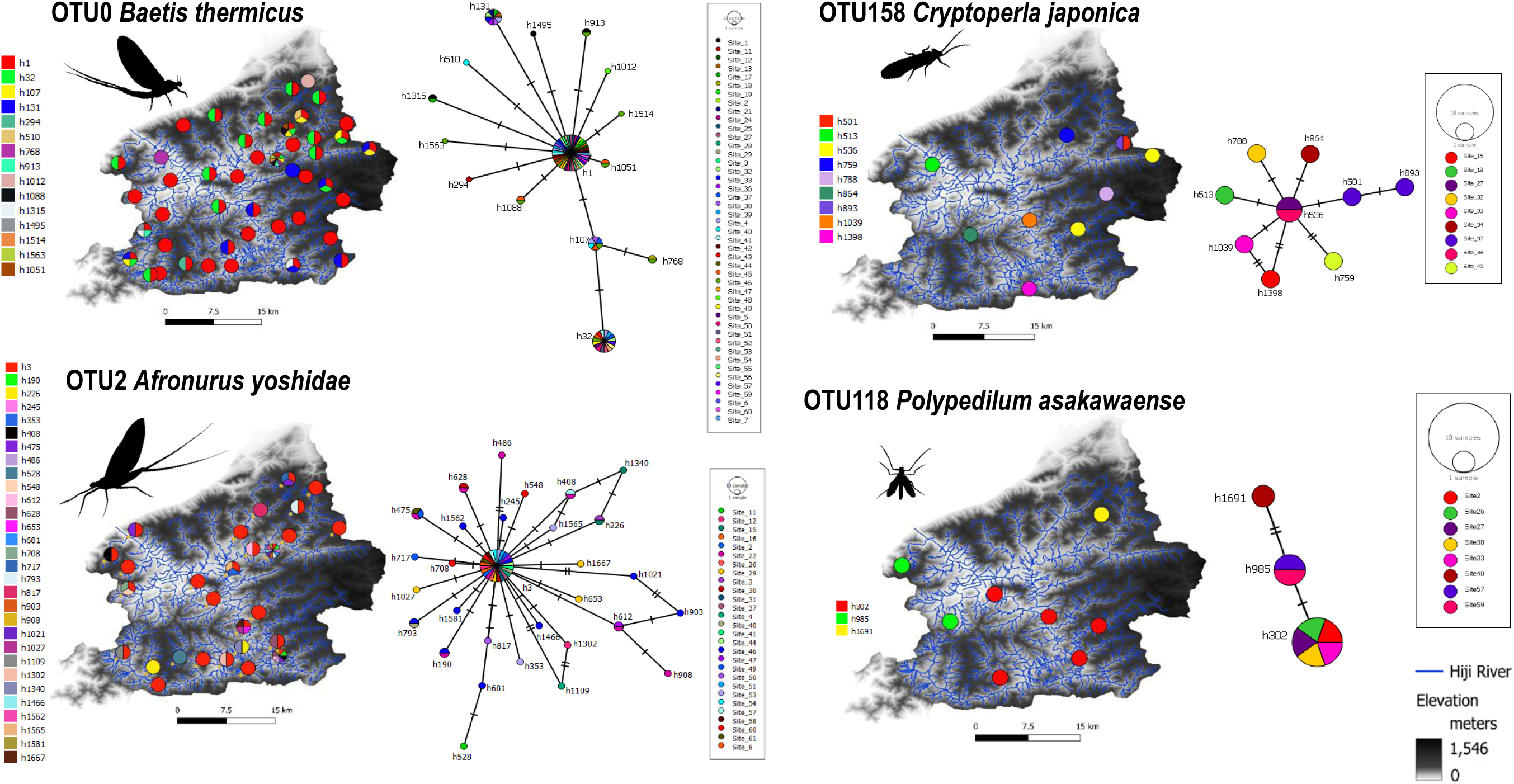
Phylogeography and haplotype network of more frequently detected OTUs (OTU0 *B. thermicus*; OTU2 *A. yoshidae*) and less frequently detected OTUs in the catchment (OTU 158 *C. japonica*; OTU118 *P. asakawaense*)

### Species-Genetic Diversity Correlations

We found two less frequently detected OTUs in the catchment exhibiting a negative α-SGDC, namely, OTU87 (*B. japonica*) (*R* = −0.77, *p* = 0.042) and OTU70 (*R. japonica*) (*R* = −0.89, *p* = 0.037). On the other hand, two moderately detected OTUs in the catchment, namely, OTU34 (Baetidae 1) (*R* = 0.23, *p* = 0.027) and OTU58 (*N. chinonsis*) (*R* = 0.29, *p* = 0.017), demonstrated positive β-SGDC in the catchment. In addition, two more frequently detected OTUs in the catchment also showed this trend: OTU75 (*Antocha* sp. 1) (*R* = 0.17, *p* = 0.045) and OTU6 (*Rhyacophila* sp.) (*R* = 0.26, *p* = 0.001) (Table 2).

**Table 2.** OTUs with statistically significant α and β-SGDCs in the catchment.

| OTU No. | Species | α- SGDC | |
| --- | --- | --- | --- |
|  |  | Kendall R | p-value |
| 87 | *Brillia japonica* | -0.77 | 0.042 |
| 70 | *Rhithrogena japonica* | -0.89 | 0.037 |
| | | $\beta$ - SGDC | |
|  |  | Kendall R | p-value |
| 75 | *Antocha* sp. 1 | 0.17 | 0.045 |
| 34 | Baetidae 1 | 0.23 | 0.027 |
| 58 | *Nemoura chinonsis* | 0.29 | 0.017 |
| 6 | *Rhyacophila* sp. | 0.26 | 0.001 |

### Relationship between female wing length and genetic diversity

We found a positive correlation between the number of sites and the longest female wing length category (*R* = 0.33, *p* = 0.01) (Figure 4). However, we did not found correlation between longest female wing length category with haplotype richness (*R* = 0.23, *p* = 0.071), average pairwise ɸST (*R* = 0.044, *p* = 0.74) and GɸST (*R* = 0.031, *p* = 0.82) (supplementary fig. S4).

**Figure 4.**
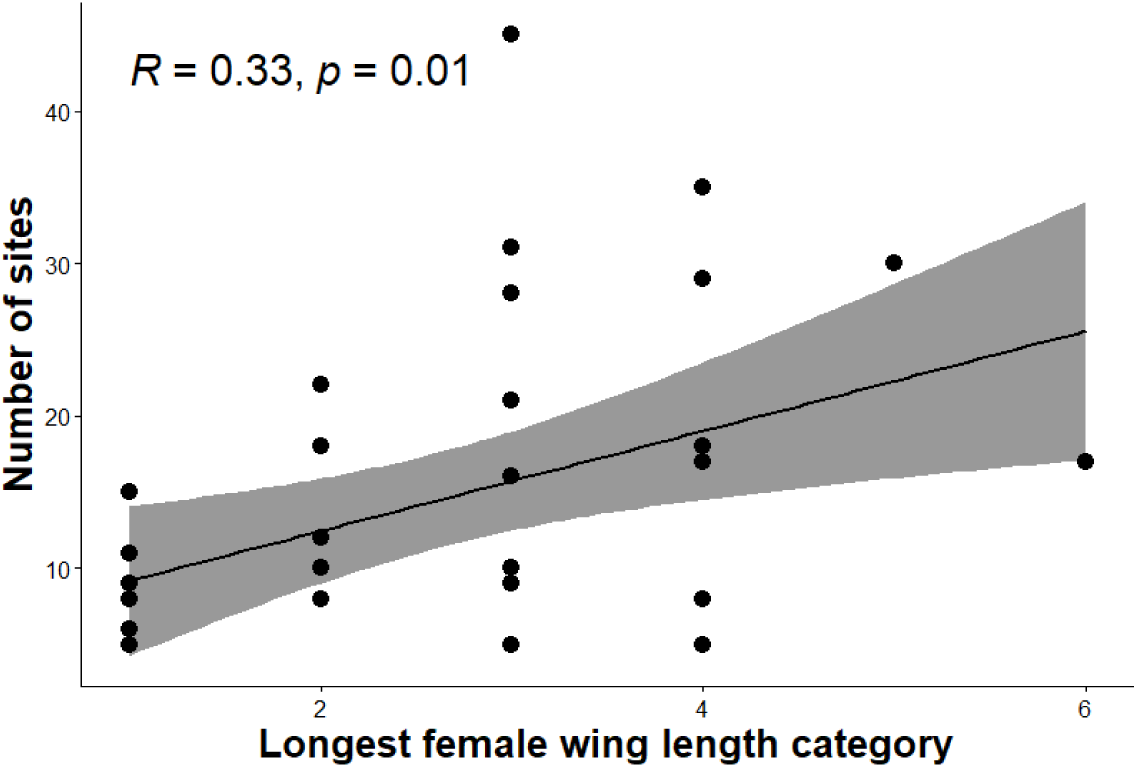
Correlation between number of sites and longest female wing length category of aquatic insects detected in the catchment

### Correlation with Environmental Parameters

Our results showed a positive correlation between the number of sites and the maximum pairwise difference in all environmental parameters (Figure 5a). However, there was no correlation between the number of sites and the mean pairwise difference in these environmental parameters (supplementary fig. S5). Except for NH_4_-N, the maximum pairwise difference in environmental parameters was positively correlated with haplotype richness (Figure 5b). Likewise, except for NH_4_-N, NO_2-_N and elevation, the mean pairwise differences in environmental parameters also showed a positive correlation with haplotype richness (supplementary fig. S5). No correlation was found between either the maximum or mean pairwise difference in environmental parameters and the average pairwise ɸST and GɸST (supplementary fig. S5). However, pH exhibited a positive correlation between the mean pairwise difference in environmental parameters and GɸST (*R* = 0.28, *p* = 0.04985).

**Figure 5.**
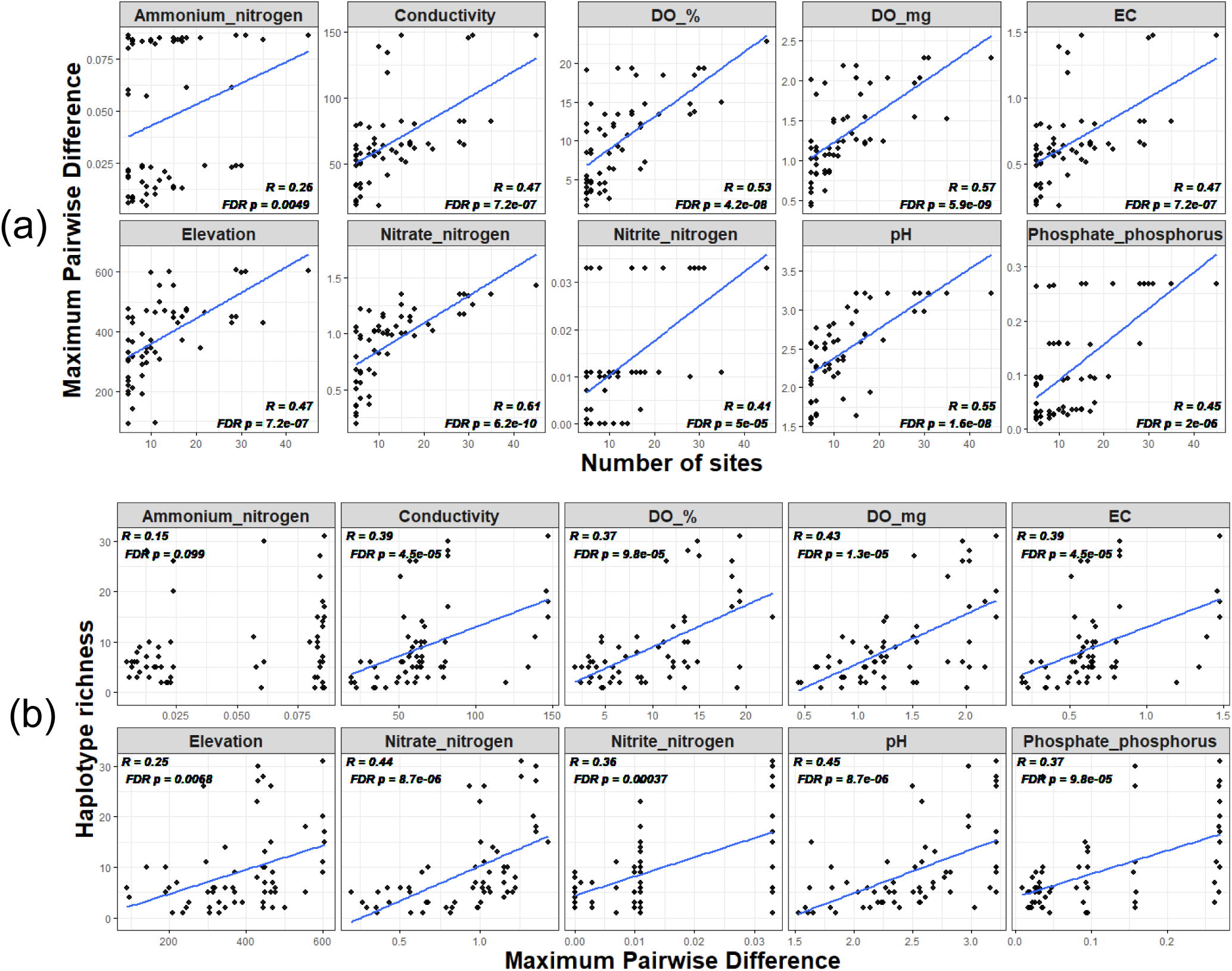
Correlation between the maximum pairwise difference of each environmental parameter with the number of sites (a) and haplotype richness (b) among 59 OTUs

The Mantel test showed that geographic distance significantly influenced the population genetic structure of four moderately detected OTUs and four more frequently detected OTUs within the catchment (supplementary table S2). Community distance decay was observed in five OTUs with moderate detection and five with more frequent detection. However, we did not observe genetic or community distance decay for less frequently detected OTUs, except for OTU165 (*Antocha* sp. 2), which showed a positive correlation between geographic distance and differences in community composition (*R* = 0.60, *p* = 0.025).

Univariate dbRDA indicated that conductivity affected the genetic structure of OTU3 (*Onychogomphus viridicostus*) and OTU60 (*Amphinemura bulla*). PO_4_-P influenced OTU44 (*Hexatoma* sp.)(supplementary table S3). Similarly, ordiR2step showed that conductivity, PO_4_-P, and pH influenced OTU7 (*Afronurus* sp.), OTU26 (*Antocha bifida*), and OTU13 (*Mataeopsephus japonicus*), respectively (supplementary table S4). At the community level, univariate dbRDA showed that elevation explained the most variation in community composition (7 OTUs). NO_3_-N explained variation in 3 OTUs, and NH_4_-N in 1 OTU. This trend was also confirmed by ordiR2step: elevation influenced most community compositions (12/16 OTUs, 75%), followed by NO_3_-N (3/16 OTUs, 18.75%). We did not observe a correlation between genetic structure or community composition and environmental parameters for OTUs less frequently detected in the catchment.

## Discussion

### Relationship between species occurrence and α-, β-, ɣ-genetic diversity

We found a positive correlation between the number of sites and haplotype richness and maximum pairwise difference in all environmental parameters, which supports our hypothesis (1) that species more frequently detected in the catchment exhibit higher ɣ-genetic diversity and a wider range of environmental conditions. A plausible explanation for this pattern is that species more frequently detected in the catchment experienced a wider range of environmental conditions across sites, thereby promoting adaptive divergence among local populations and increasing ɣ-genetic diversity. This pattern may arise from the wider environmental heterogeneity often encompassed by extensive habitat ranges, which drives adaptive divergence among local populations. Previous studies by Bonte & Bafort (2019) and Martin et al. (2021) have similarly shown that spatial environmental heterogeneity enhances ɣ-genetic diversity through adaptive divergence. Our study provides preliminary evidence that the tendency may be more pronounced in species with higher occurrence in the catchment. Future studies should utilize additional genetic markers, preferably non-neutral markers, to achieve a more robust interpretation of adaptive divergence. However, species identification using non-neutral markers may be limited by their limited representation in public databases.

In addition to adaptive divergence, other mechanisms may also explain the observed patterns. As previous studies demonstrated that species abundance is positively correlated with the proportion of sites it occupies within a region (Gaston et al., 2000; Ten Caten et al., 2022), we expect that species more frequently detected might support metapopulations with larger local population sizes at the catchment, diminishing the effects of genetic drift and increasing ɣ-genetic diversity.

However, we did not find a correlation between the number of sites and the average pairwise ɸST and GɸST. We also found that only 2 of 14 OTUs with high average pairwise ɸST (14%) are more frequently detected in the catchment (supplementary table S5). These results did not support our hypothesis (2) that species more frequently detected in the catchment exhibit high β-genetic diversity due to adaptive divergence. One possible explanation is that, assuming species with higher dispersal ability can expand their range within the catchment and occupy more sites, those species occupying more sites may have had the effects of adaptive differentiation described above masked by strong gene flow. Additional support for this explanation comes from our results showing a positive correlation between female wing length and the number of sites, suggesting that more frequently detected species in the catchment have greater dispersal ability. This is consistent with previous studies showing that strong-dispersal aquatic insects exhibit homogeneous populations due to gene flow (Alp et al., 2012; Phillipsen et al., 2015). However, the resolution of our ɸST analysis may be limited for low-frequency haplotypes because kick-net sampling can miss rare or elusive species, and small-bodied, rare species may contribute minimal DNA to the bulk community DNA extract. Such low DNA quantity may fail to amplify during PCR, potentially affecting the interpretation of the ɸST results.

### Species – Genetic Diversity Correlations

Our results suggest that species less frequently detected in the catchment tend to exhibit negative α-SGDCs. This finding did not support our hypothesis (3), which posits that species more frequently detected in the catchment are more likely to exhibit positive α-SGDC. The negative α-SGDCs may be explained by interspecific competition. In highly diverse communities, increased competition among species can reduce each species’ local population size, leading to increased genetic drift, reduced gene flow, and decreased α-genetic diversity. Assuming that species less frequently detected in the catchment are more vulnerable to interspecific competition, this provides a reasonable explanation for the more negative α-SGDCs observed in these species. Conversely, species more frequently detected in the catchment may fail to exhibit the negative α-SGDC due to the weak spatial structuring of haplotypes and species within the catchment, which is mainly driven by the species’ strong dispersal ability. This agrees with Seymour et al. (2016), who found that species and communities with restricted habitat and movement in the river network are more likely to show positive α-SGDC than strong dispersers, due to more spatially structured populations and communities and stronger local selection pressures.

We observed positive β-SGDC in two moderately detected OTUs and two more frequently detected OTUs in the catchment. However, we did not find β-SGDC in the less frequently detected OTUs in the catchment. This partially supports our hypothesis (3), which states that more frequently detected OTUs in the catchment are more likely to exhibit positive β-SGDC. The observed β-SGDC may be attributed to parallel divergences in communities and populations resulting from geographic isolation. Our analysis of the correlation between geographic distance, pairwise ɸST and Sørensen dissimilarity may support this interpretation, as we observed both genetic and community distance decay in moderately and more frequently detected OTUs but not in less frequently detected OTUs (supplementary table S2). Drift-migration equilibrium may lead to community distance decay (Nekola and White 1999; Soininen et al. 2007; Cañedo-Arguelles 2015) and genetic isolation by distance (IBD) among populations (Wright 1943; Phillipsen et al. 2015), resulting in β-SGDC (Moritz 2002), especially for species with moderate dispersal abilities (Cañedo-Arguelles 2015; Phillipsen et al. 2015). Species with broad habitat ranges, particularly those with moderate dispersal abilities, are likely to experience significant reductions in connectivity between distant sites. Thus, the detectability of the β-SGDC may benefit from these parallel influences on communities and populations driven by geographic distance and species’ dispersal capability.

Spatially varying selection among heterogeneous environments between sites may also cause adaptive divergence at the community level (Webb et al. 2010) or the population level (Kawecki and Ebert 2004). If common environmental factors drive adaptive divergence at both levels, such divergence is expected to correlate, thereby contributing to β-SGDC (Kawecki and Ebert 2004; Vellend 2003). However, our findings do not support the notion that divergences in communities and populations are driven by common environmental factors. Our overall analysis indicated that water chemistry (conductivity, PO_4_P, pH) primarily influences genetic differences among local populations, whereas elevation plays a more significant role in shaping differences among local communities (supplementary tables S3 and S4). These contrasting findings challenge our expectation of a common environmental factor influencing β-SGDC. Although this aligns with Martin et al. (2021), who found that metapopulations and metacommunities are influenced by distinct spatial and environmental factors, further research is needed to clarify this relationship.

### Haplotype-level metabarcoding limitations and future perspectives

Our results demonstrate the effectiveness of haplotype-level metabarcoding for analyzing multiple parallel populations. This approach enabled us to compare genetic diversity, population genetic structure, and SGDC across species inhabiting the same watershed. This interspecific comparison provides evidence that species-specific evolutionary traits are strongly associated with the extent of each species’ distribution, enabling the testing of evolutionary and ecological hypotheses that were previously difficult to evaluate using population genetic analyses of a single or few species.

However, there are limitations to haplotype-level metabarcoding that warrant further improvements in future research. First, there are concerns about potential biases arising from reliance on the number of sequence reads for estimating allele frequencies. This bias can be significantly influenced by PCR amplification, next-generation sequencing depth, and specimen size. In this study, we transformed haplotype sequence reads into presence/absence data to remove bias, and the only source of variation is mutational differences among haplotypes (Azarian et al., 2020). However, to increase the likelihood of obtaining statistically significant population differentiation, future studies should consider using replicate samples (Azarian et al., 2020). Additionally, we recommend that future studies explore whether approaches such as quantitative MiSeq (qMiSeq) sequencing (Ushio et al. 2018), which can estimate the relative abundance of each species in species-level metabarcoding, can also be adapted to estimate allele frequencies at the haplotype level. In qMiSeq, by incorporating a known quantity of short DNA fragments (internal standard DNA) from haplotypes of species absent from the study area into all samples during PCR, the number of reads from non-standard DNA may be adjusted subsequently.

In future studies, environmental DNA (eDNA) may also be used for eDNA haplotyping. This approach allows less invasive, time-saving, and cost-effective sampling. It enables the collection of more samples and a broader comparison of intraspecific variation across taxa and regions. As a result, the impact of false negatives in rare and small species can be minimized (Elbrecht et al. 2018). However, reliable data require careful attention to the challenges of environmental samples, including eDNA degradation and PCR inhibition.

## Conclusions

In summary, we investigated relationships among species occurrence, population genetic structure, and SGDC using haplotype-level bulk DNA metabarcoding. This work expands our understanding of the applications of DNA metabarcoding in biodiversity studies. We found contrasting population genetic structure and SGDC between species with higher and lower occurrence in the catchment, likely driven by dispersal ability and environmental heterogeneity. These findings help ongoing efforts to reduce the impacts of rapid environmental change on biodiversity, influenced by human activities and climate change.

## Supporting information

supplementary protocol 1

## Acknowledgements

We extend our gratitude to Dr. Naohito Tokunaga of the Division of Analytical BioMedicine, Advanced Research Support Center, Ehime University, for his assistance in performing the Illumina Miseq Sequencing.

## Author Contributions

Conceptualization: D.J.L., K.W., M-C.C. Data curation, Investigation, Methodology: D.J.L., M-C.C., A.D-L., L-P.K., J.M.S. Formal analysis: D.J.L., K.W., K.H. Writing–original draft: D.J.L. Writing–review and editing: D.J.L., K.W., J.M.S., K.H.

## Data Statement

All demultiplexed sequences and their corresponding metadata as part of this study were available through NCBI SRA under the Bioproject number PRJNA1191972.

## References

Alp M, Keller I, Westram AM, Robinson CT (2012). How river structure and biological traits influence gene flow: a population genetic study of two stream invertebrates with differing dispersal abilities. Freshwater Biology, 57(5): 969–981.10.1111/j.1365-2427.2012.02758.x

Ameijeiras-Alonso J, Crujeiras RM, Rodríguez-Casal A (2019). “Mode Testing, Critical Band width and Excess Mass.” Test, 28(3), 900–919. doi:10.1007/s11749-018-0611-5.

Ameijeiras-Alonso J, Crujeiras RM, Rodríguez-Casal A (2021). multimode: Mode Testing and Exploring. R package version 1.5, URL https://CRAN.R-project.org/package=multimode.

Andrews S (2010) FastQC: A quality control tool for high throughput sequence data.

Arulandhu AJ, Staats M, Hagelaar R, Voorhuijzen MM, Prins TW, Scholtens I, … & Kok E (2017). Development and validation of a multi-locus DNA metabarcoding method to identify endangered species in complex samples. Gigascience, 6(10), gix080. 10.1093/gigascience/gix080

Azarian C, Foster S, Devloo - Delva F, Feutry P (2021) Population differentiation from environmental DNA: Investigating the potential of haplotype presence/absence - based analysis of molecular variance. Environmental DNA, 3(3), 541–552. 10.1002/edn3.143

Bandelt H, Forster P, Röhl A (1999) Median-joining networks for inferring intraspecific phylogenies. Molecular Biology Evolution 16(1):37–48. 10.1093/oxfordjournals.molbev.a026036

Berry KJ, Johnston JE, Mielke Jr PW (2011). Permutation methods. Wiley Interdisciplinary Reviews: Computational Statistics, 3(6): 527–542. 10.1002/wics.177

Bonte D, Bafort Q (2019) The importance and adaptive value of life - history evolution for metapopulation dynamics. Journal of Animal Ecology 88(1):24–34. 10.1111/1365-2656.12928

Callahan BJ, McMurdie PJ, Rosen MJ, Han AW, Johnson AJA, Holmes SP (2016) DADA2: High-resolution sample inference from Illumina amplicon data. Nature Methods 13(7):581–583. 10.1038/nmeth.3869

Cañedo - Argüelles M, Boersma KS, Bogan MT, Olden JD, Phillipsen I, Schriever TA, Lytle DA (2015) Dispersal strength determines meta - community structure in a dendritic riverine network. Journal of Biogeography 42(4):778–790. 10.1111/jbi.12457

Darlington Jr. PJ (1943). Carabidae of mountains and islands: data on the evolution of isolated faunas, and on atrophy of wings. Ecological Monographs, 13(1):37–61.10.2307/1943589

Dijkstra KDB, Monaghan MT, Pauls SU (2014) Freshwater biodiversity and aquatic insect diversification. The Annual Review of Entomology 59(1):143–163. 10.1146/annurev-ento-011613-161958

Elbrecht V, Vamos EE, Steinke D, Leese F (2018) Estimating intraspecific genetic diversity from community DNA metabarcoding data. PeerJ 6:e4644. 10.7717/peerj.4644

Elbrecht V, Leese F (2017) Validation and development of freshwater invertebrate metabarcoding COI primers for environmental impact assessment. Frontiers in Freshwater Science 5(11):1–11. 10.3389/fenvs.2017.00011

Elbrecht V, Steinke D (2019) Scaling up DNA metabarcoding for freshwater macrozoobenthos monitoring. Freshwater Biology 64(2):380–387. 10.1111/fwb.13220

Excoffier L, Smouse P, Quattro J (1992) Analysis of molecular variance inferred from metric distances among DNA haplotypes: Application to human mitochondrial DNA restriction data. Genetics 131:479–491.10.1093/genetics/131.2.479

Excoffier L, Lischer HE (2010) Arlequin suite ver 3.5: A new series of programs to perform population genetics analyses under Linux and Windows. Molecular Ecology Resources 10:564–567.10.1111/j.1755-0998.2010.02847.x

Fourcade Y (2016). Comparing species distributions modelled from occurrence data and from expert-based range maps. Implication for predicting range shifts with climate change. Ecological Informatics, 36, 8–14. 10.1016/j.ecoinf.2016.09.002

Fourtune L, Paz - Vinas I, Loot G, Prunier JG, Blanchet S (2016) Lessons from the fish: A multi - species analysis reveals common processes underlying similar species - genetic diversity correlations. Freshwater Biology 61(11):1830–1845.10.1111/fwb.12826.

Gaston KJ, Blackburn TM, Greenwood JJ, Gregory RD, Quinn RM, Lawton JH (2000). Abundance–occupancy relationships. Journal of Applied Ecology, 37, 39–59. 10.1046/j.1365-2664.2000.00485.x

Gaston H, He F (2011) Species occurrence and occupancy. In: Magurran AE, McGill BJ (Eds) Biological Diversity: Frontiers in Measurement and Assessment. Oxford University Press, 141–151.

Gaston KJ, Kunin WE (1997) Rare-common differences: an overview. In: Gaston KJ, Kunin WE (Eds) The Biology of Rarity. Chapman & Hall, London, 12–13.

Gentile G, Argano R, Taiti S (2022). Evaluating the correlation between area, environmental heterogeneity, and species richness using terrestrial isopods (Oniscidea) from the Pontine Islands (West Mediterranean). Organisms Diversity & Evolution, 22(1):275–284.10.1007/s13127-021-00523-x

Gitzendanner MA, Soltis PS (2000). Patterns of genetic variation in rare and widespread plant congeners. American journal of botany, 87(6):783–792. 10.2307/2656886

Gotelli NJ, Stanton - Geddes J (2015). Climate change, genetic markers and species distribution modelling. Journal of Biogeography, 42(9): 1577–1585. 10.1111/jbi.12562

Guillera - Arroita G (2017). Modelling of species distributions, range dynamics and communities under imperfect detection: advances, challenges and opportunities. Ecography, 40(2): 281–295. doi: 10.1111/ecog.02445

Holt RD (2003) On the evolutionary ecology of species’ ranges. Evolutionary Ecology Research 5(2):159–178.

Jenkins TL, Castilho R, Stevens JR (2018) Meta-analysis of northeast Atlantic marine taxa shows contrasting phylogeographic patterns following post-LGM expansions. PeerJ 6:e5684.10.7717/peerj.5684

Kahilainen A, Puurtinen M, Kotiaho JS (2014) Conservation implications of species–genetic diversity correlations. Global Ecology and Conservation 2:315–323. 10.1016/j.gecco.2014.10.013

Kassambara A (2023) ggpubr: ‘ggplot2’ Based Publication Ready Plots. R package version 0.6.0. https://CRAN.R-project.org/package=ggpubr

Kawecki T, Ebert D (2004) Conceptual issues in local adaptation. Ecology Letters 7(12):1225–1241. 10.1111/j.1461-0248.2004.00684.x

Loveless, M. D., & Hamrick, J. L. (1984). Ecological determinants of genetic structure in plant populations. Annual review of ecology and systematics, 15:65–95.

Martin M (2011) Cutadapt removes adapter sequences from high-throughput sequencing reads. EMBnet.Journal 17(1):10–12. 10.14806/ej.17.1.200

Martin GK, Beisner BE, Chain FJ, Cristescu ME, Del Giorgio PA, Derry AM (2021) Freshwater zooplankton metapopulations and metacommunities respond differently to environmental and spatial variation. Ecology 102(1):e03224. 10.1002/ecy.3224

Moritz C (2002) Strategies to protect biological diversity and the evolutionary processes that sustain it. Systematic Biology 51:238–254. 10.1080/10635150252899752

Nekola JC, White PS (1999) The distance decay of similarity in biogeography and ecology. Journal of Biogeography 26:867–878. 10.1046/j.1365-2699.1999.00305.x

Oksanen J, Simpson G, Blanchet F, Kindt R, Legendre P, Minchin P, O’Hara R, Solymos P, Stevens M, Szoecs E, et al. (2022) vegan: Community Ecology Package. R package version 2.6–4. https://CRAN.R-project.org/package=vegan

Phillipsen IC, Kirk EH, Bogan MT, Mims MC, Olden JD, Lytle DA (2015) Dispersal ability and habitat requirements determine landscape - level genetic patterns in desert aquatic insects. Molecular Ecology 24(1):54–69. 10.1111/mec.13003

Pierce AA, Gutierrez R, Rice AM, Pfennig KS (2017) Genetic variation during range expansion: effects of habitat novelty and hybridization. Proceedings of the Royal Society B 284(1852):20170007. 10.1098/rspb.2017.0007

Polechová J (2018) Is the sky the limit? On the expansion threshold of a species’ range. PLoS Biology 16(6): e2005372. 10.1371/journal.pbio.2005372

Porter TM, Hajibabaei M (2021) Profile hidden Markov model sequence analysis can help remove putative pseudogenes from DNA barcoding and metabarcoding datasets. BMC Bioinformatics 22(1):256. 10.1186/s12859-021-04180-x

Ranwez V, Harispe S, Delsuc F, Douzery EJP (2011) MACSE: Multiple Alignment of Coding SEquences accounting for frameshifts and stop codons. PLoS One 6(9):e22594, 10.1371/journal.pone.0022594

Ranwez V, Douzery EJP, Cambon C, Chantret N, Delsuc F (2020) MACSE v2: Toolkit for the Alignment of Coding Sequences Accounting for Frameshifts and Stop Codons. Molecular Biology and Evolution 35(10): 2582–2584. 10.1093/molbev/msy159

Rognes T, Flouri T, Nichols B, Quince C, Mahé F (2016) VSEARCH: a versatile open source tool for metagenomics. PeerJ 4:e2584. 10.7717/peerj.2584

Sarremejane R, Cid N, Stubbington R, Datry T, Alp M, Cañedo-Argüelles M, & Bonada, N (2020) DISPERSE, a trait database to assess the dispersal potential of European aquatic macroinvertebrates. Scientific data, 7(1): 386. 10.1038/s41597-020-00732-7

Serrana JM, Watanabe K (2023) Haplotype-level metabarcoding of freshwater macroinvertebrate species: a prospective tool for population genetic analysis. PLoS One 18(7): e0289056. 10.1371/journal.pone.0289056

Seymour M, Seppälä K, Mächler E, Altermatt F (2016) Lessons from the macroinvertebrates: species - genetic diversity correlations highlight important dissimilar relationships. Freshwater Biology 61(11):1819–1829. 10.1111/fwb.12816

Soininen J, McDonald R, Hillebrand H (2007) The distance decay of similarity in ecological communities. Ecography 30:3–12. 10.1111/j.0906-7590.2007.04817.x

Starr SM, Wallace JR (2021) Ecology and Biology of Aquatic Insects. Insects 12(1):51. 10.3390/insects12010051

Ten Caten C, Holian L, Dallas T (2022) Weak but consistent abundance–occupancy relationships across taxa, space and time. Global Ecology and Biogeography, 31(5), 968–977. 10.1111/geb.13472

Tingley MW, Beissinger SR (2009). Detecting range shifts from historical species occurrences: new perspectives on old data. Trends in ecology & evolution, 24(11): 625–633.

Turon X, Antich A, Palacín C, Præbel K, Wangensteen OS (2020) From metabarcoding to metaphylogeography: separating the wheat from the chaff. Ecological Applications 30(2): e02036. 10.1002/eap.2036

Ushio M, Murakami H, Masuda R, Sado T, Miya M, Sakurai S, Yamanaka H, Minamoto T, Kondoh M (2018) Quantitative monitoring of multispecies fish environmental DNA using high-throughput sequencing. Metabarcoding Metagenomics 2: e23297. 10.3897/mbmg.2.23297

Vellend M (2003) Island biogeography of genes and species. The American Naturalist 162:358–365.10.1086/377189

Vellend M, Geber MA (2005) Connections between species diversity and genetic diversity. Ecology Letters 8(7):767–781. doi: 10.1111/j.1461-0248.2005.00775.x

Watanabe K, Monaghan MT (2017). Comparative tests of the species-genetic diversity correlation at neutral and nonneutral loci in four species of stream insect. Evolution, 71(7): 1755–1764.10.1111/evo.13261

Webb CT, Hoeting JA, Ames GM, Pyne MI, Poff NL (2010) A structured and dynamic framework to advance traits-based theory and prediction in ecology. Ecology Letters 13:267–283. 10.1111/j.1461-0248.2010.01444.x

Weir BS, Cockerham CC (1984) Estimating F-statistics for the analysis of population structure. Evolution 38:1358–1370.10.1111/j.1558-5646.1984.tb05657.x

Weir BS (1996) Genetic Data Analysis II: Methods for Discrete Population Genetic Data Sinauer Assoc., Inc., Sunderland, MA, USA.

Wickham H (2016) ggplot2: Elegant Graphics for Data Analysis. Springer-Verlag New York.

Wright S (1943) Isolation by distance. Genetics 28:114–138.10.1093/genetics/28.2.114

Zhang J, Kapli P, Pavlidis P, Stamatakis A (2013) A general species delimitation method with applications to phylogenetic placements. Bioinformatics 29:2869–2876. 10.1093/bioinformatics/btt499

