## supplementary fig. S3 for "Haplotype-level metabarcoding revealed the relationship between species occurrence and population genetic structure at the catchment scale in aquatic insects"

**Supplementary Figure S3.** Phylogeography and haplotype network for 59 OTUs used for population genetic analysis

### OTU 0

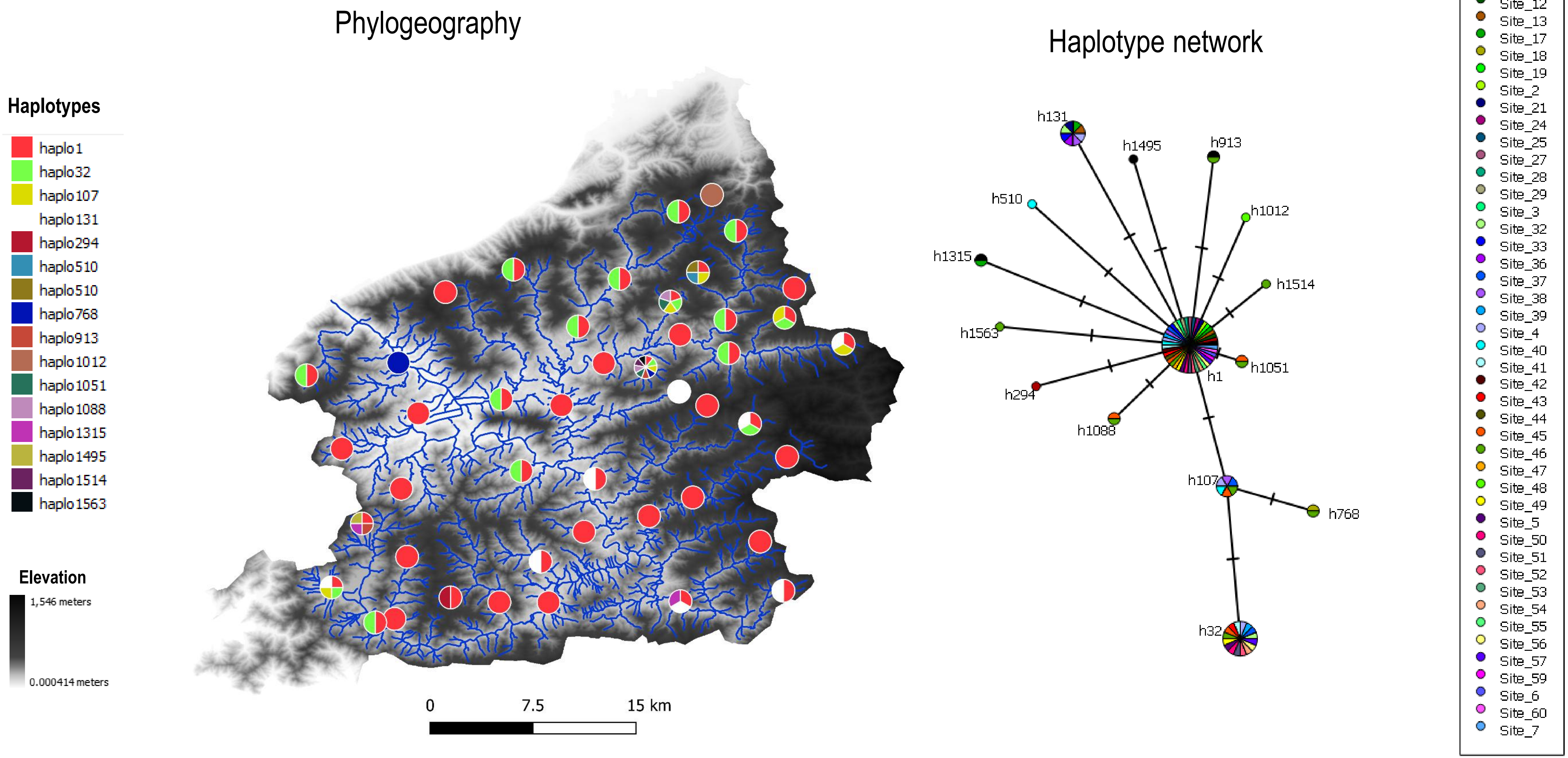
Phylogeography
Haplotype network
Haplotypes
haplo1
haplo32
haplo107
haplo131
haplo294
haplo510
haplo510
haplo768
haplo913
haplo1012
haplo1051
haplo1088
haplo1315
haplo1495
haplo1514
haplo1563
Elevation
1,546 meters
0.000414 meters
0
7.5
15 km
h131
h1495
h913
h510
h1012
h1315
h1514
h1563
h1051
h1
h294
h1088
h107
h768
h32
10 samples
1 sample
Site_1
Site_11
Site_12
Site_13
Site_17
Site_18
Site_19
Site_2
Site_21
Site_24
Site_25
Site_27
Site_28
Site_29
Site_3
Site_32
Site_33
Site_36
Site_37
Site_38
Site_39
Site_4
Site_40
Site_41
Site_42
Site_43
Site_44
Site_45
Site_46
Site_47
Site_48
Site_49
Site_5
Site_50
Site_51
Site_52
Site_53
Site_54
Site_55
Site_56
Site_57
Site_59
Site_6
Site_60
Site_7

### OTU 26

#### Phylogeography

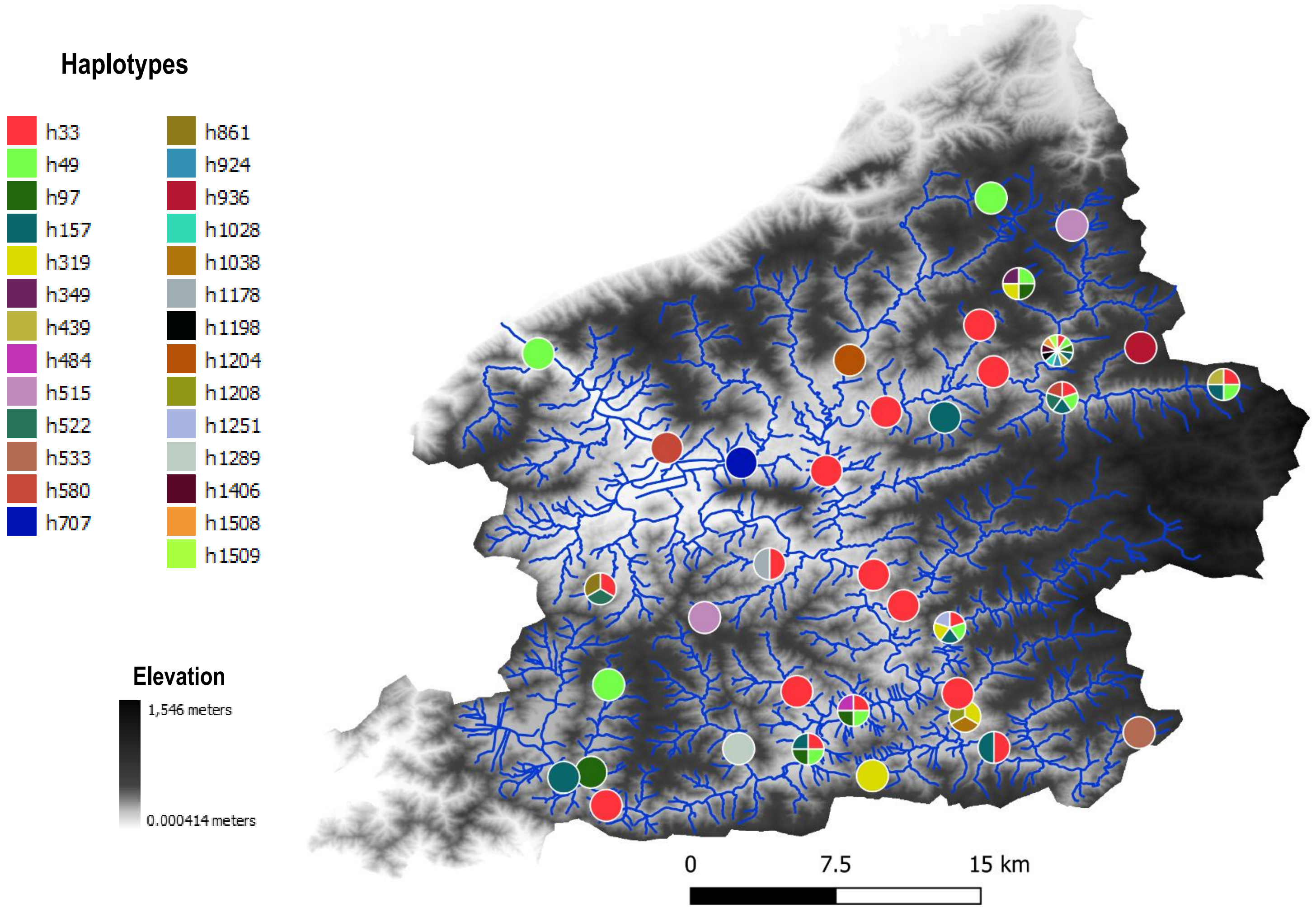


#### Haplotype network

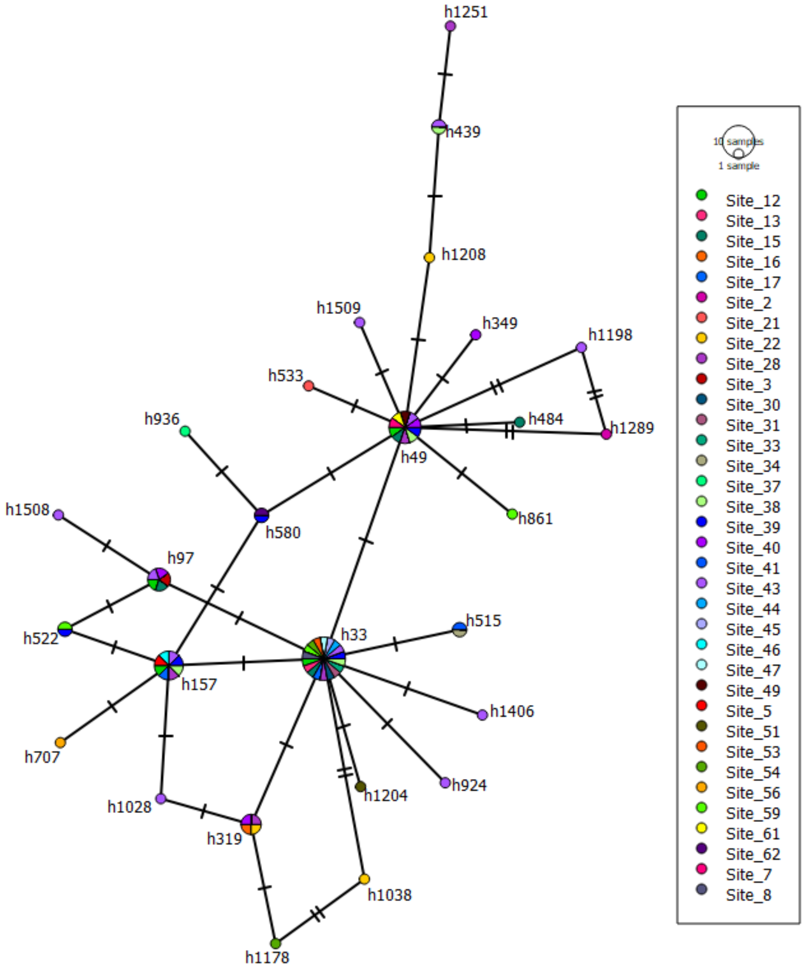

### OTU 7

#### Phylogeography

Haplotypes

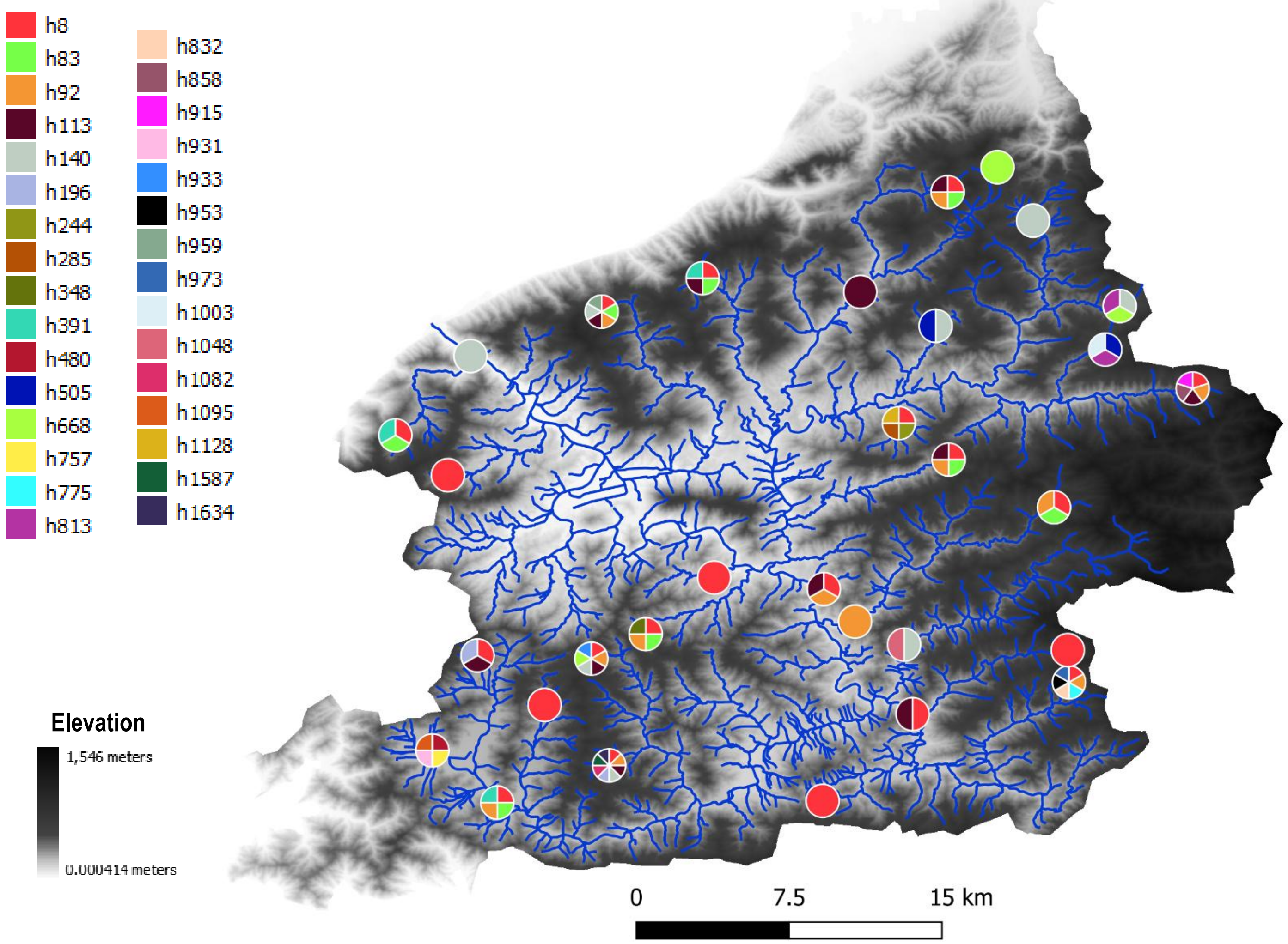


#### Haplotype network

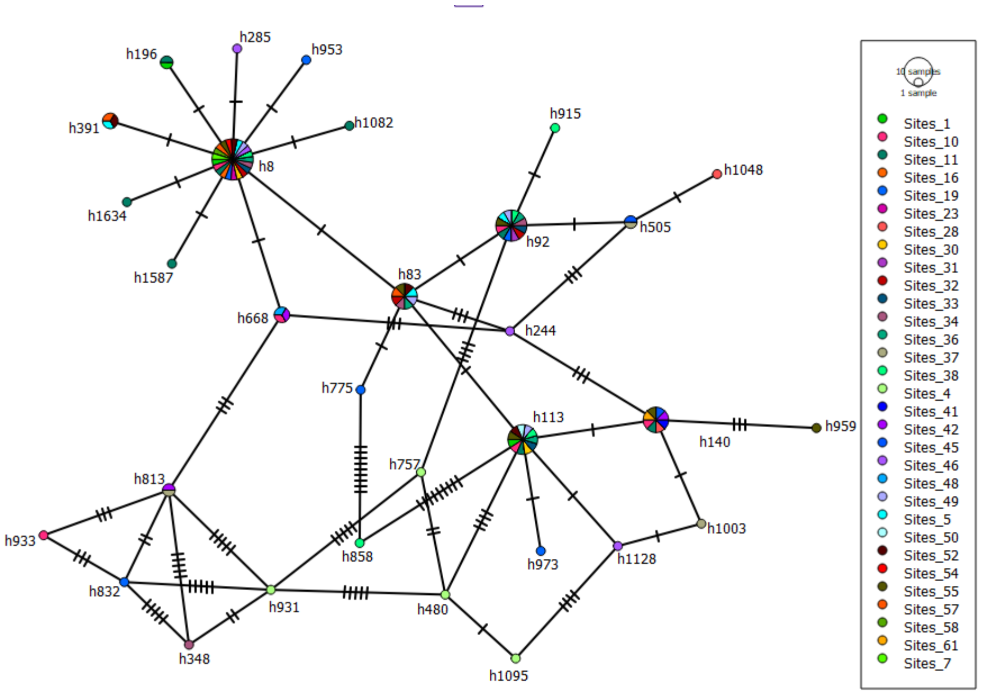

### OTU 10

#### Phylogeography

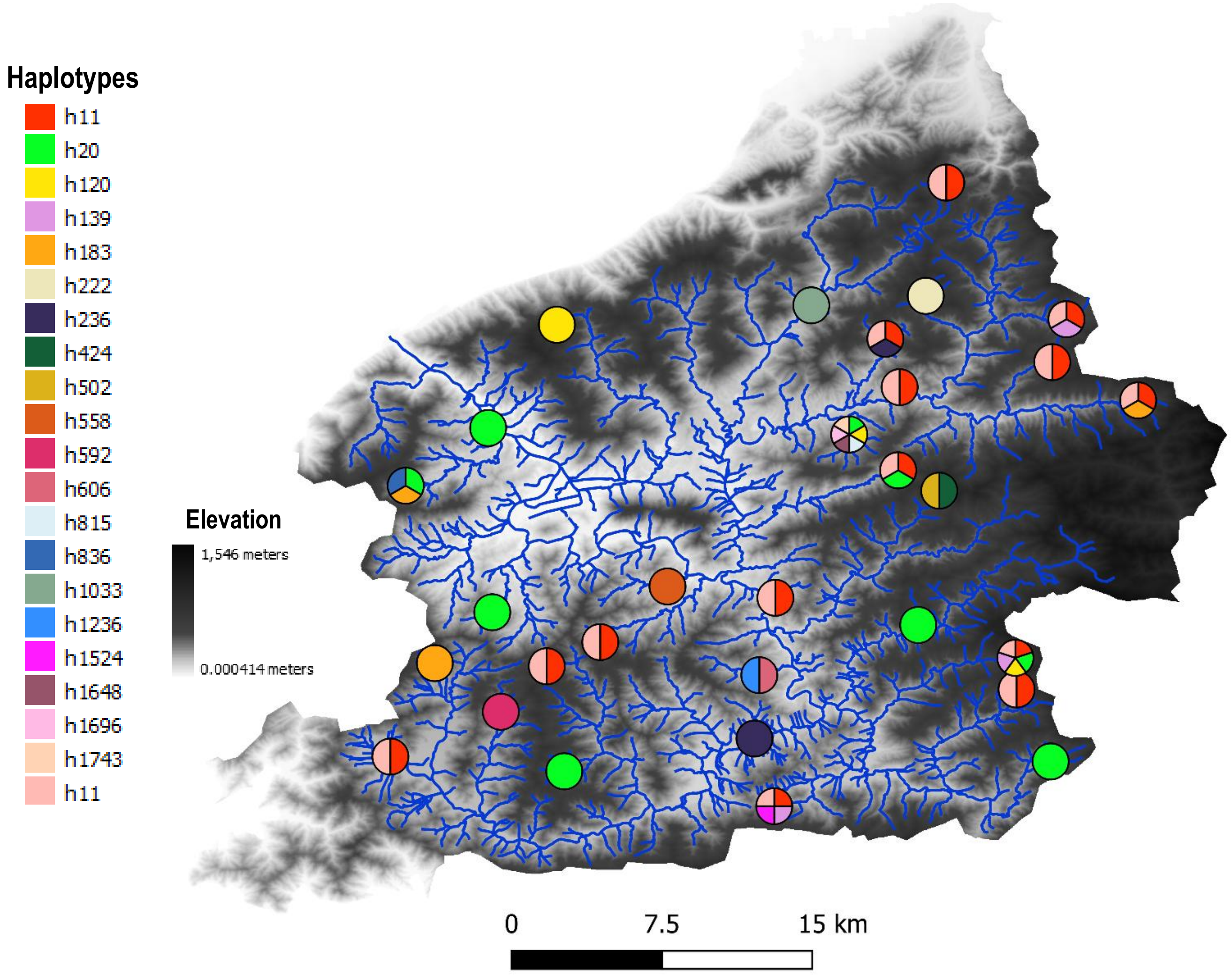


#### Haplotype network

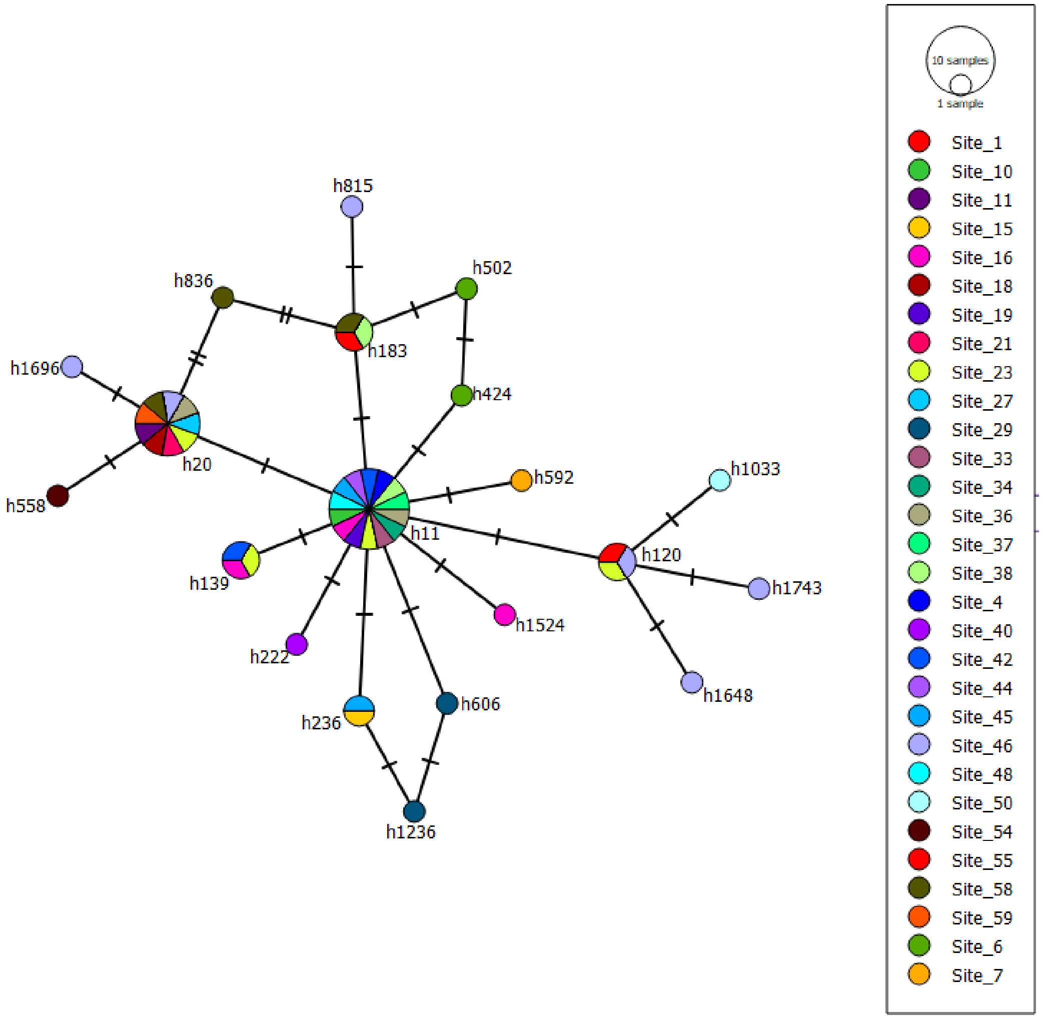

### OTU 5

#### Phylogeography

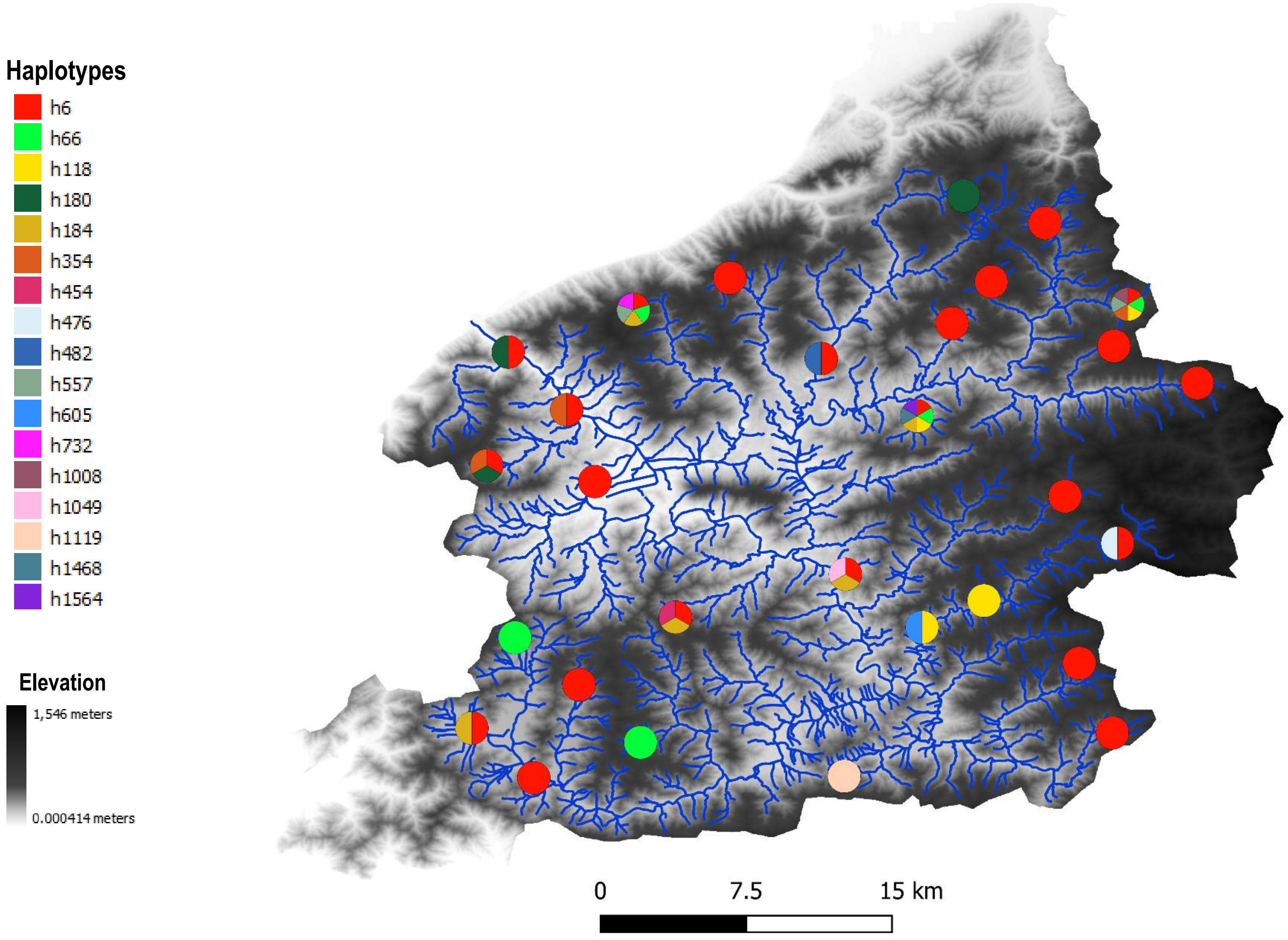


#### Haplotype network

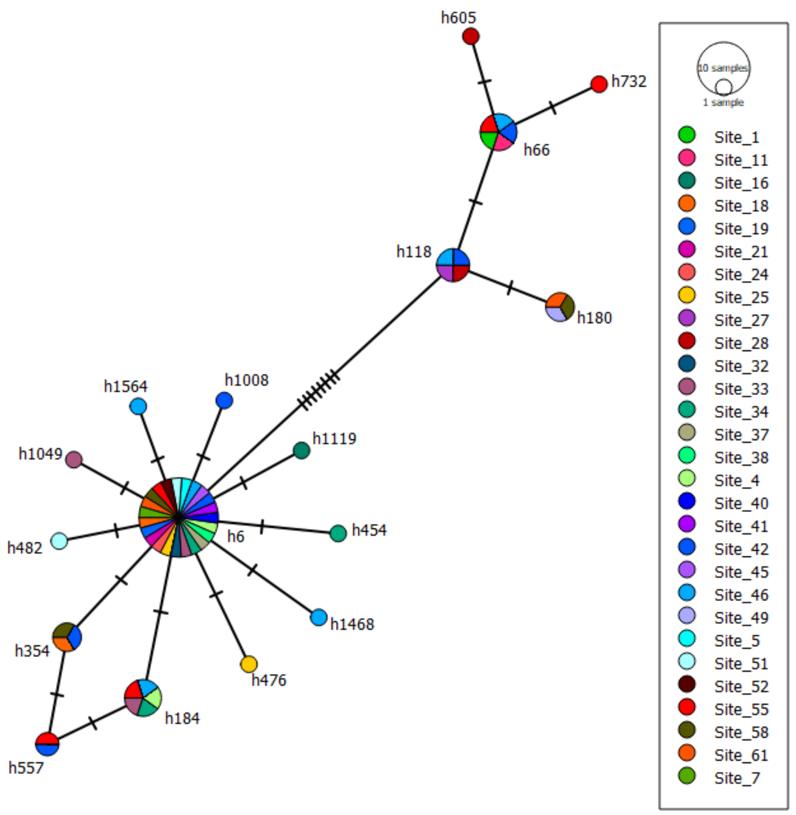

### OTU 6

#### Phylogeography

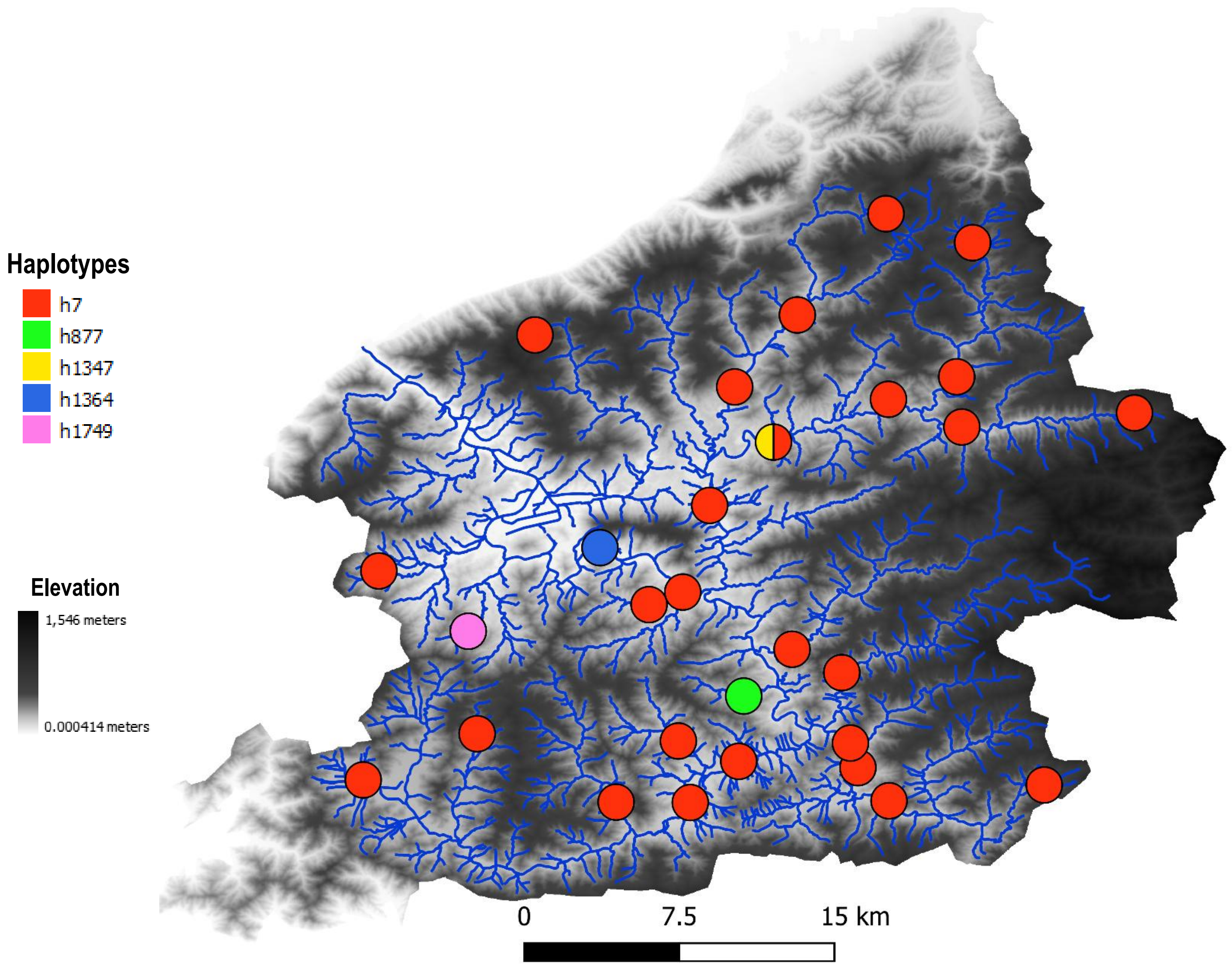


#### Haplotype network

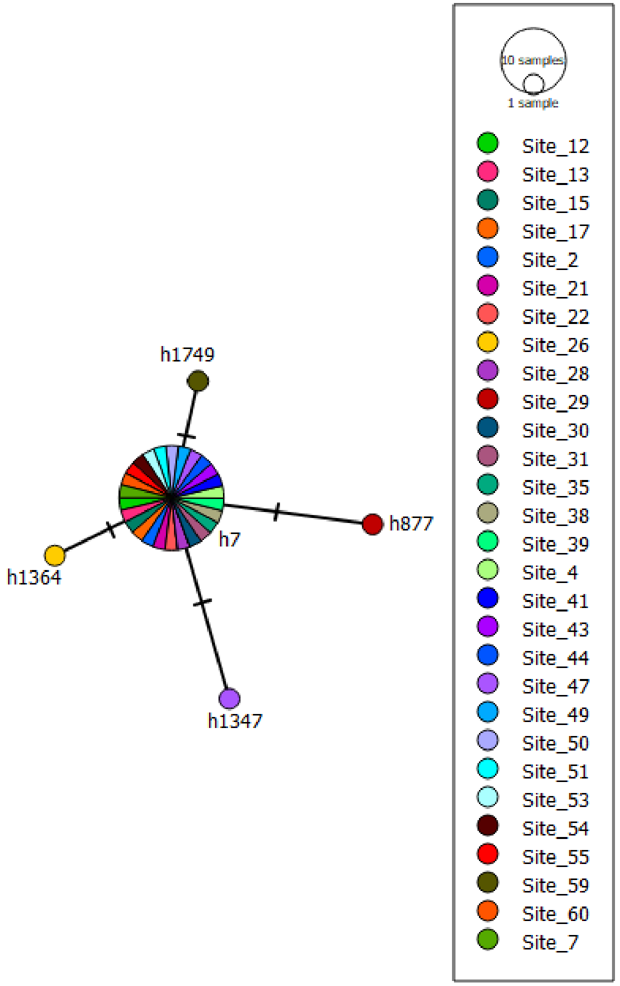

### OTU 2

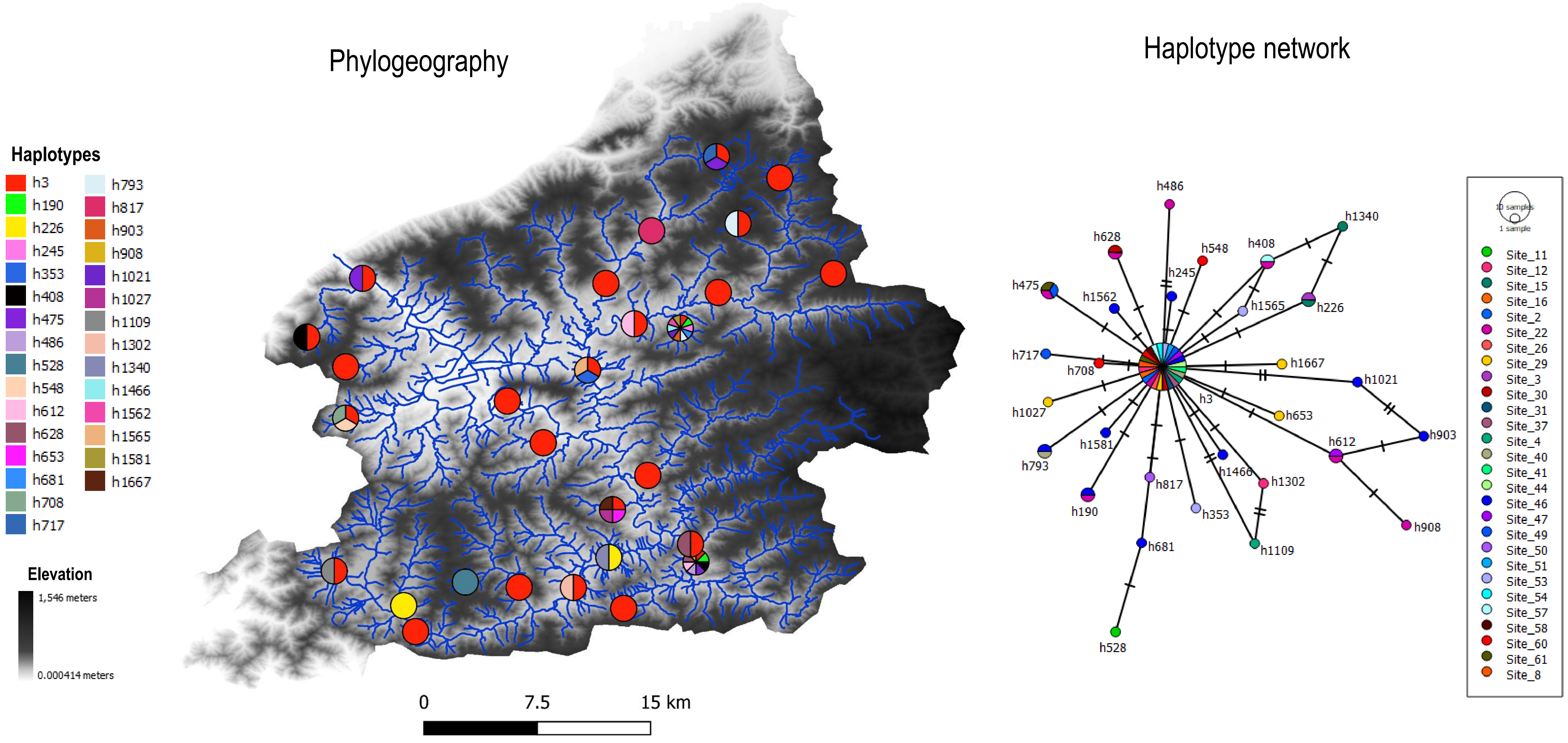
Phylogeography
Haplotype network
Haplotypes
h3
h190
h226
h245
h353
h408
h475
h486
h528
h548
h612
h628
h653
h681
h708
h717
h793
h817
h903
h908
h1021
h1027
h1109
h1302
h1340
h1466
h1562
h1565
h1581
h1667
Elevation
1,546 meters
0.000414 meters
0
7.5
15 km
10 samples
1 sample
Site_11
Site_12
Site_15
Site_16
Site_2
Site_22
Site_26
Site_29
Site_3
Site_30
Site_31
Site_37
Site_4
Site_40
Site_41
Site_44
Site_46
Site_47
Site_49
Site_50
Site_51
Site_53
Site_54
Site_57
Site_58
Site_60
Site_61
Site_8

### OTU 31

#### Phylogeography

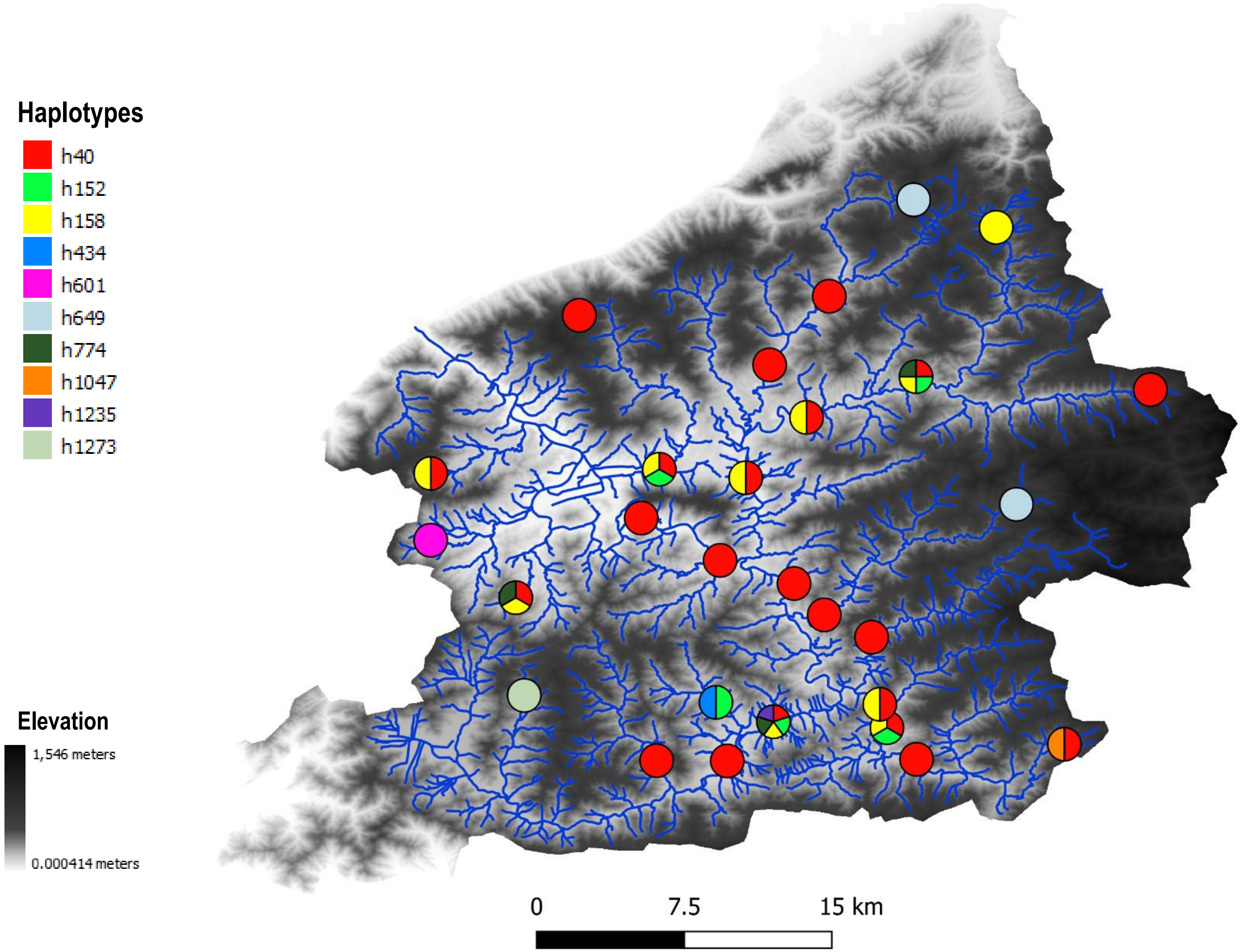


#### Haplotype network

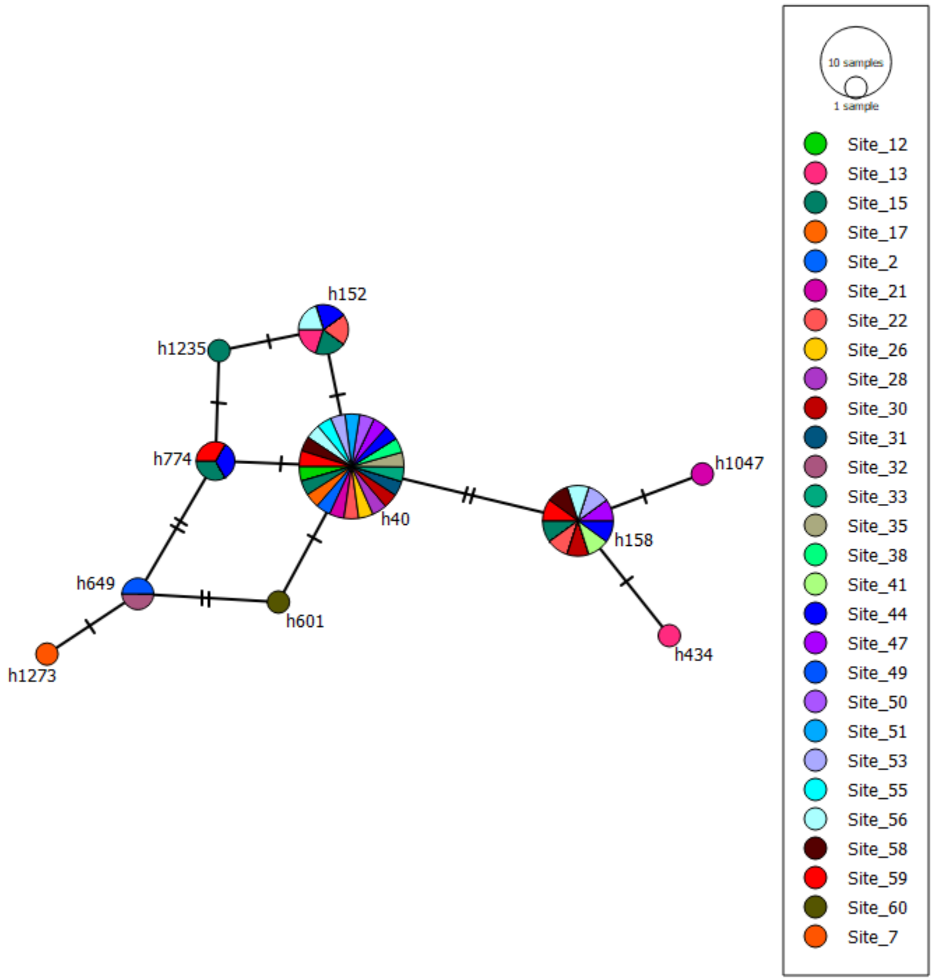

### OTU 25

#### Phylogeography

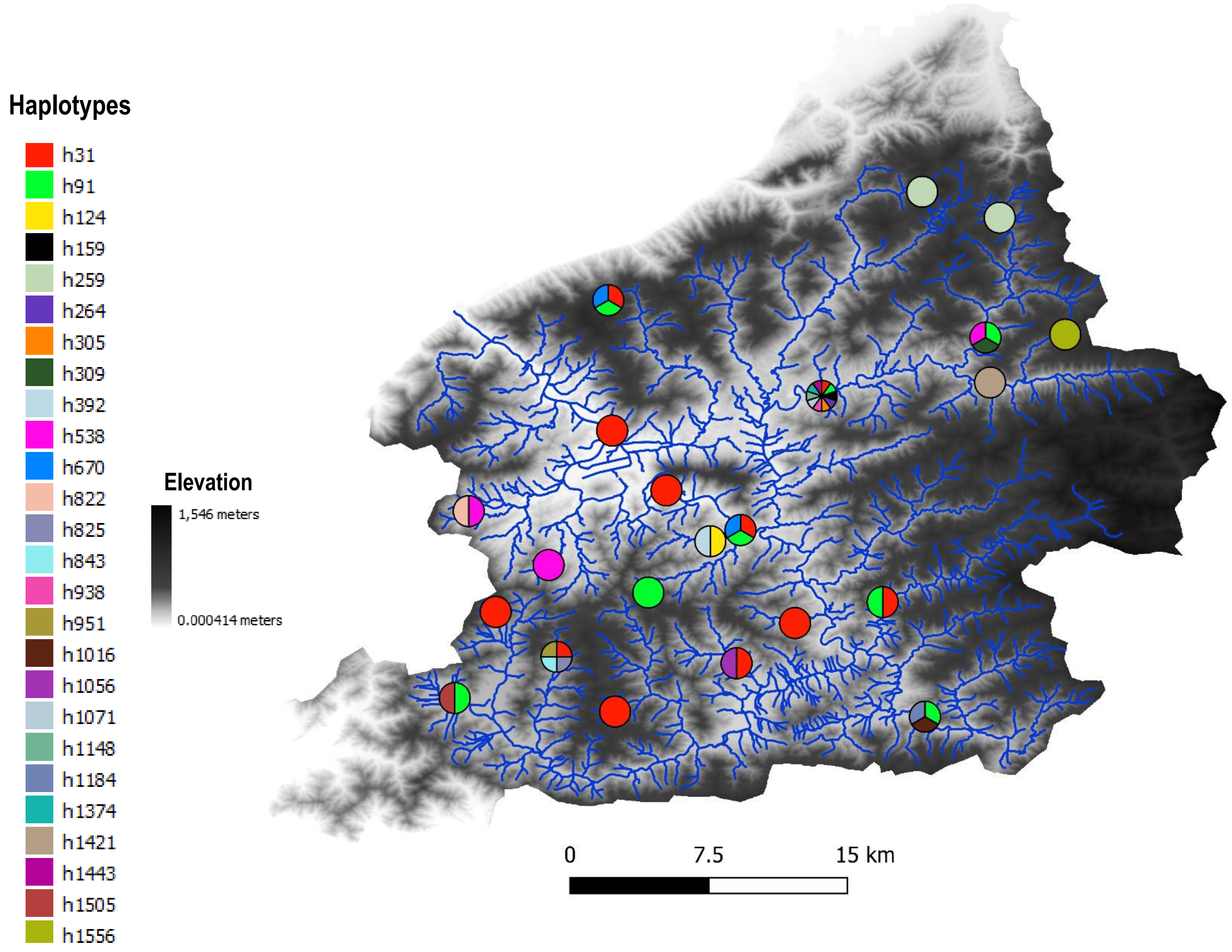


#### Haplotype network

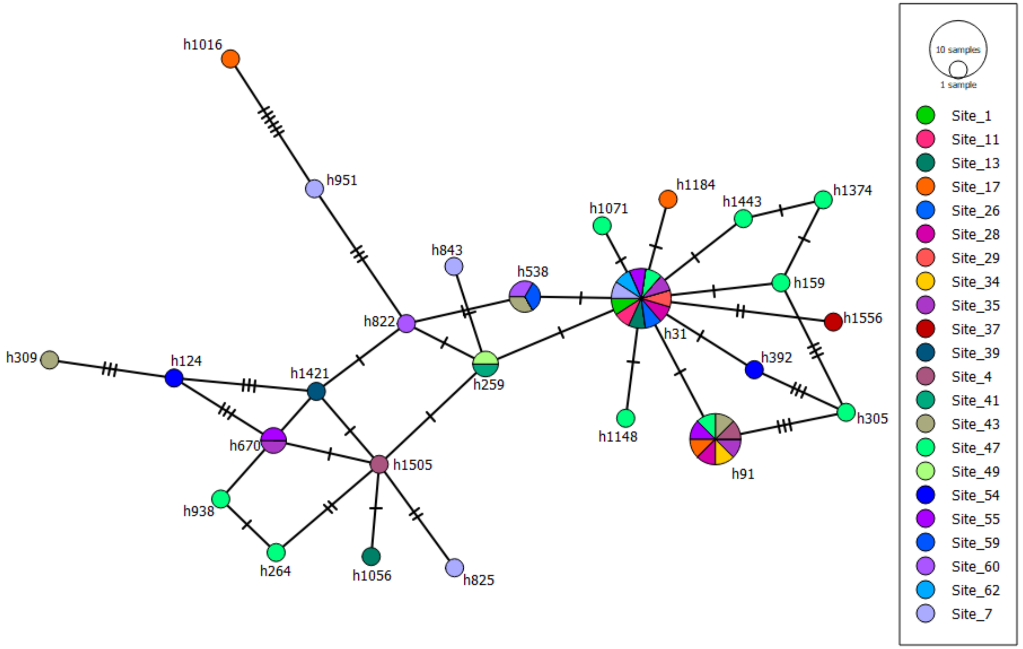

### OTU 21

#### Phylogeography

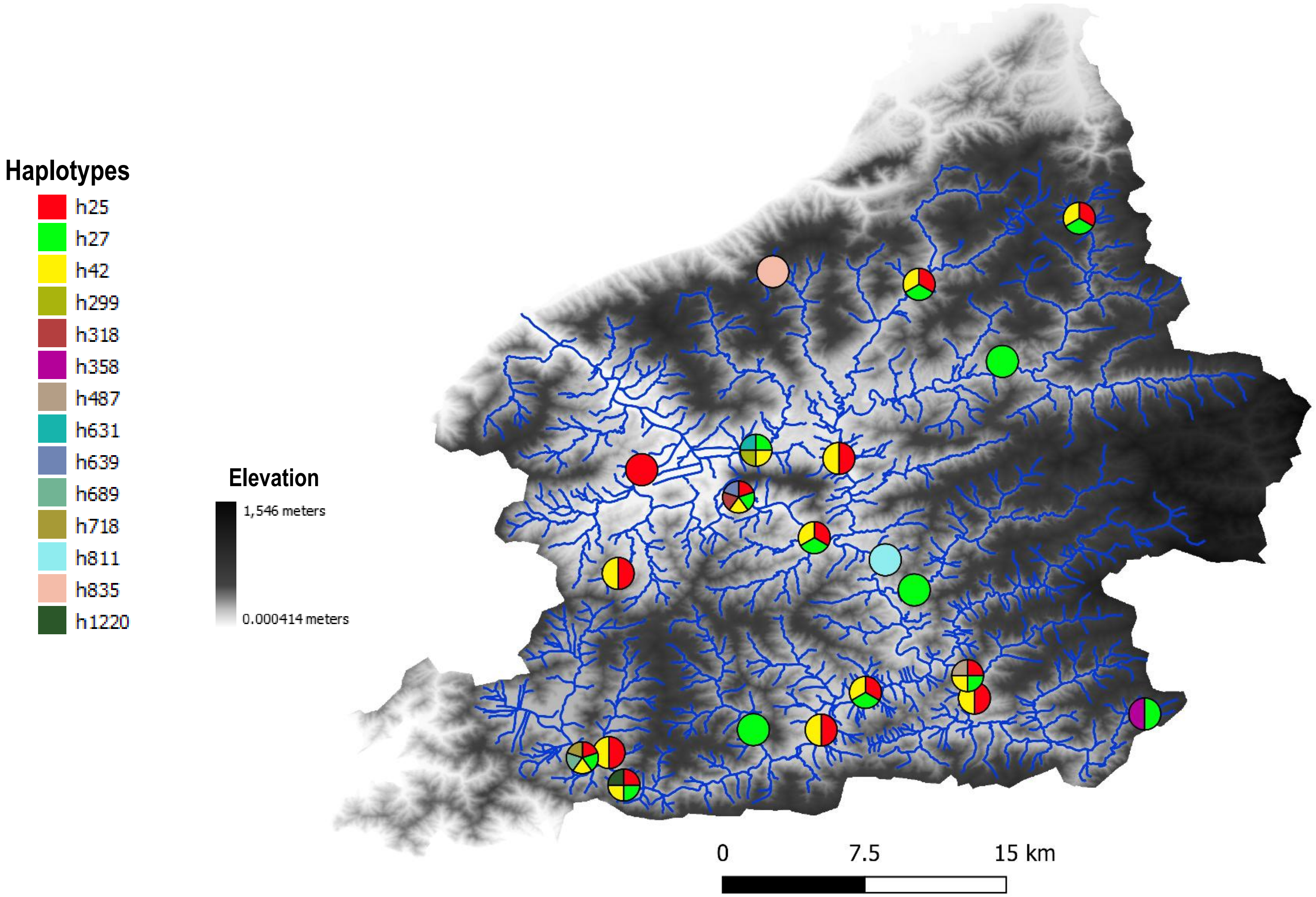


#### Haplotype network

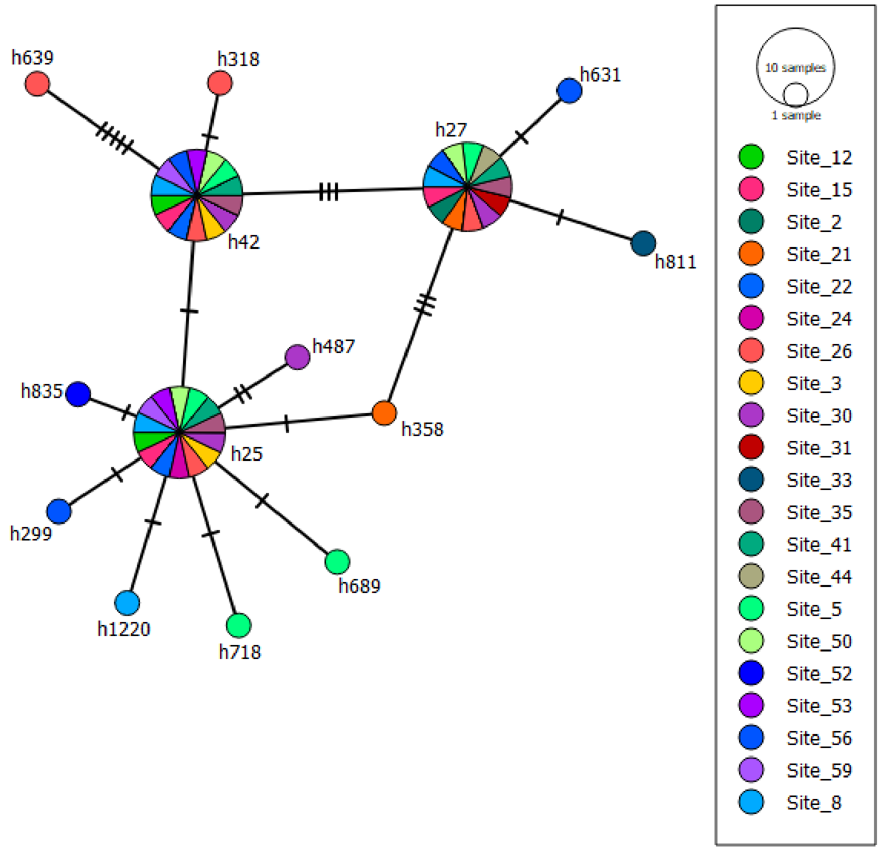

### OTU 9

#### Phylogeography

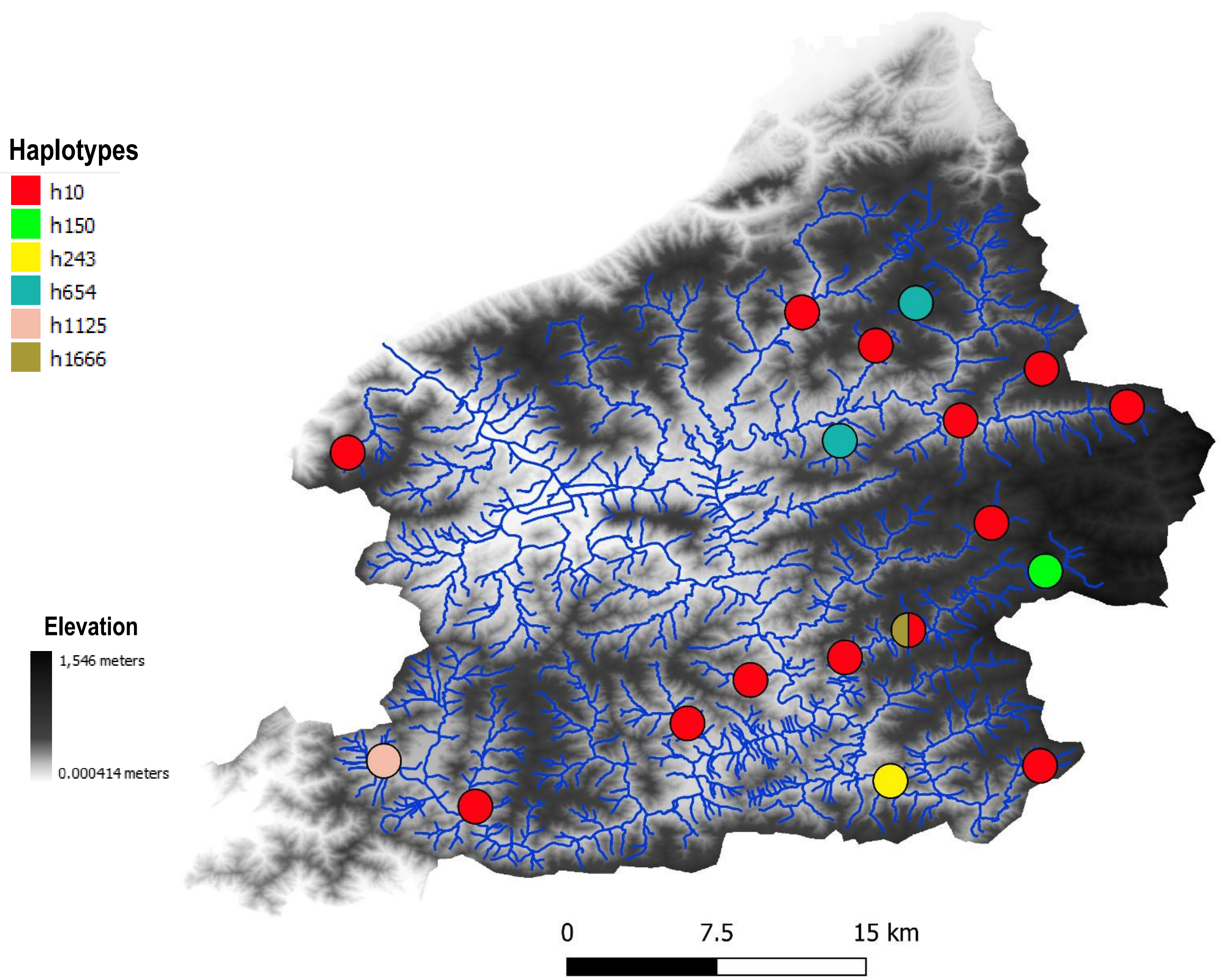


#### Haplotype network

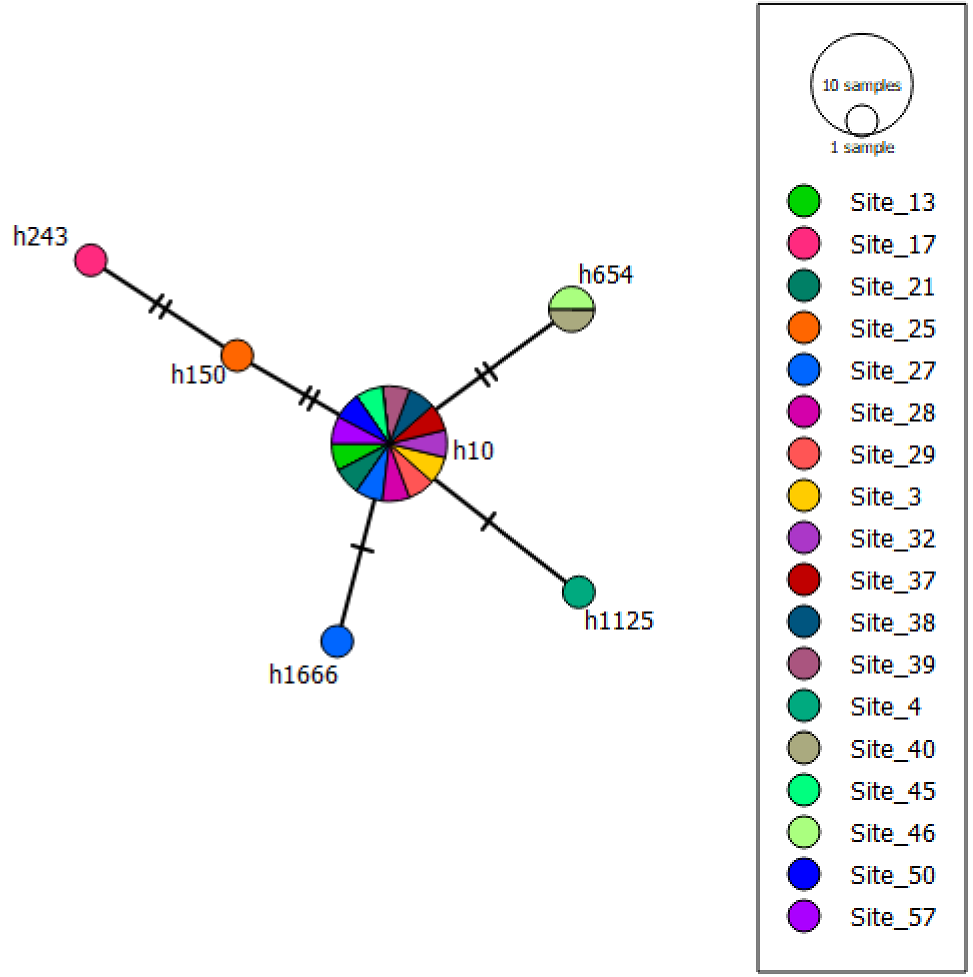

### OTU 23

#### Phylogeography

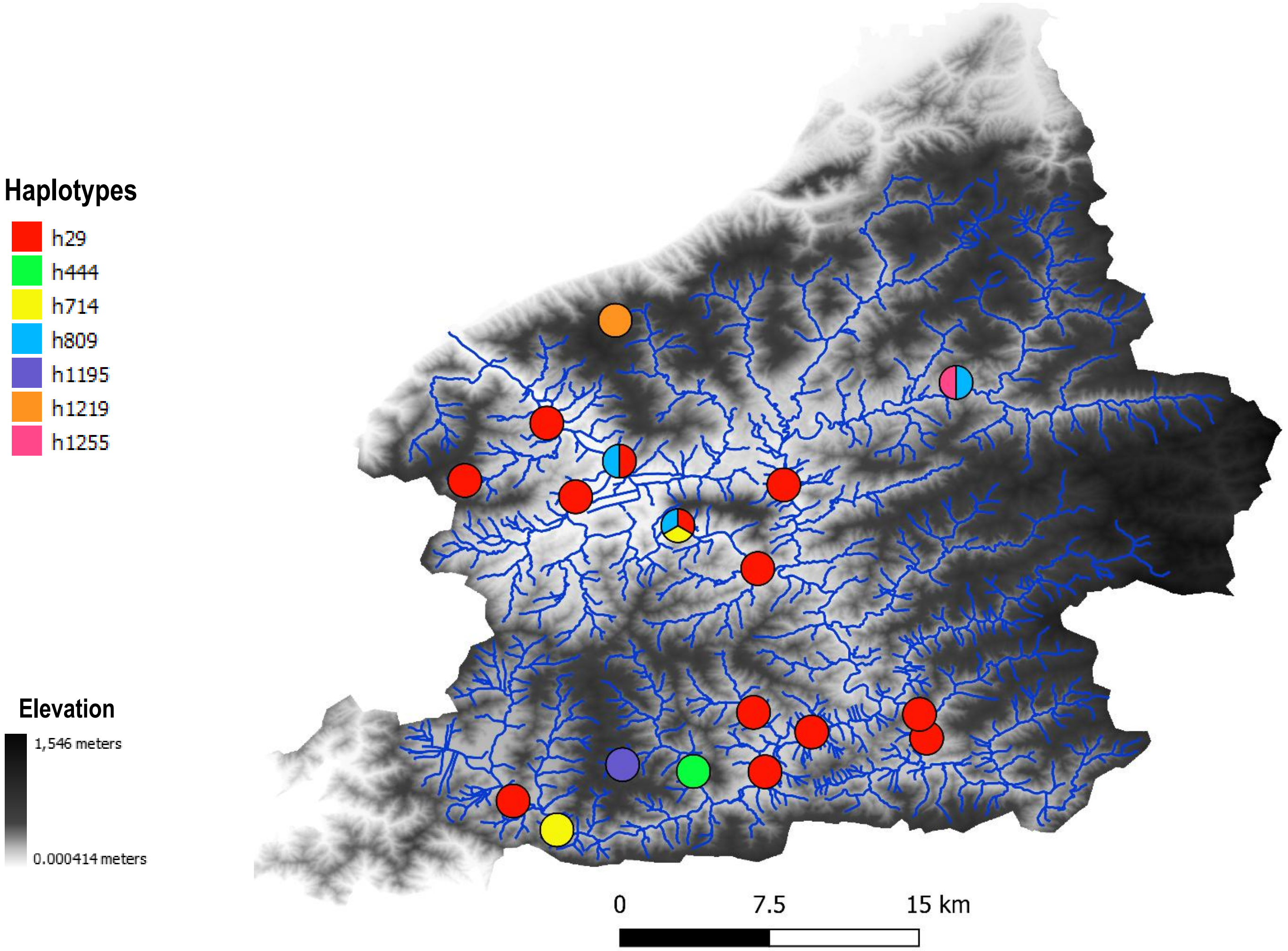


#### Haplotype network

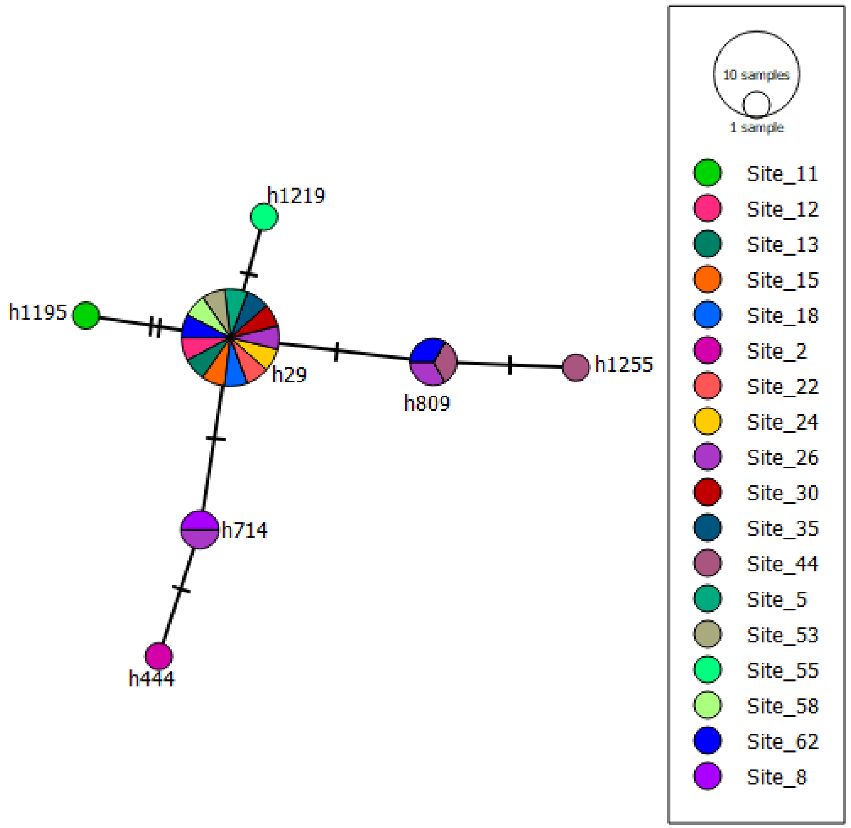

### OTU 3

#### Phylogeography

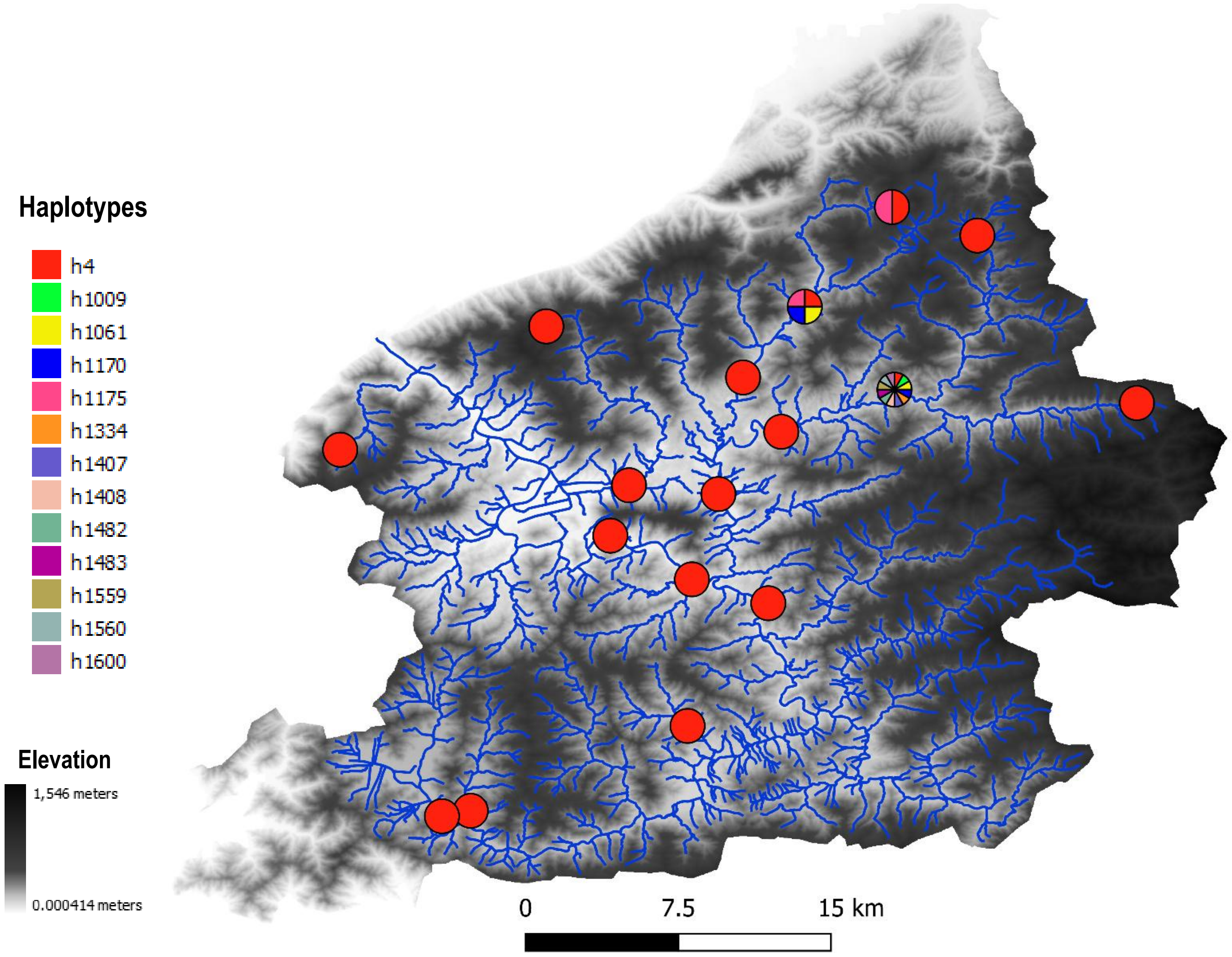


#### Haplotype network

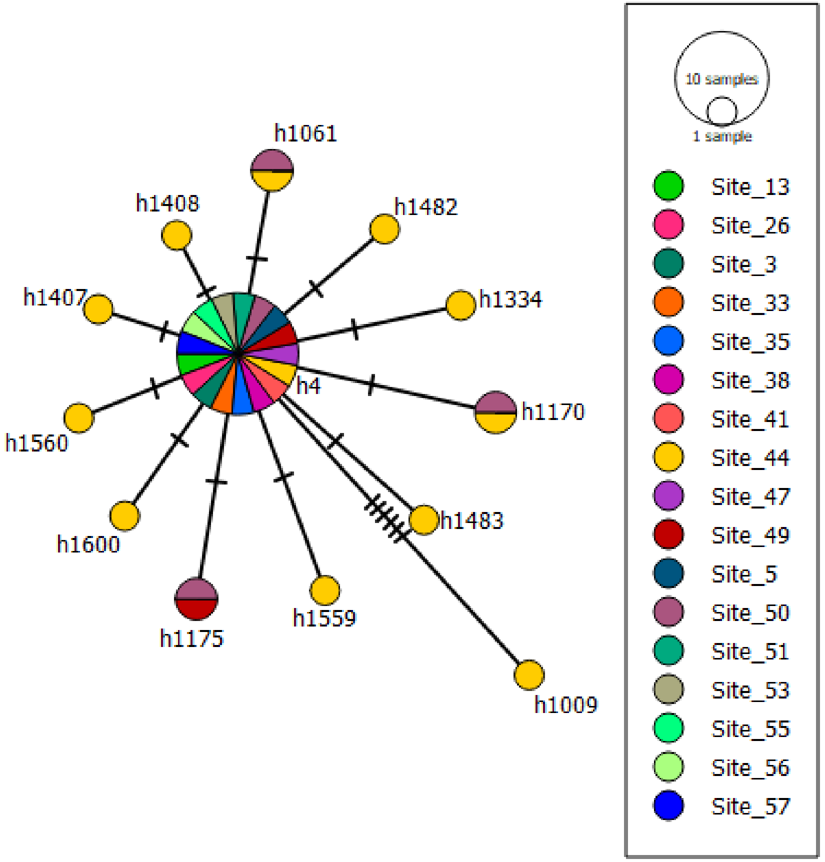

### OTU 13

#### Phylogeography

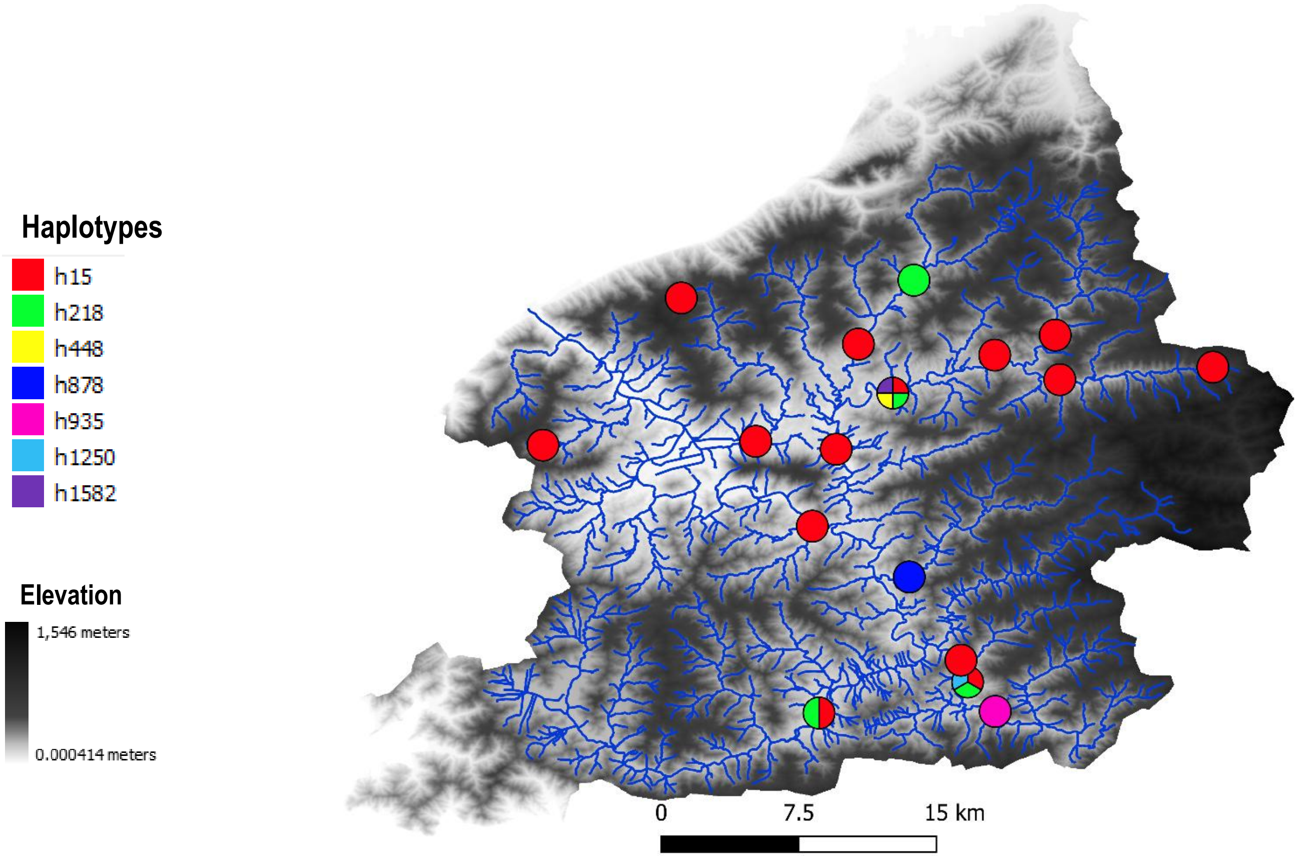


#### Haplotype network

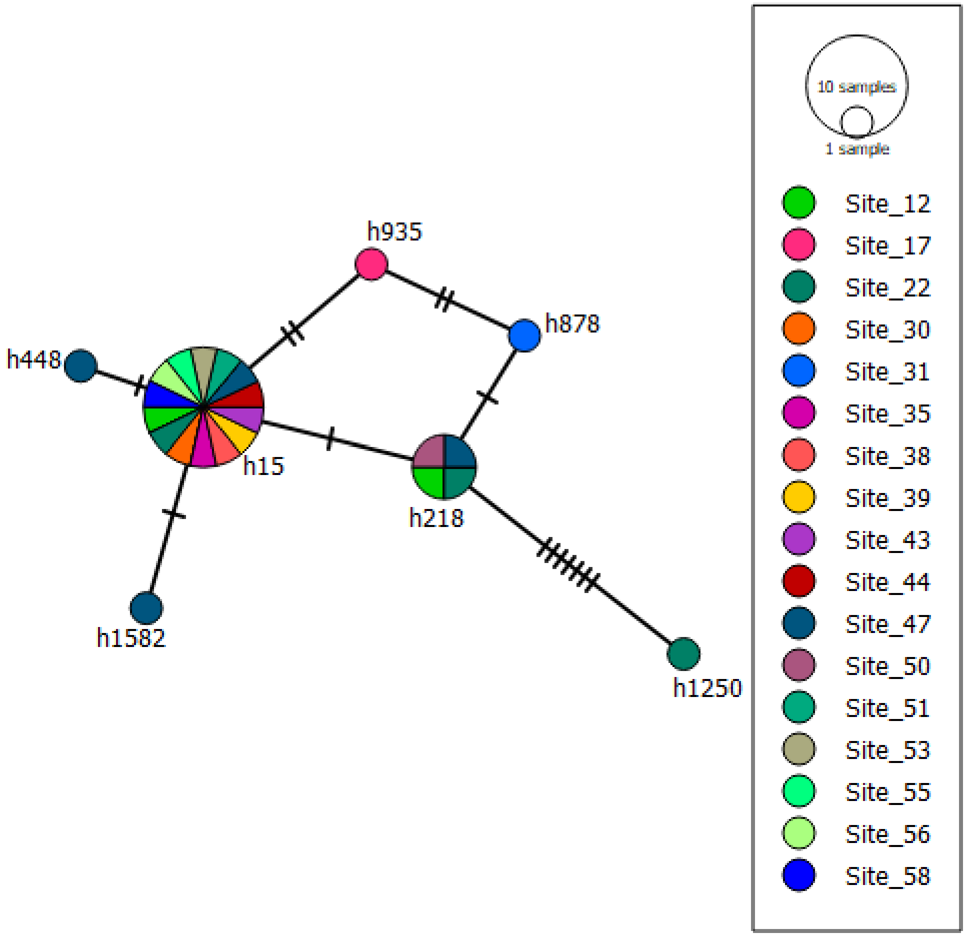

### OTU 75

#### Phylogeography

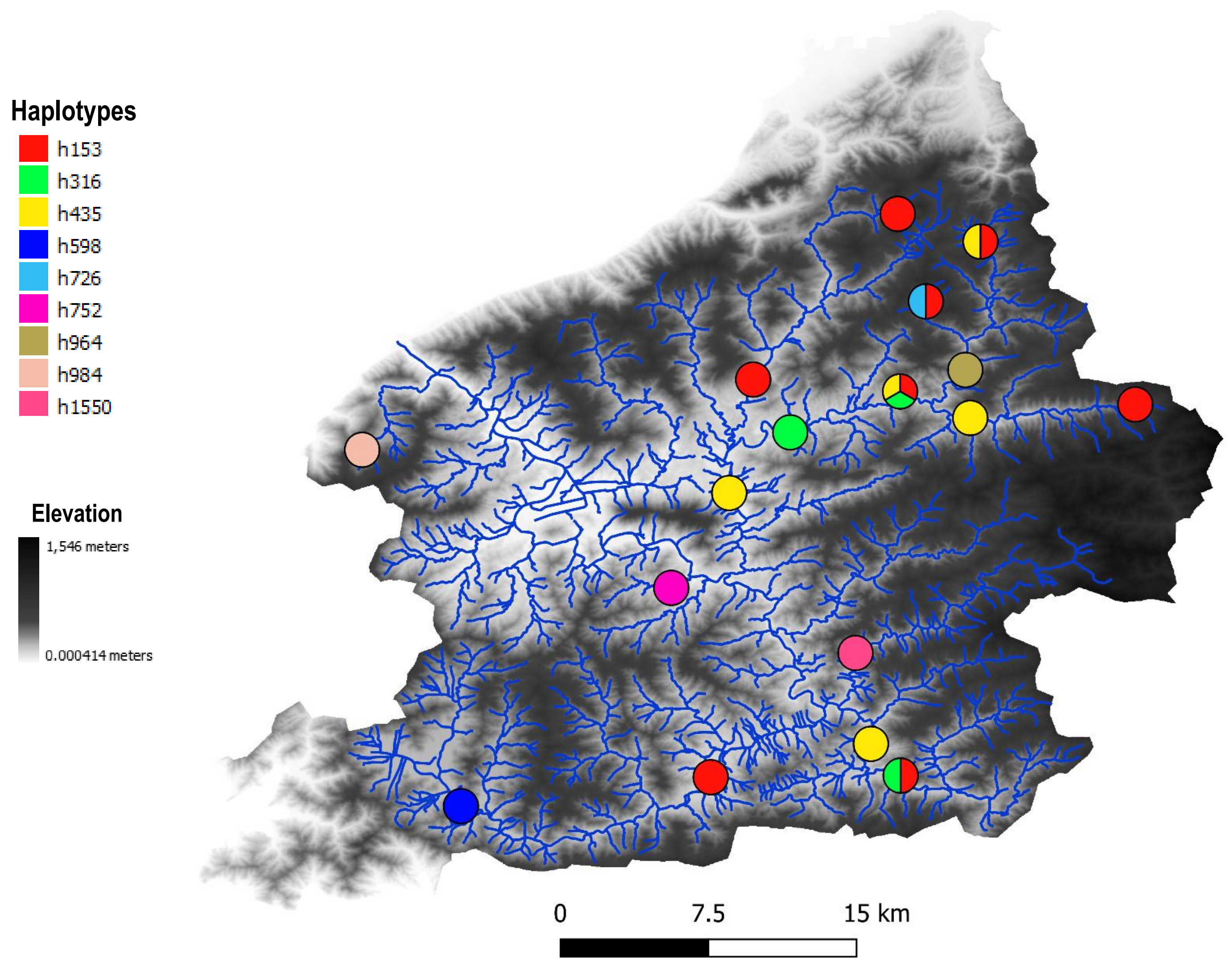


#### Haplotype network

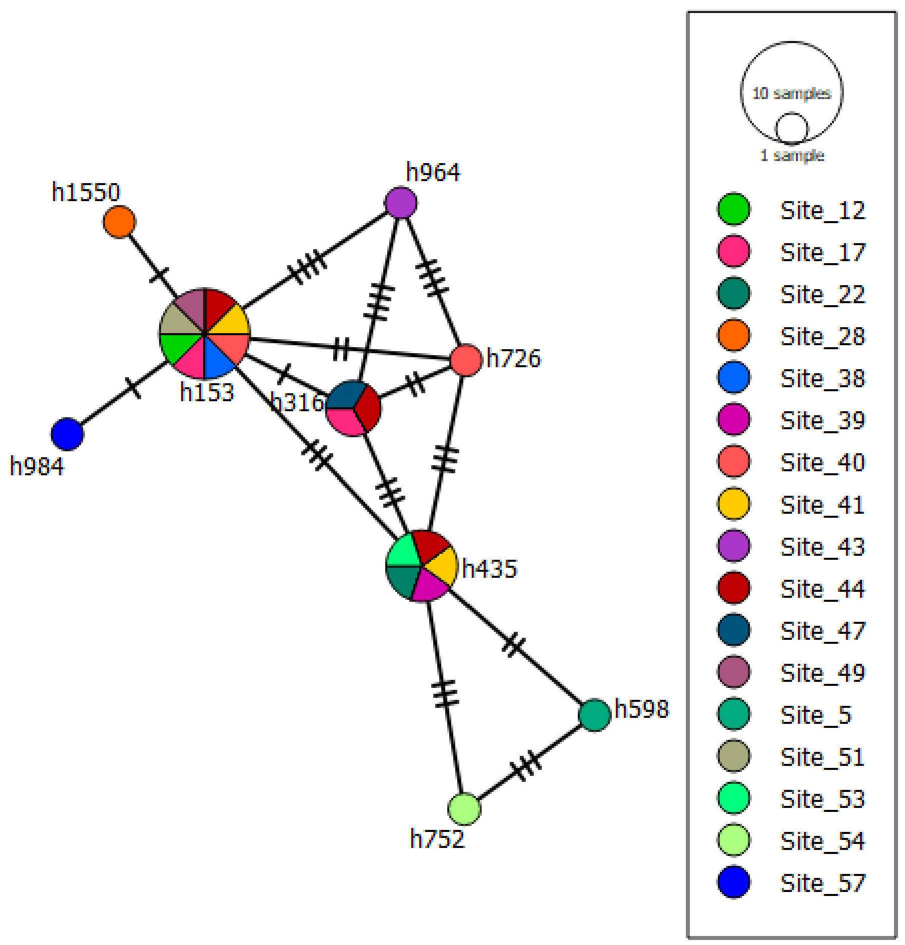

### OTU 36

#### Phylogeography

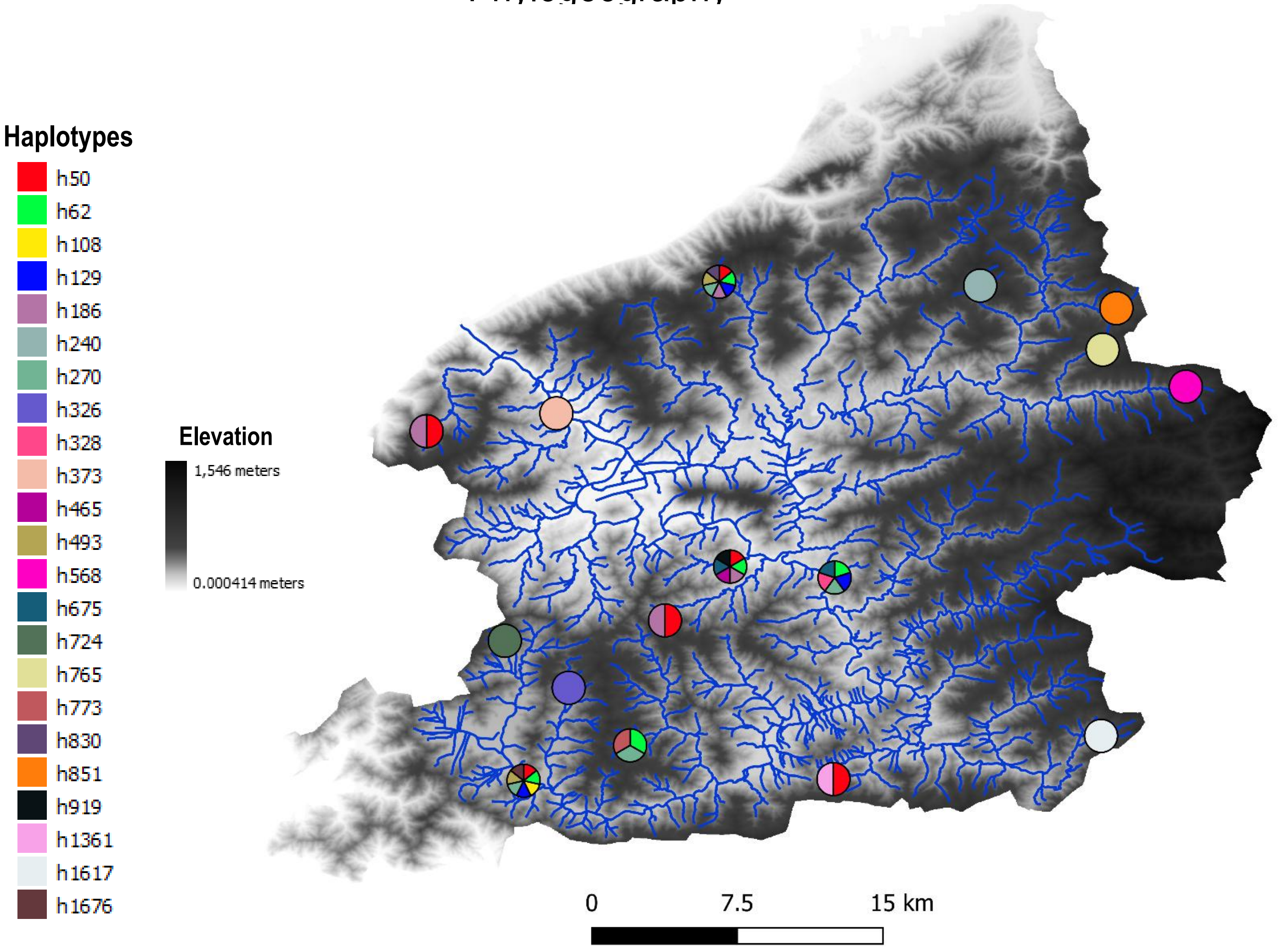


#### Haplotype network

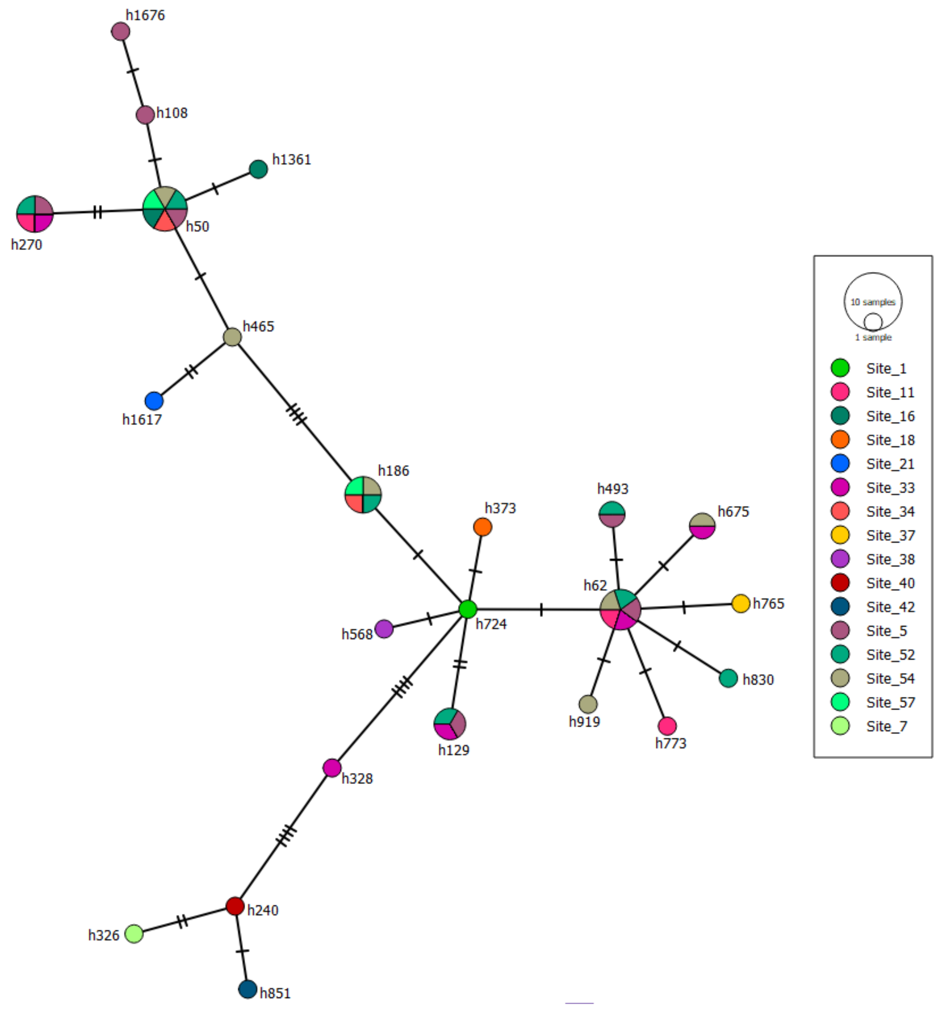

### OTU 1

#### Phylogeography

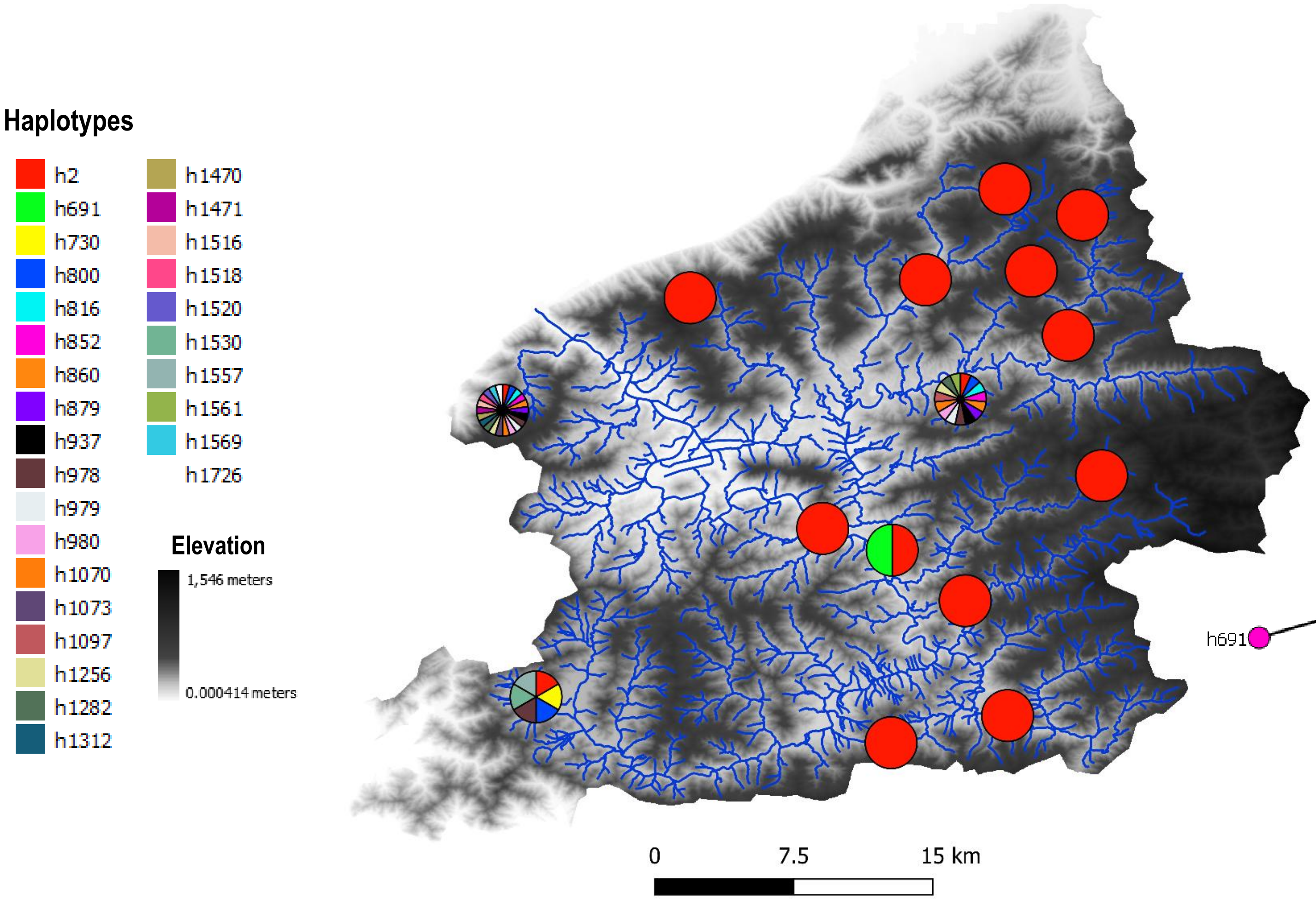


#### Haplotype network

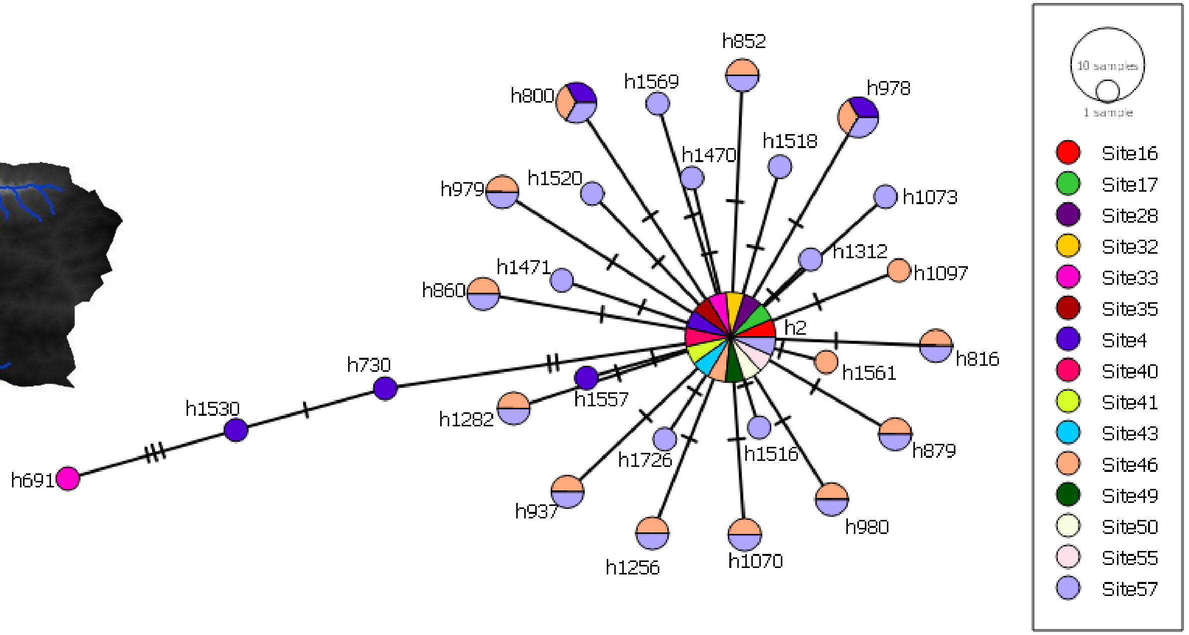

### OTU 8

#### Phylogeography

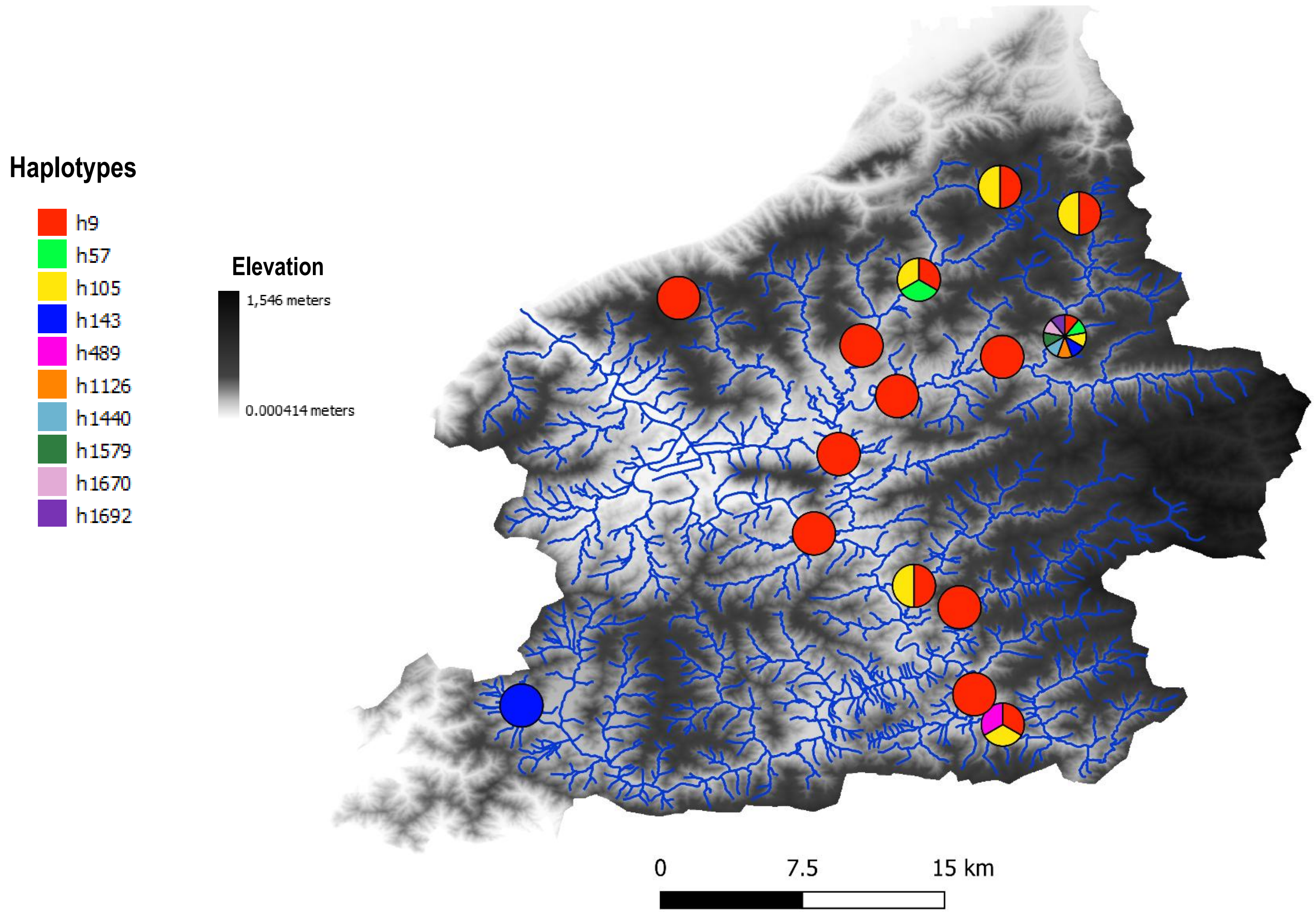


#### Haplotype network

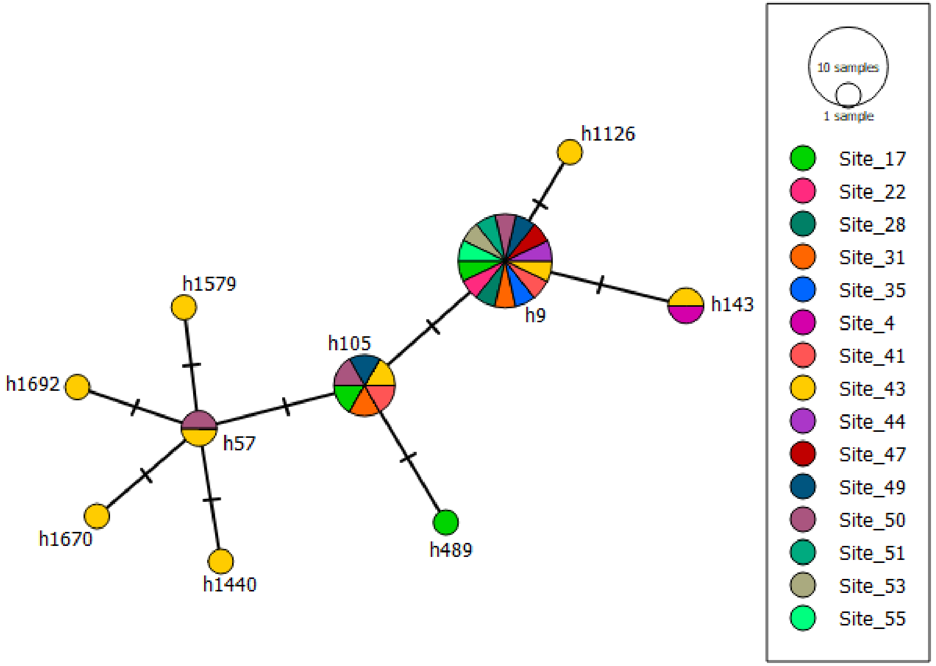

### OTU 37

#### Phylogeography

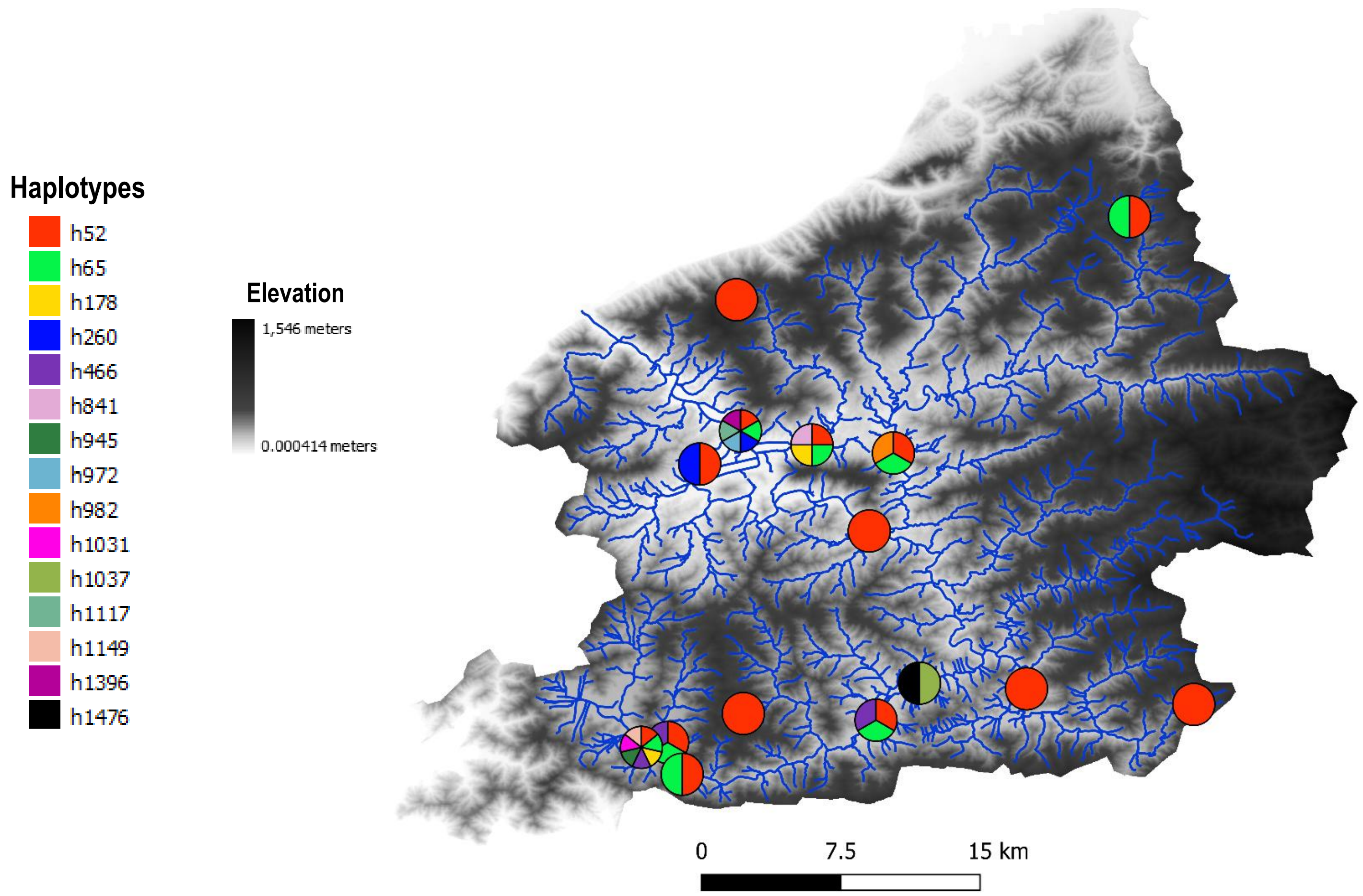


#### Haplotype network

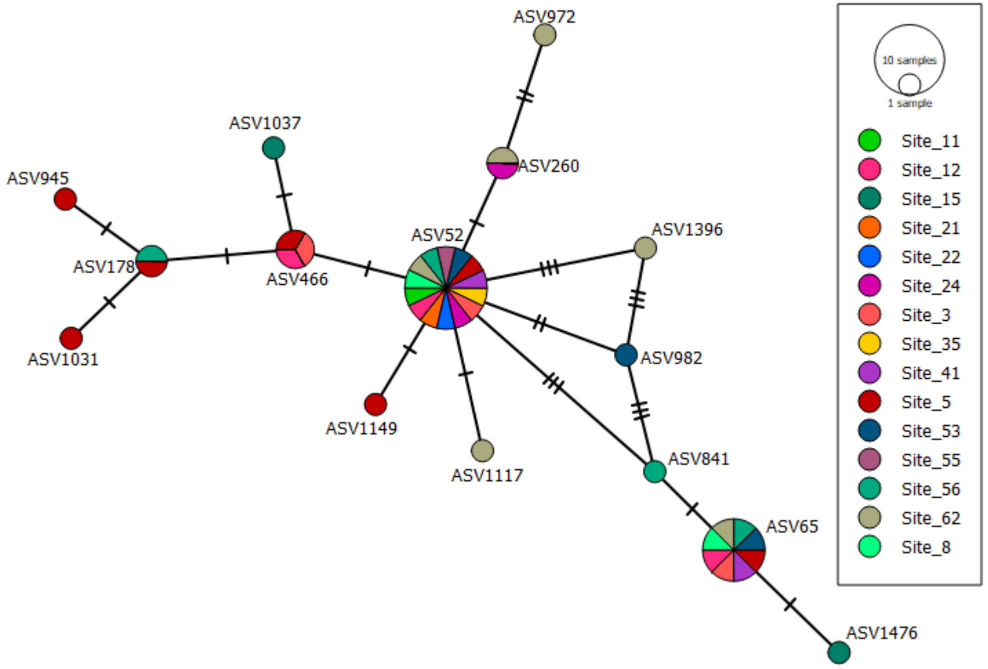

### OTU 47

#### Phylogeography

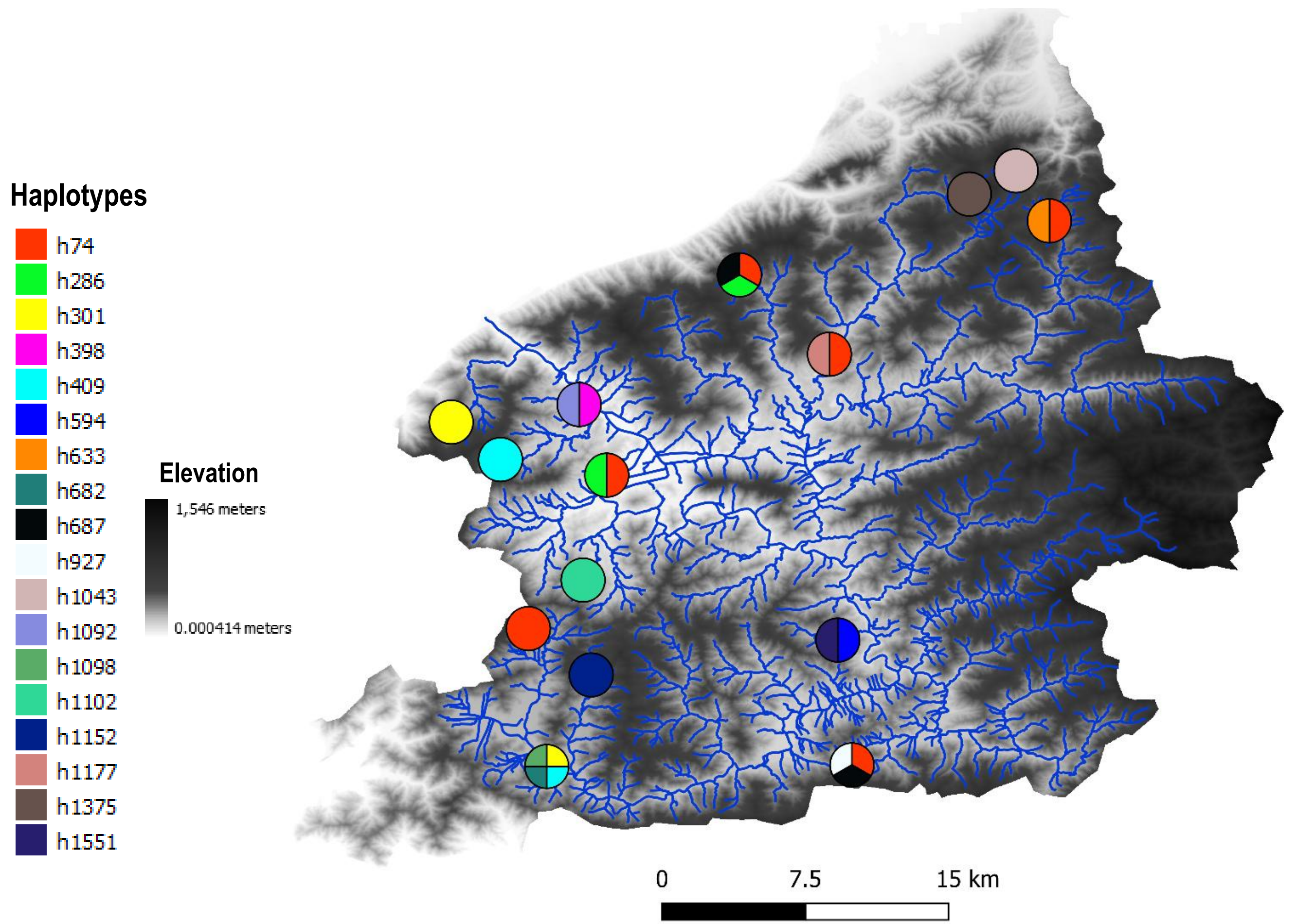


#### Haplotype network

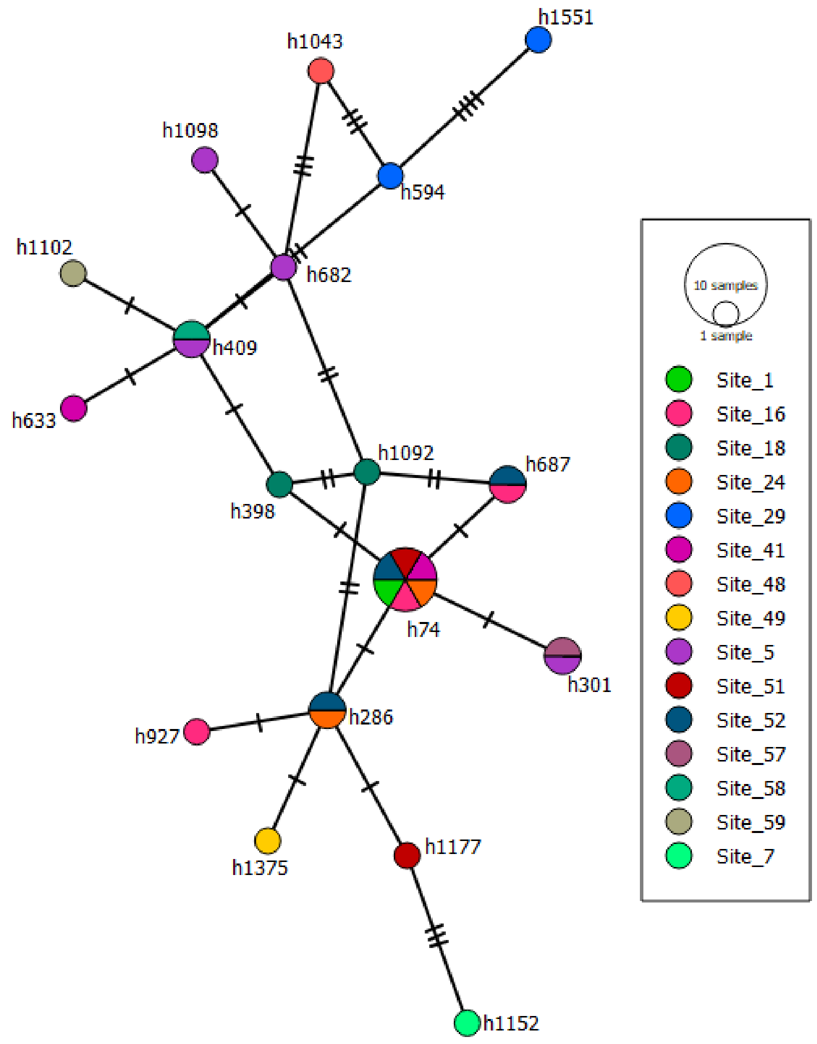

### OTU 11

#### Phylogeography

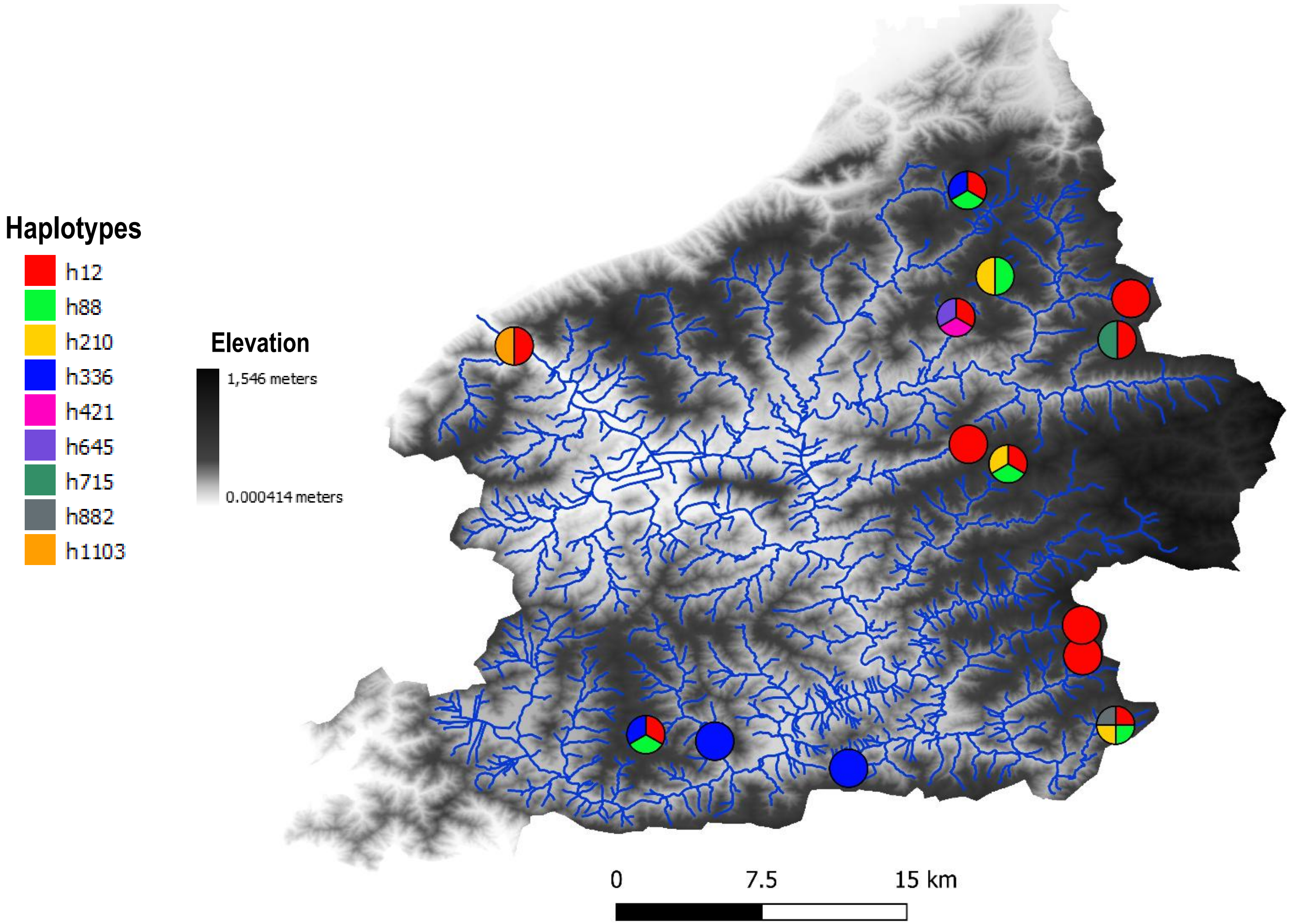


#### Haplotype network

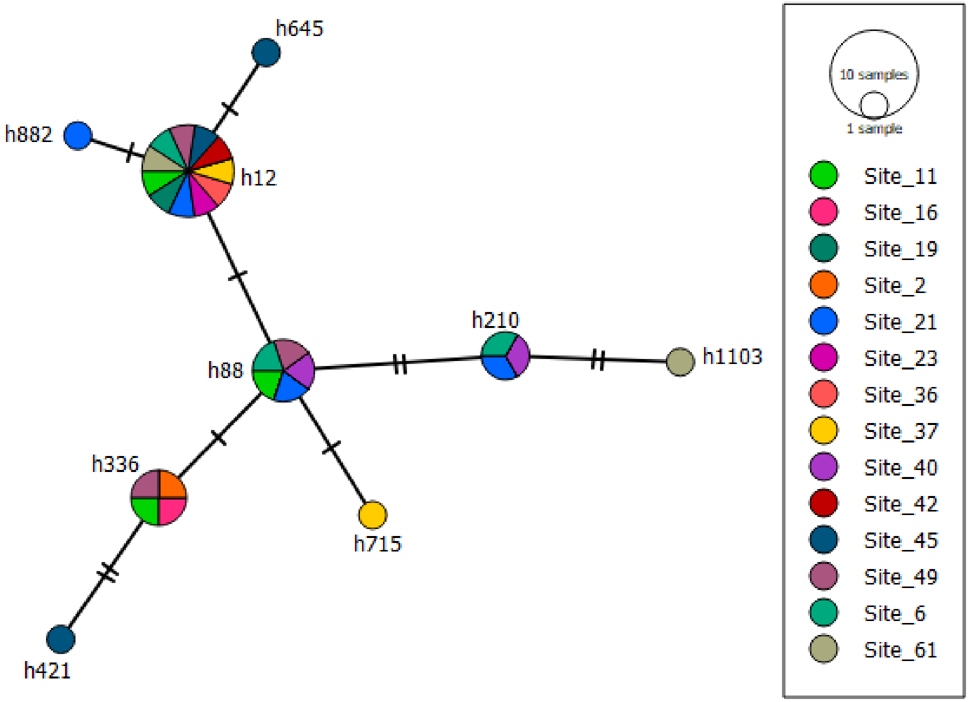

### OTU 57

#### Phylogeography

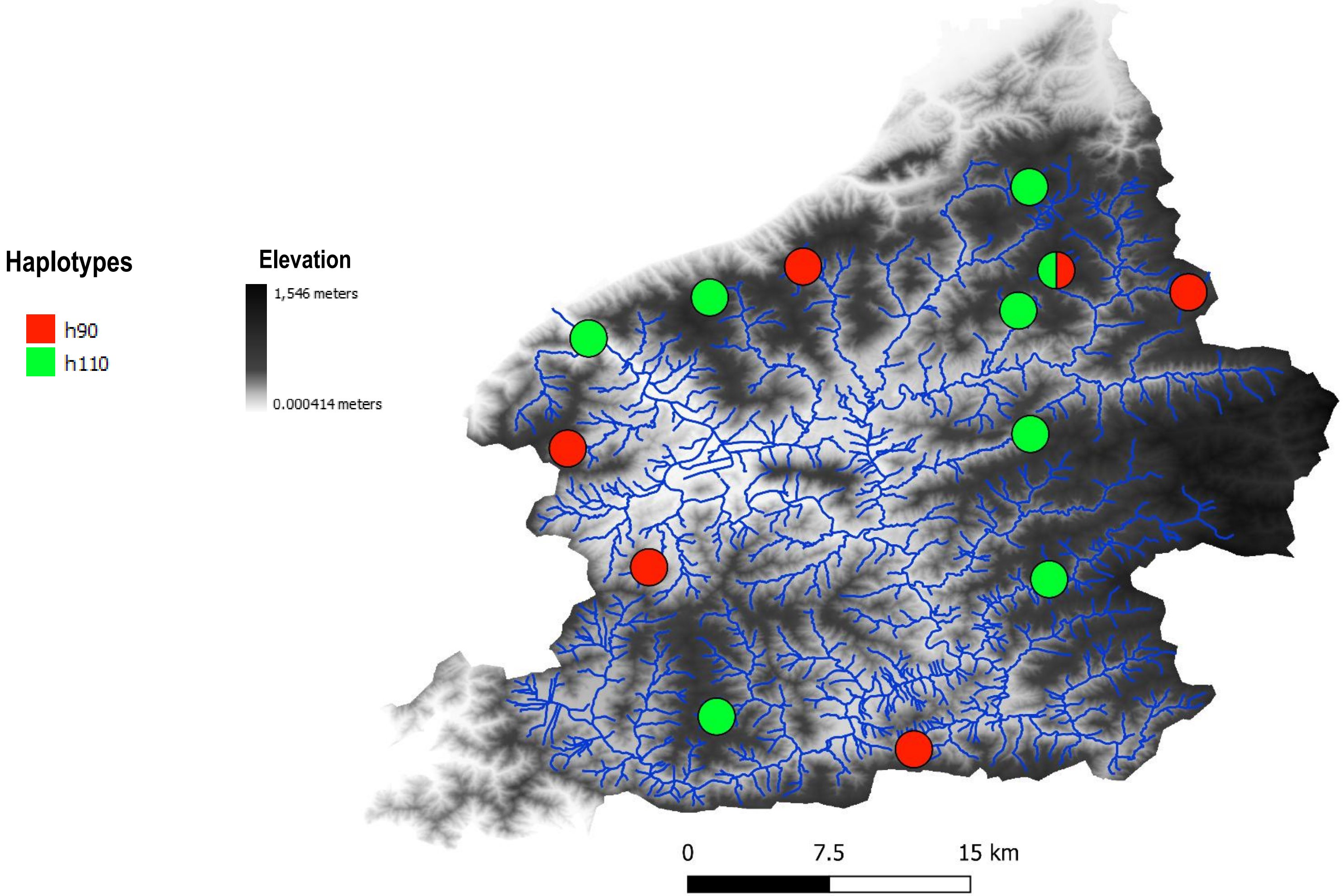


#### Haplotype network

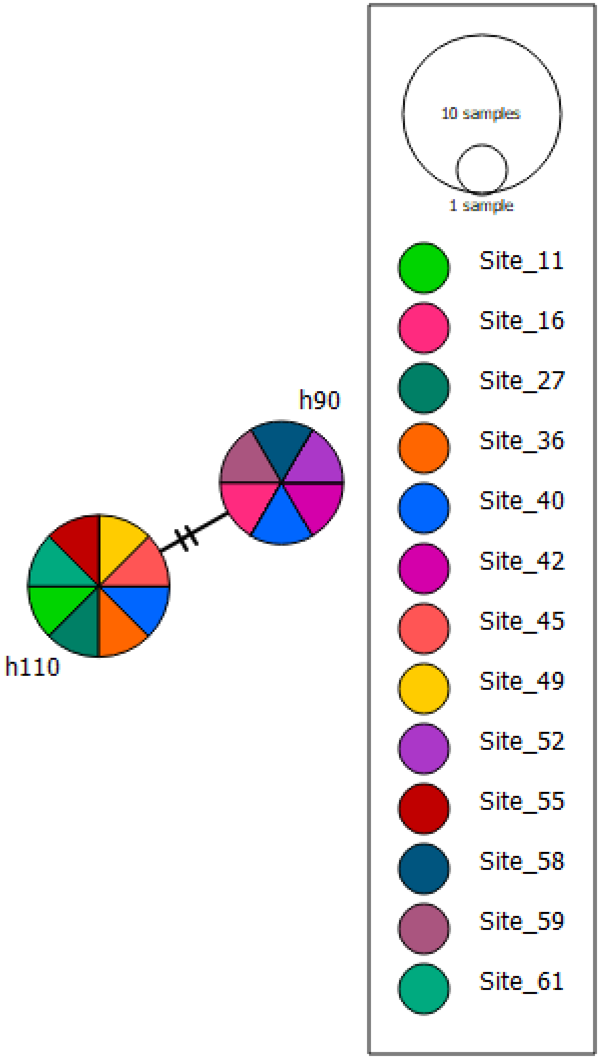

OTU 22

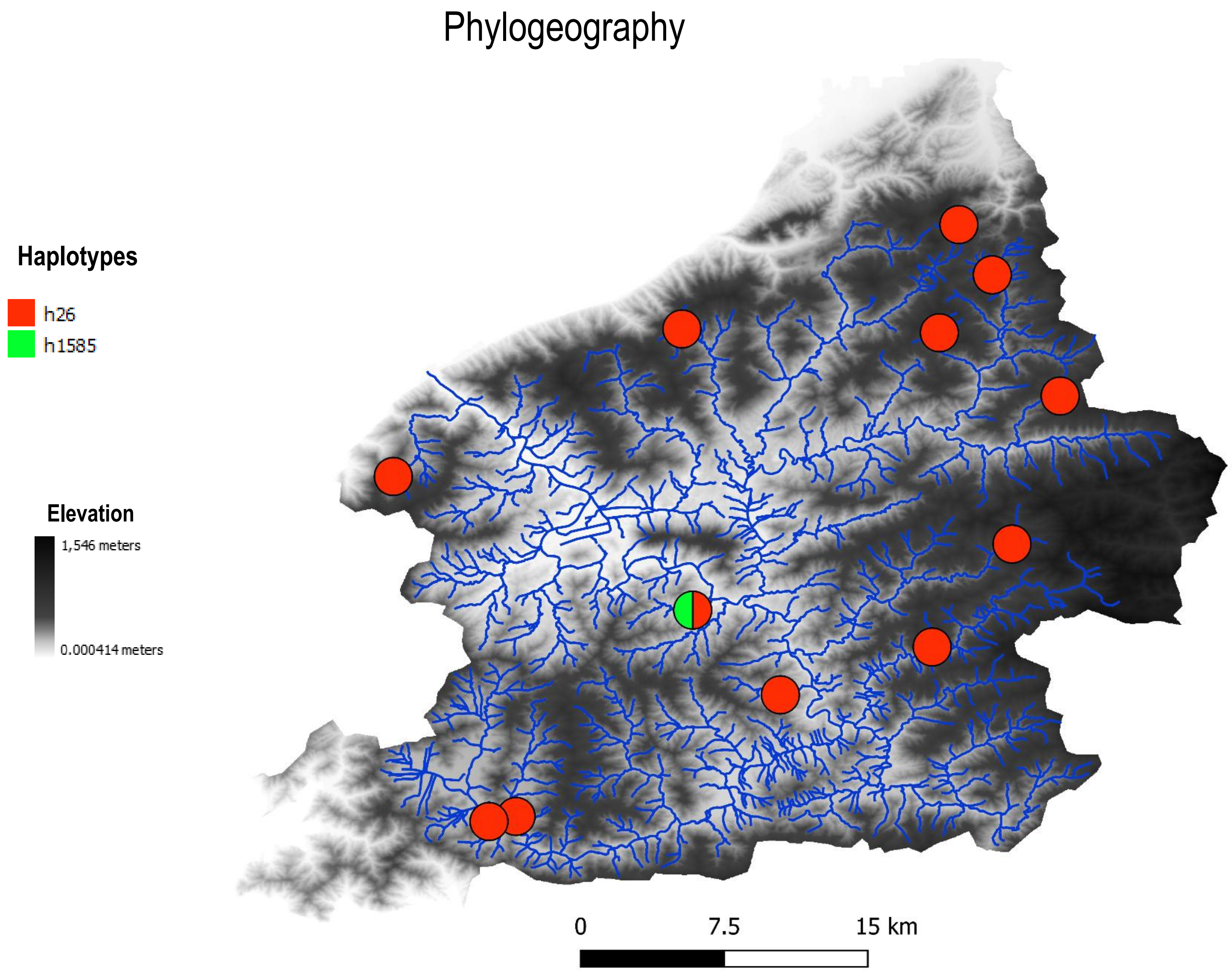

Phylogeography
Haplotypes
h26
h1585
Elevation
1,546 meters
0.000414 meters
0
7.5
15 km


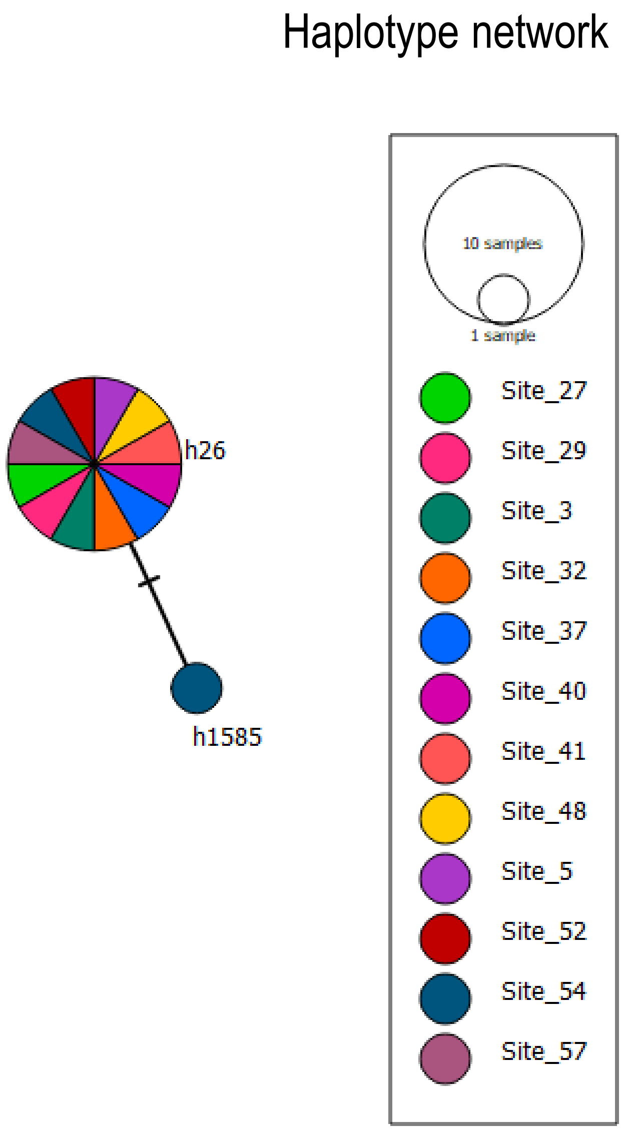

Haplotype network
h26
h1585
10 samples
1 sample
Site_27
Site_29
Site_3
Site_32
Site_37
Site_40
Site_41
Site_48
Site_5
Site_52
Site_54
Site_57

### OTU 60

#### Phylogeography

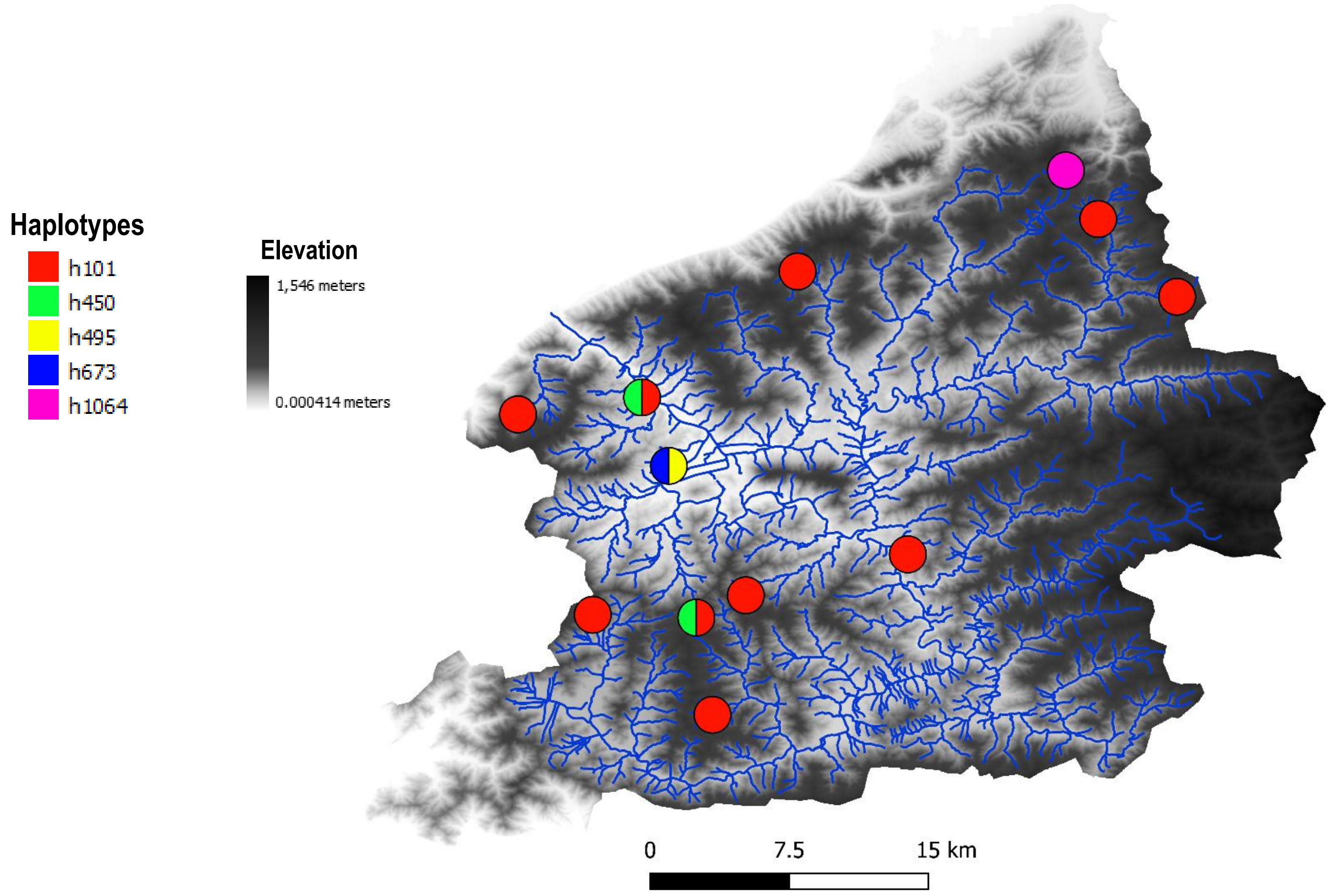


#### Haplotype network

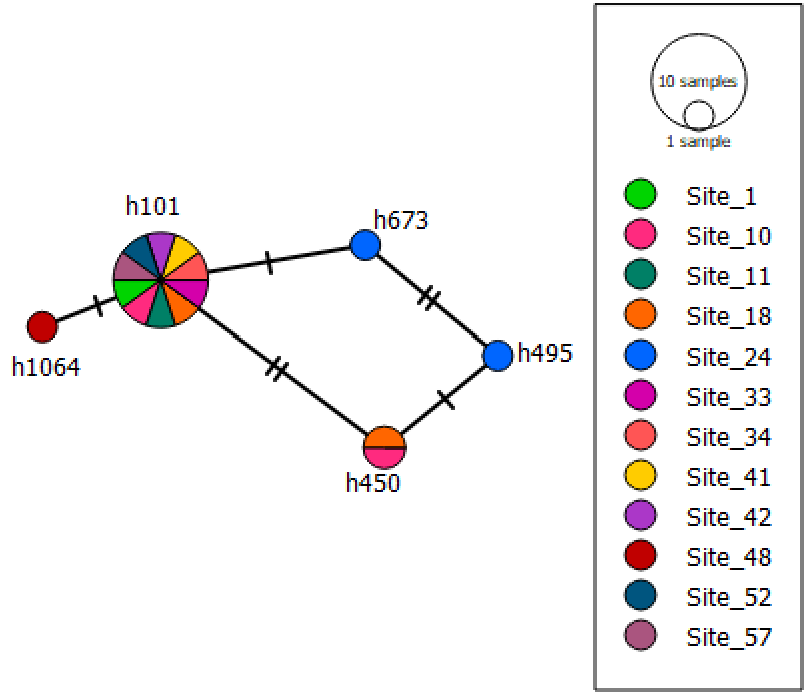

### OTU 33

#### Phylogeography

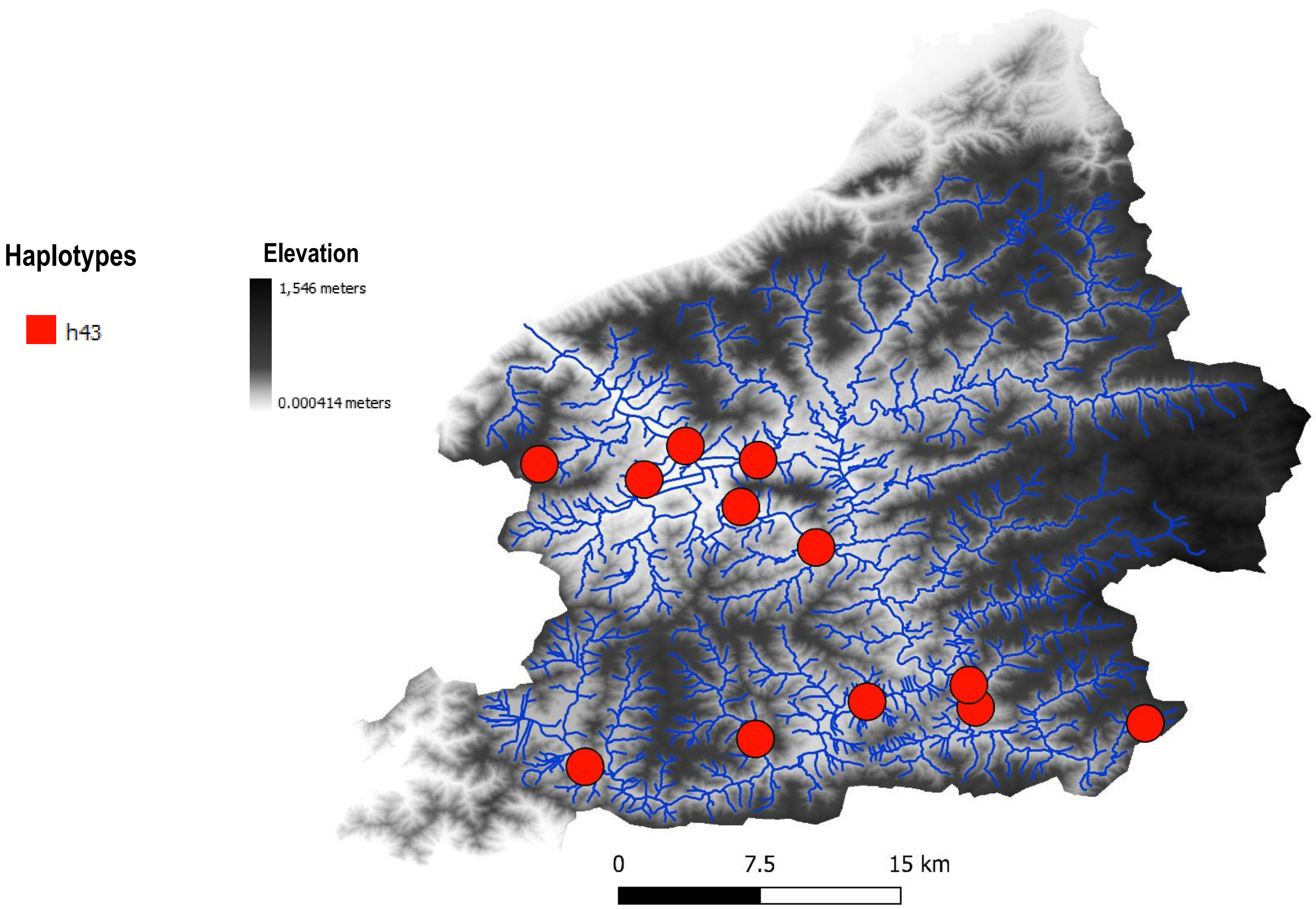


#### Haplotype network

### OTU 34

#### Phylogeography

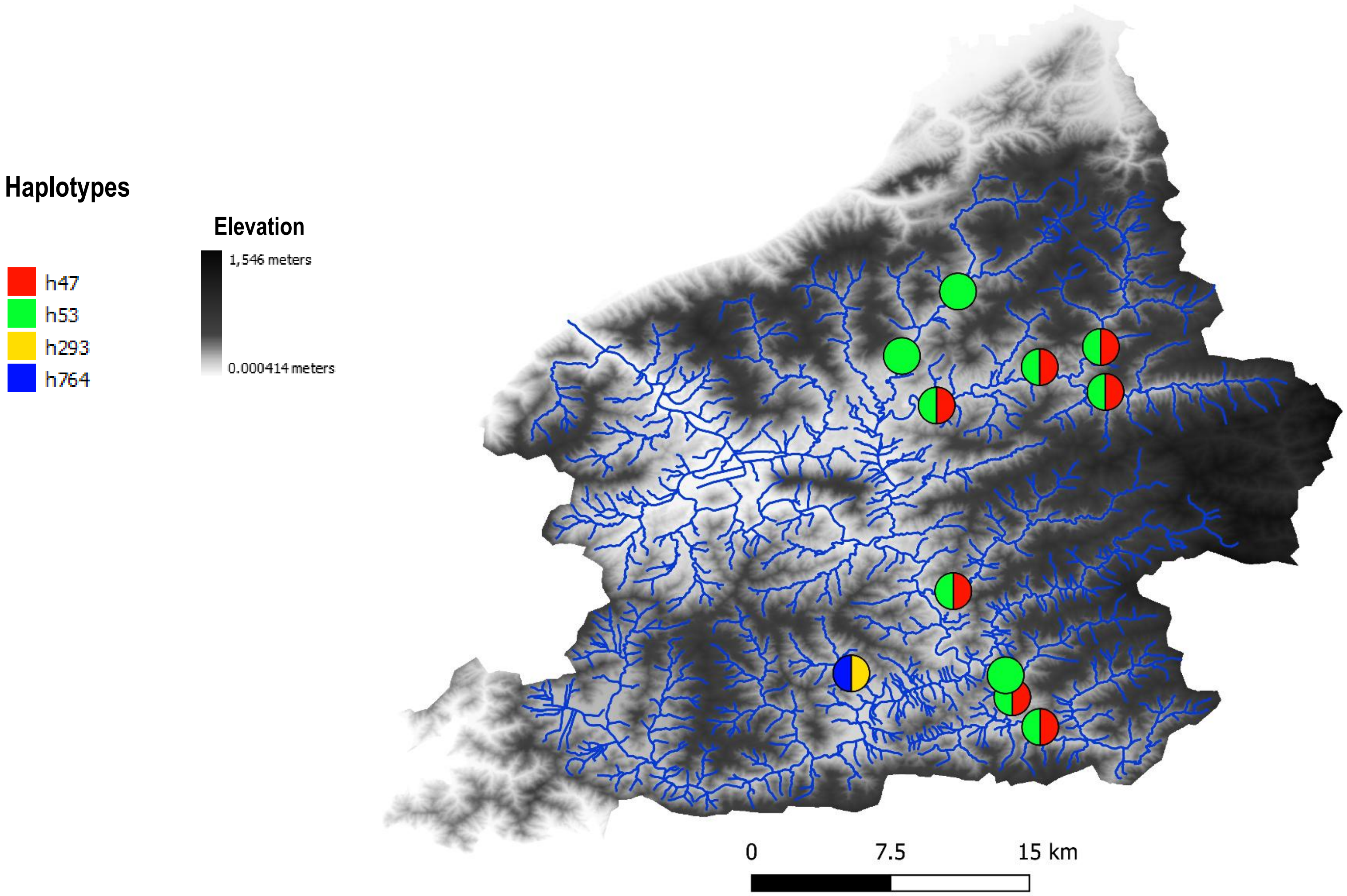


#### Haplotype network

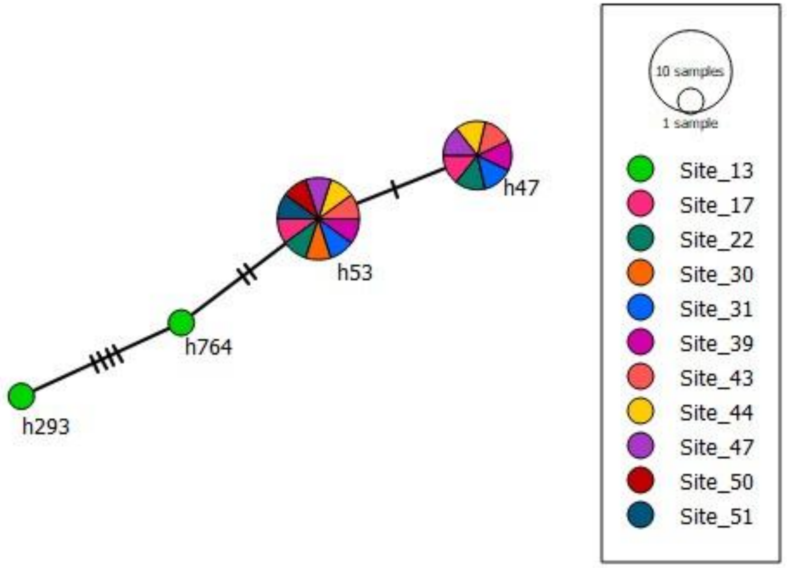

### OTU 51

#### Phylogeography

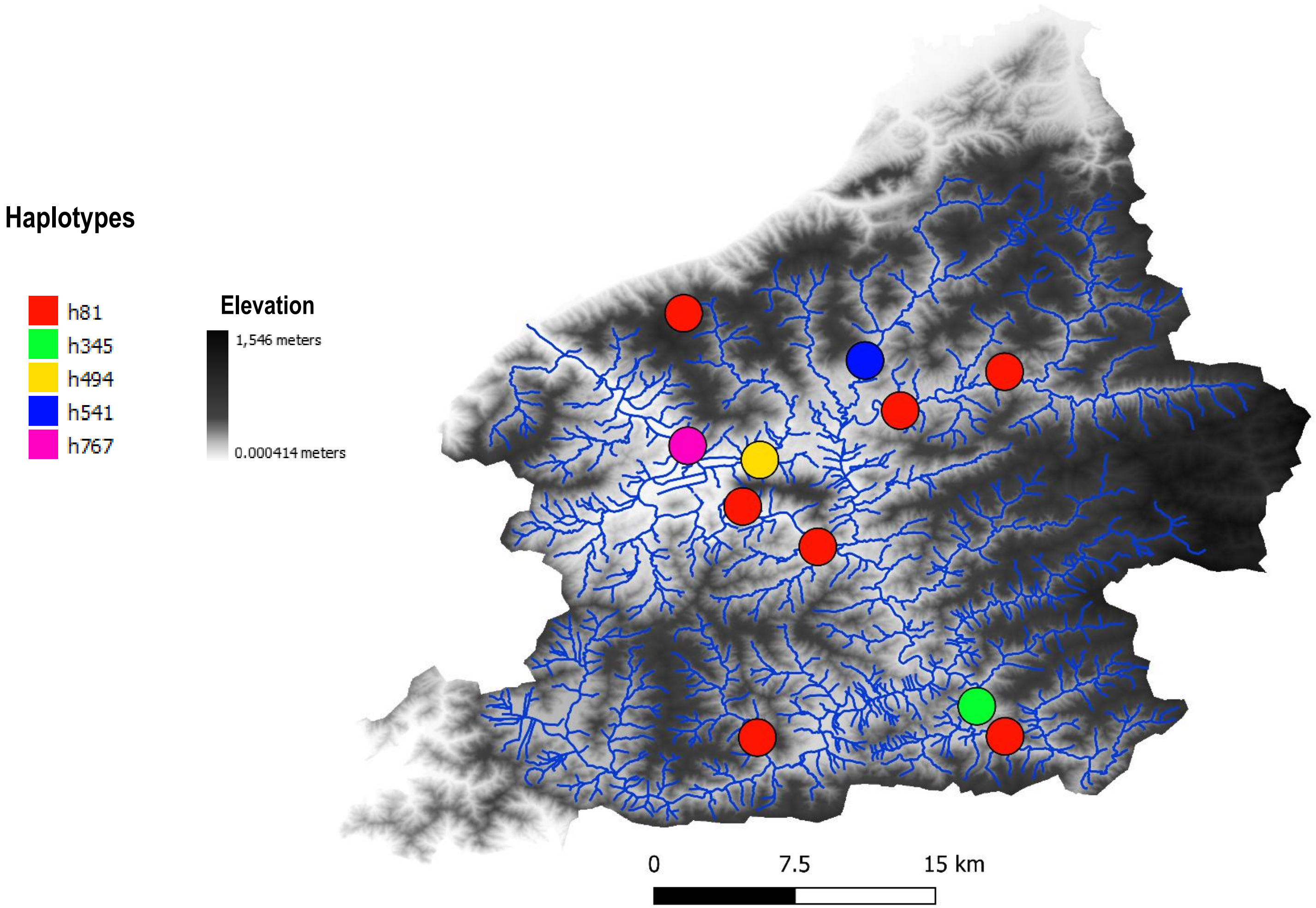


#### Haplotype network

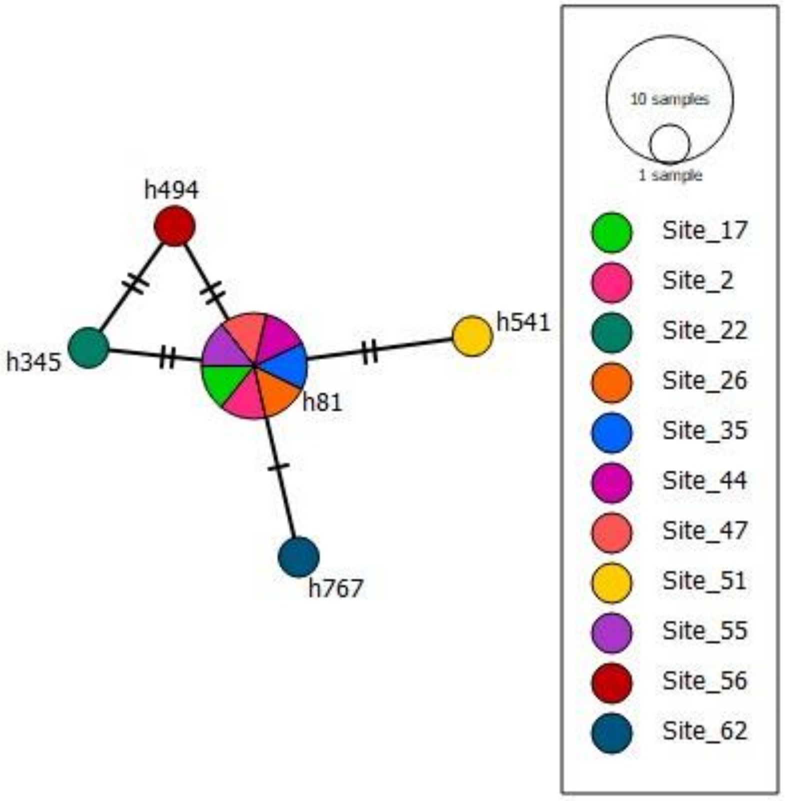

### OTU 28

#### Phylogeography

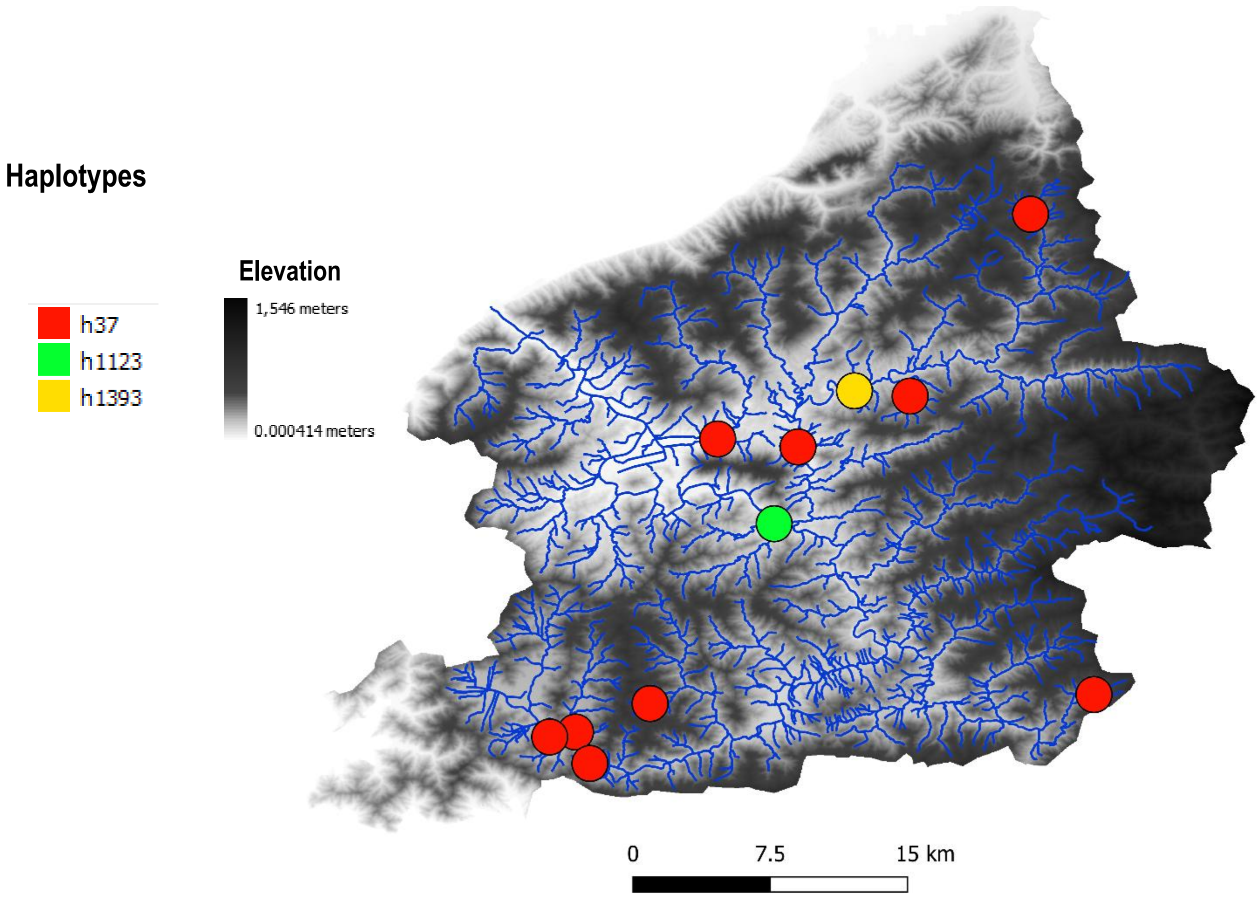


#### Haplotype network

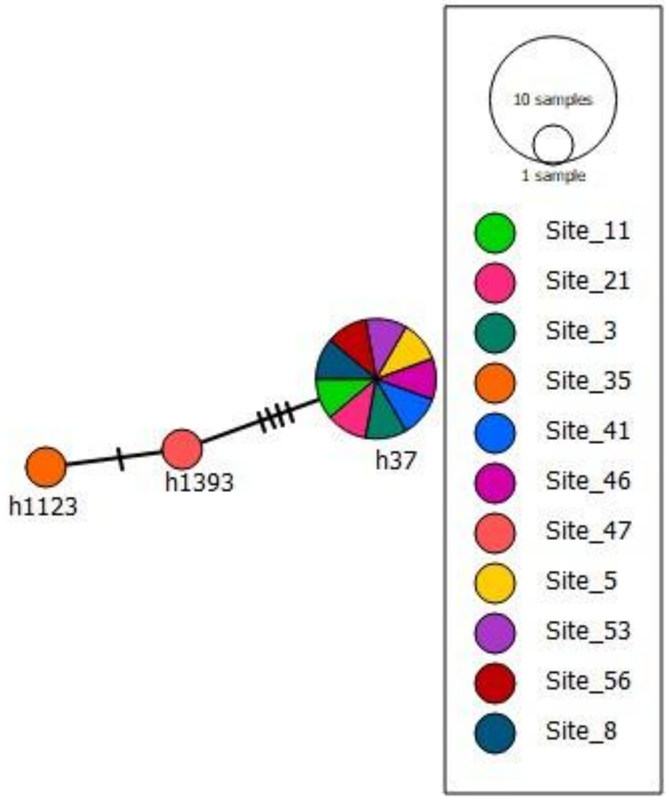

### OTU 38

#### Phylogeography

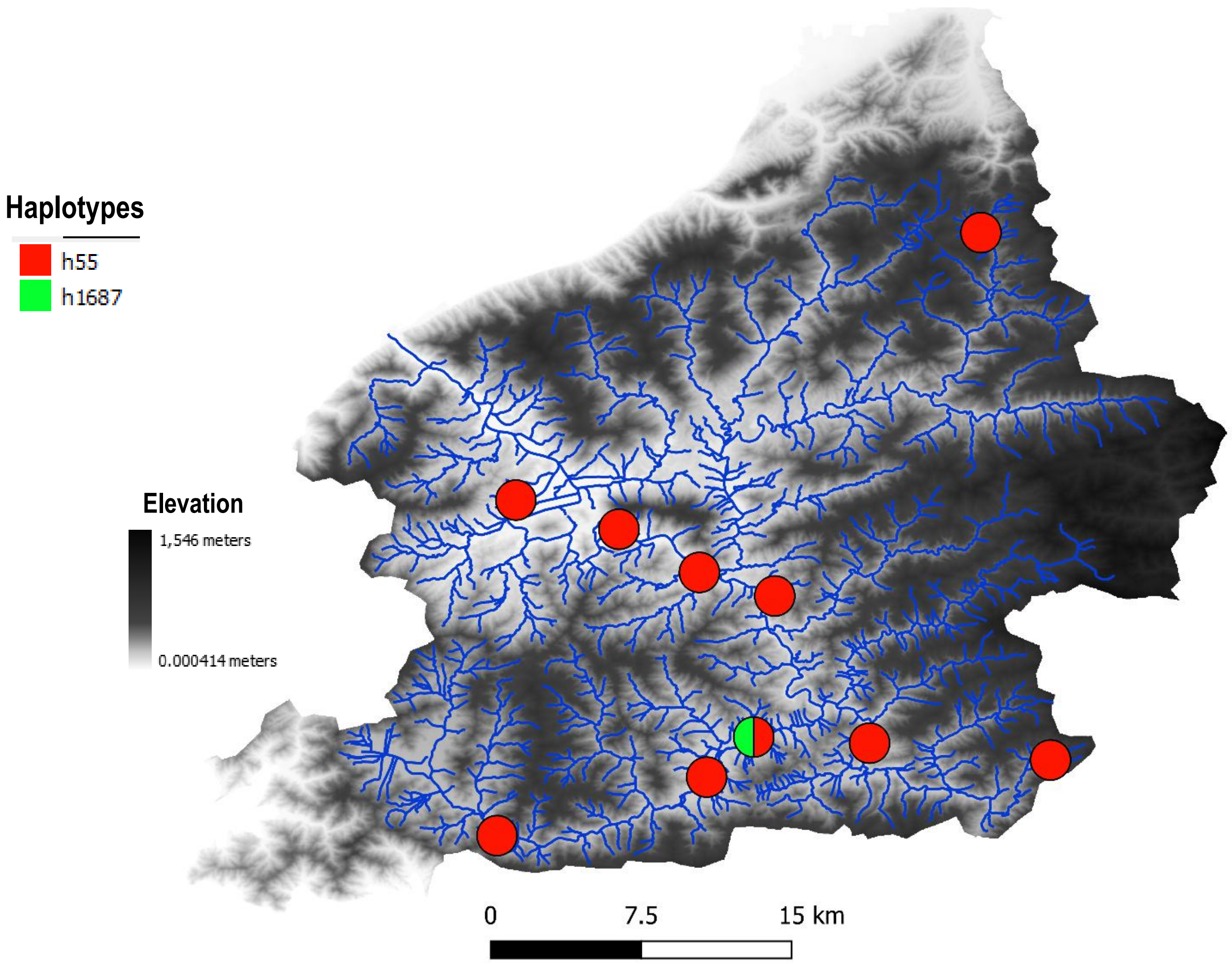


#### Haplotype network

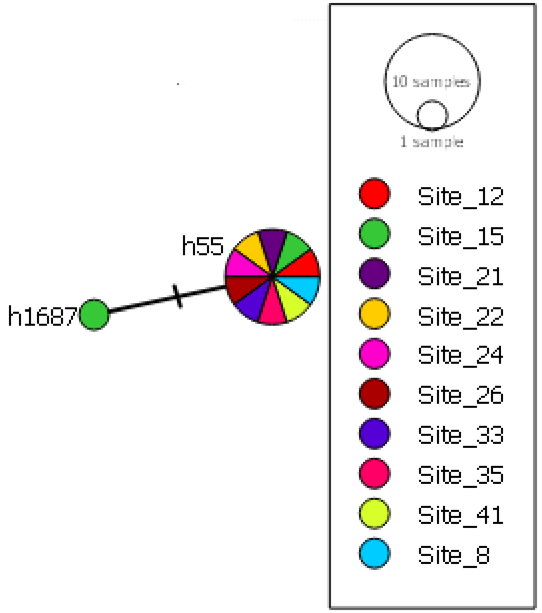

### OTU 58

#### Phylogeography

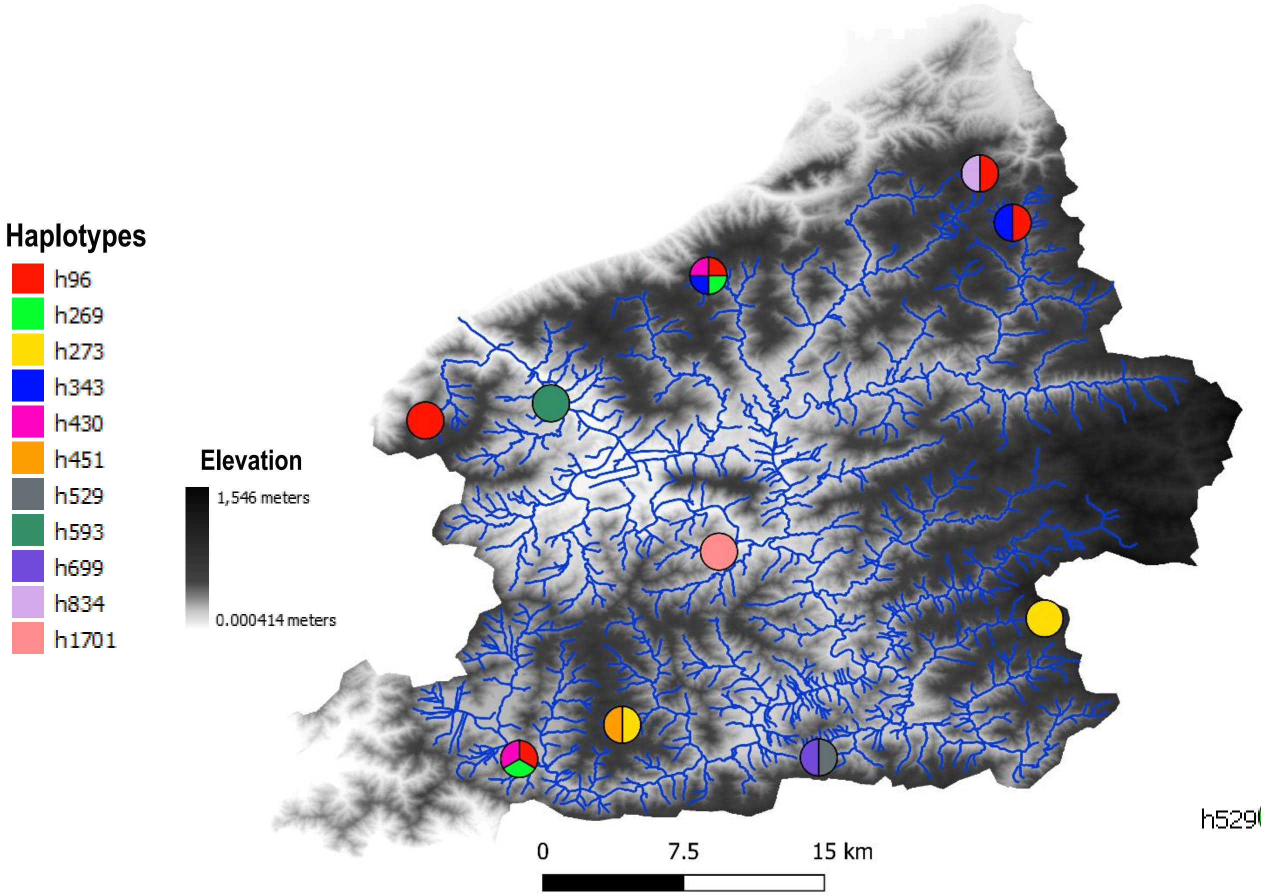


#### Haplotype network

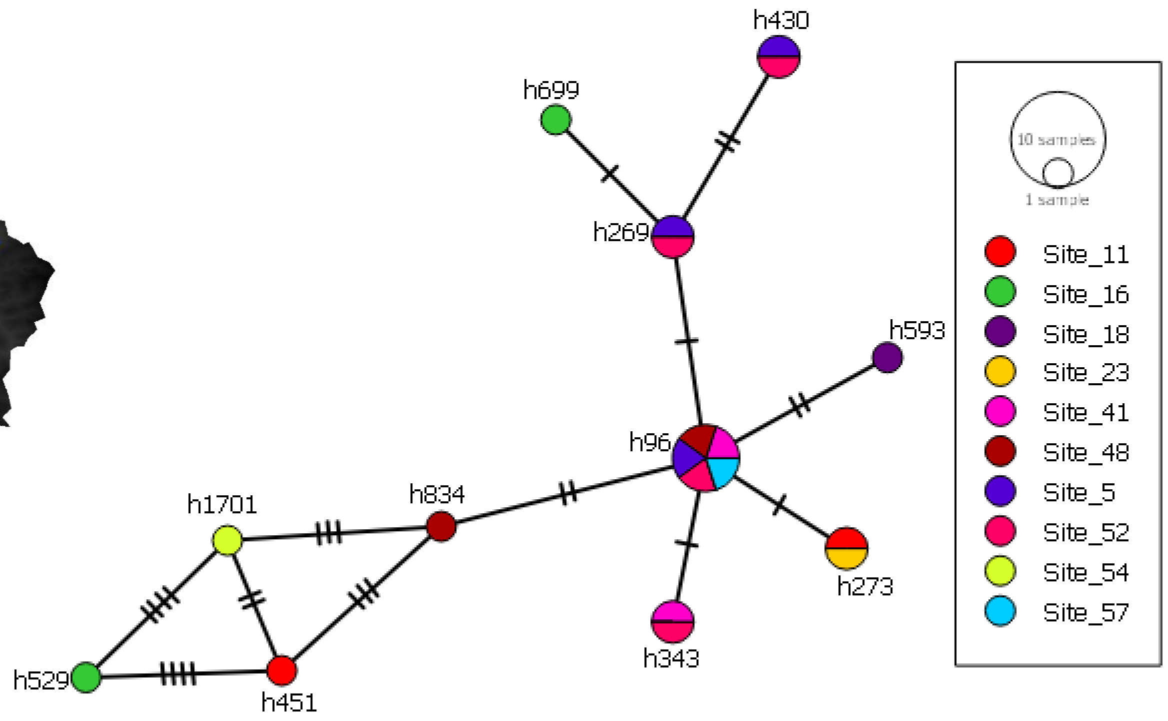

### OTU 69

#### Phylogeography

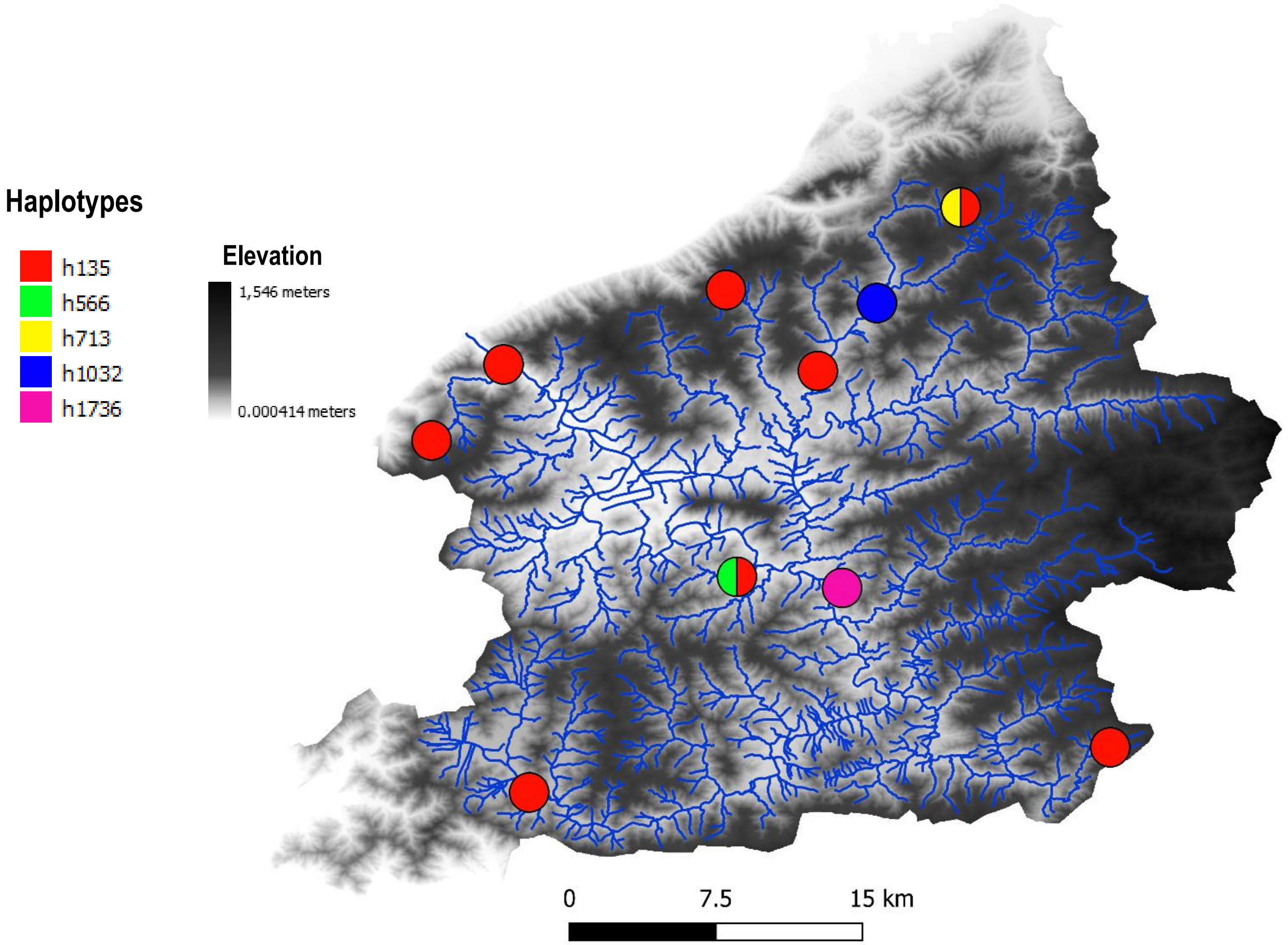


#### Haplotype network

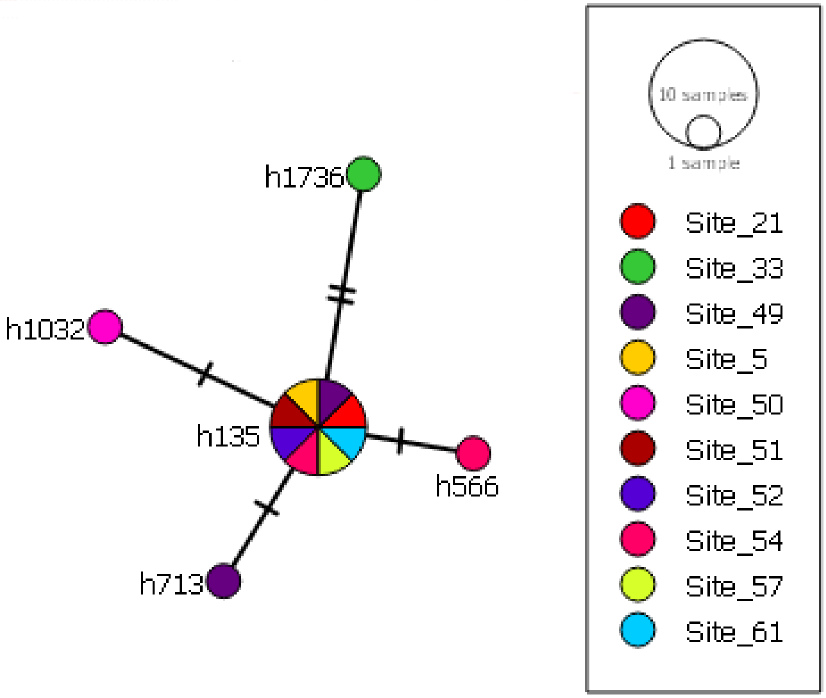

OTU 44

Phylogeography

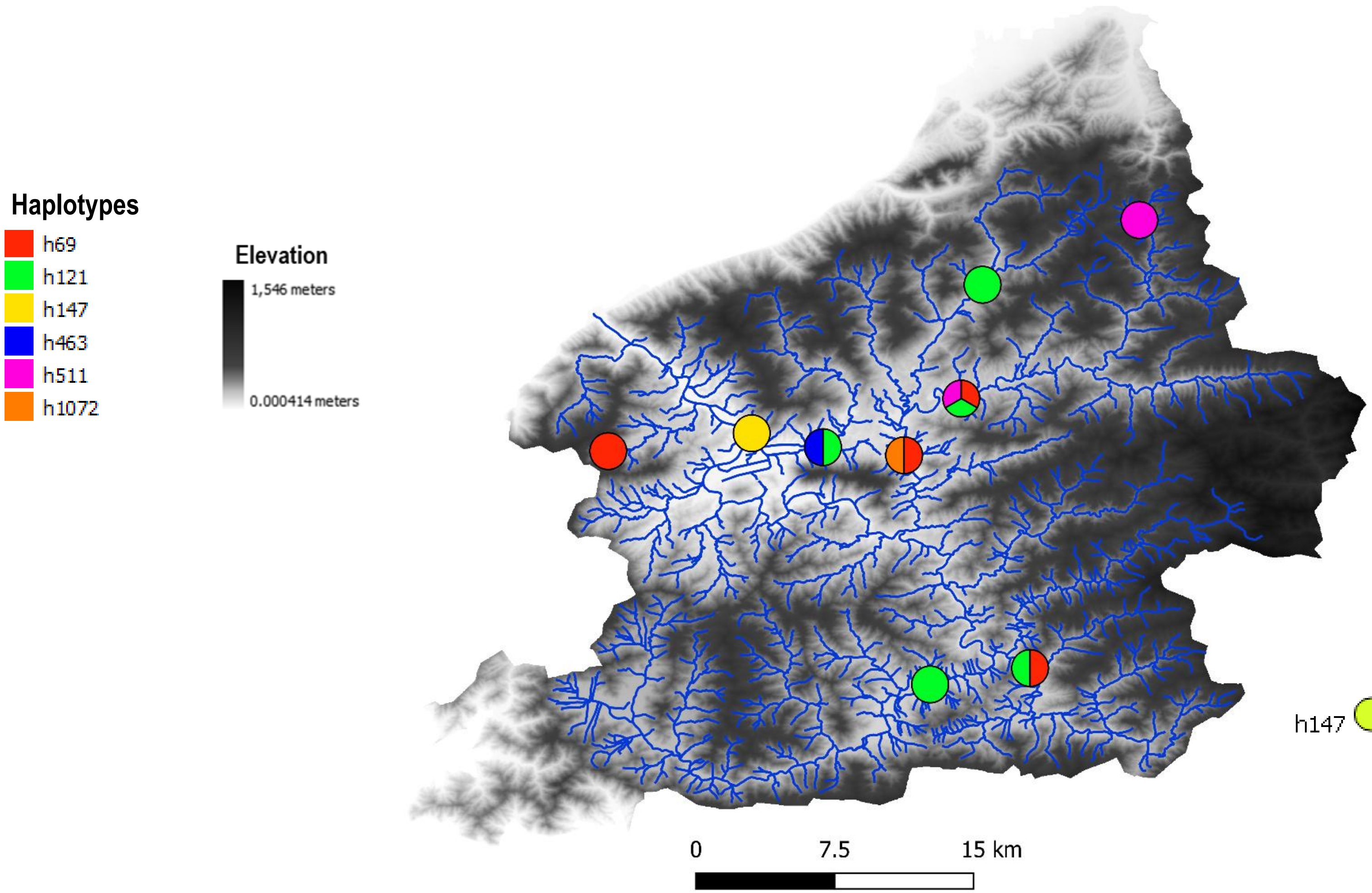


Haplotype network

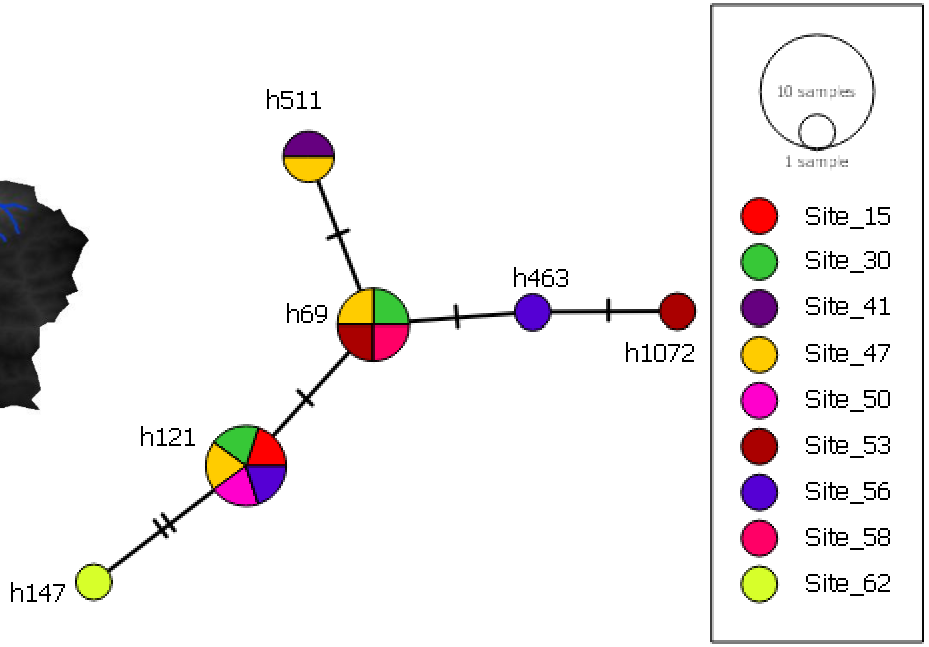

### OTU 77

#### Phylogeography

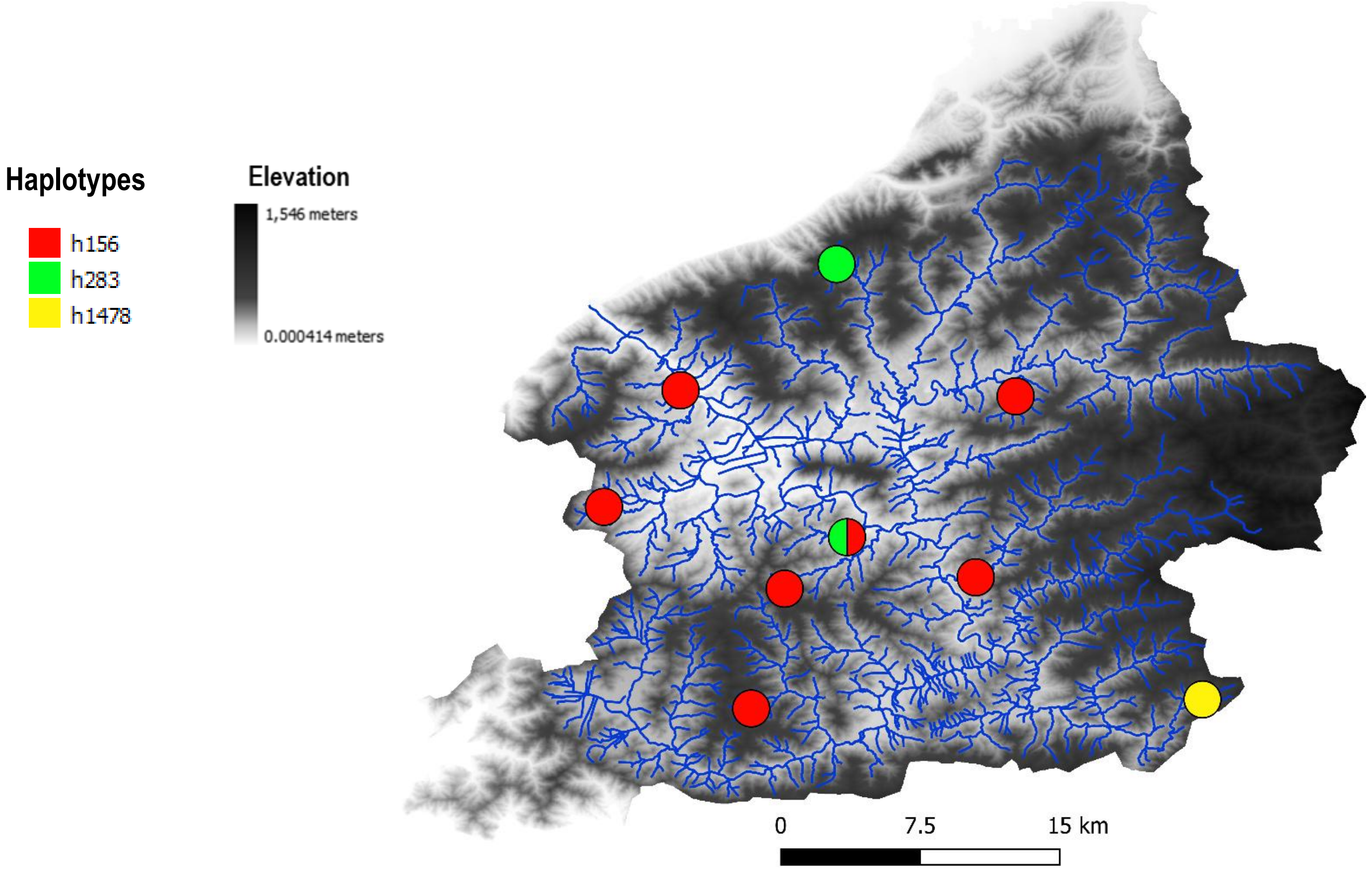


#### Haplotype network

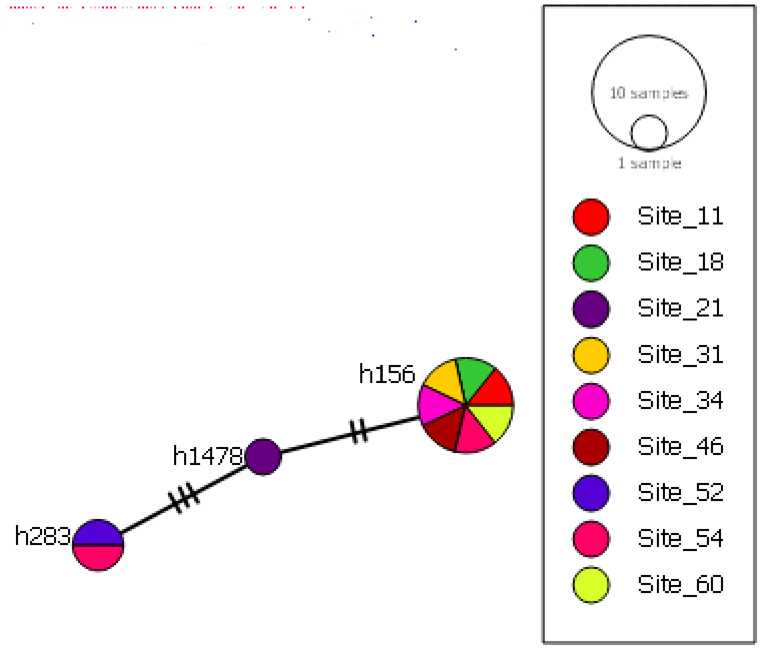

### OTU 122

#### Phylogeography

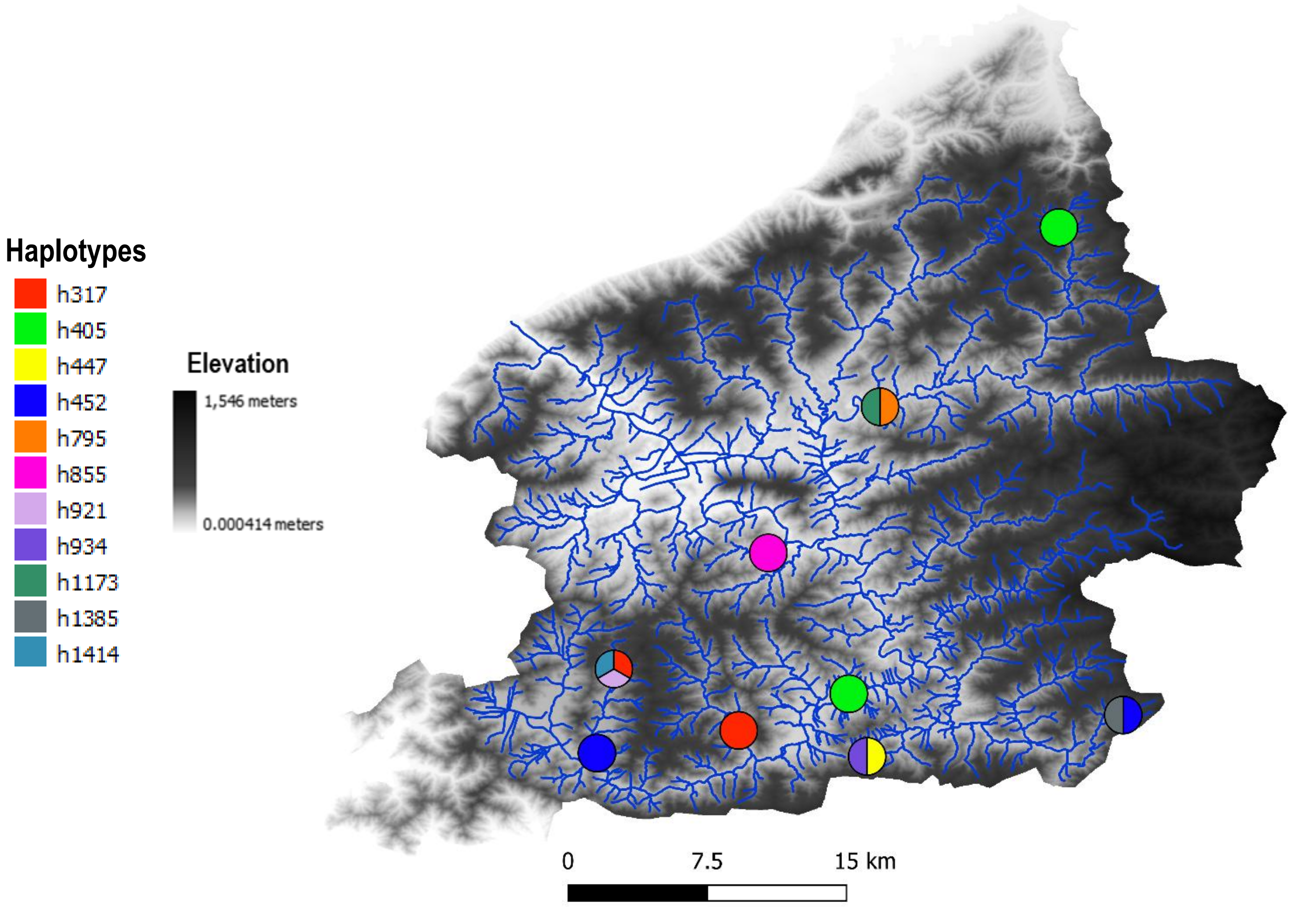


#### Haplotype network

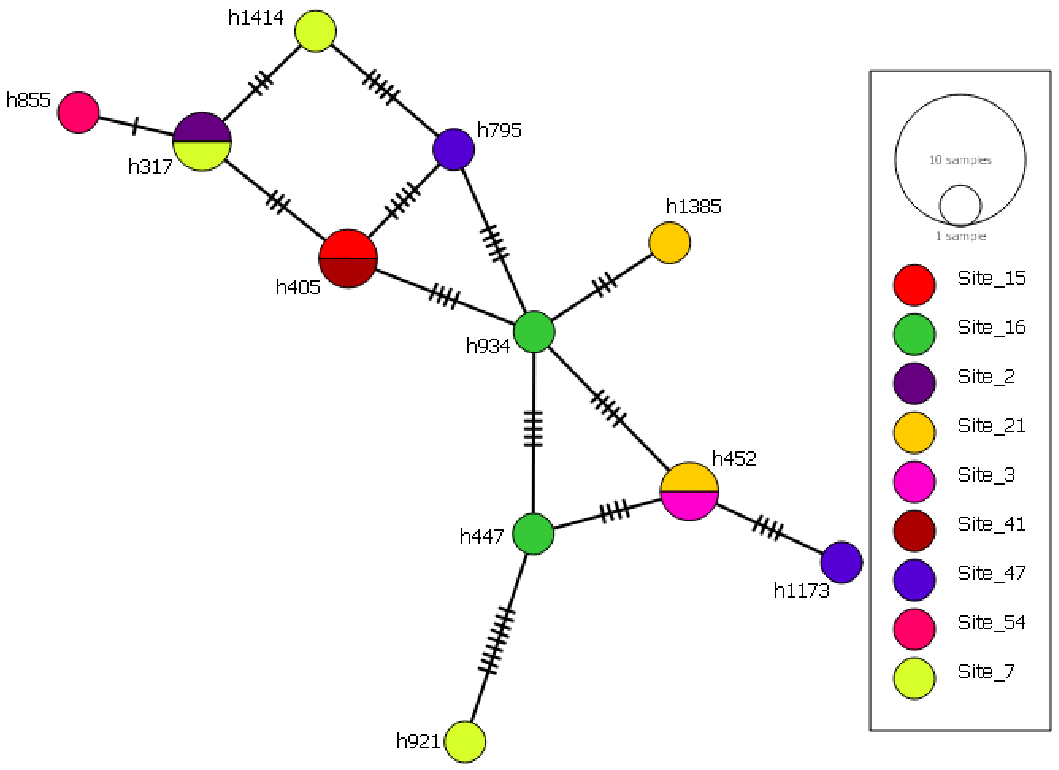

### OTU 158

#### Phylogeography

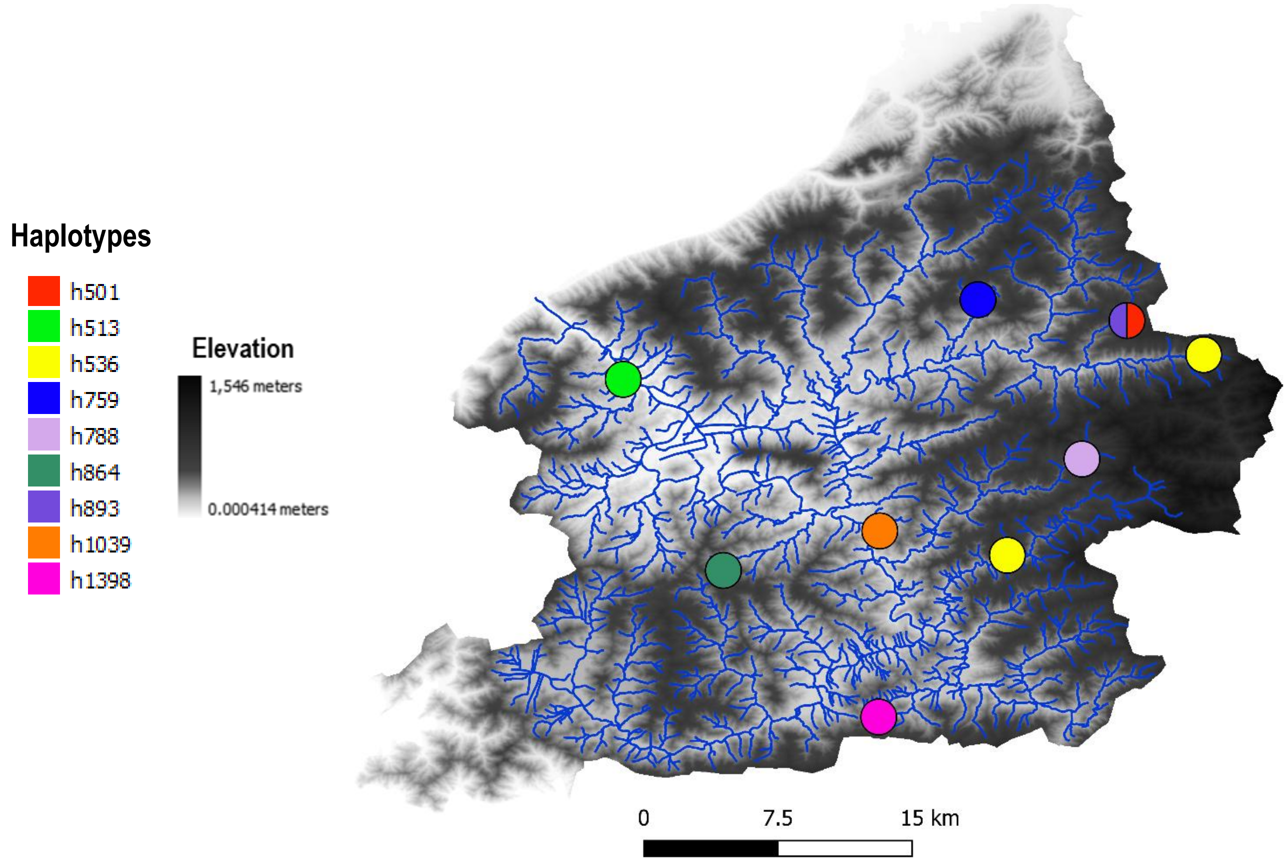


#### Haplotype network

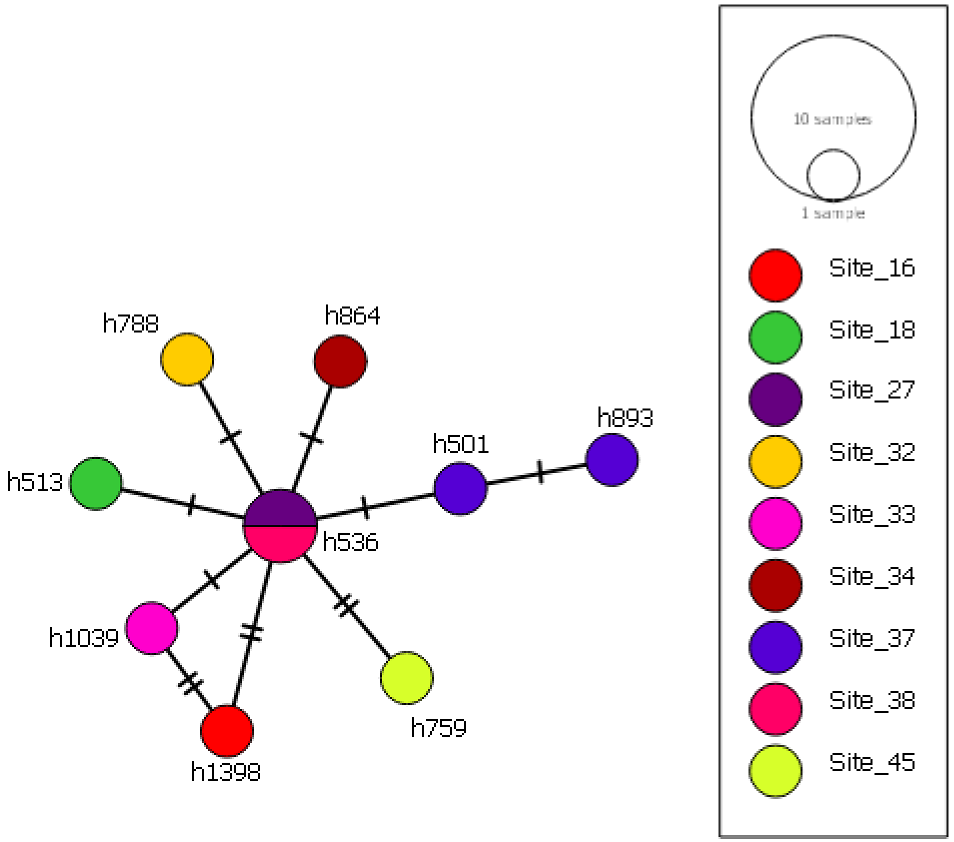

### OTU 17

#### Phylogeography

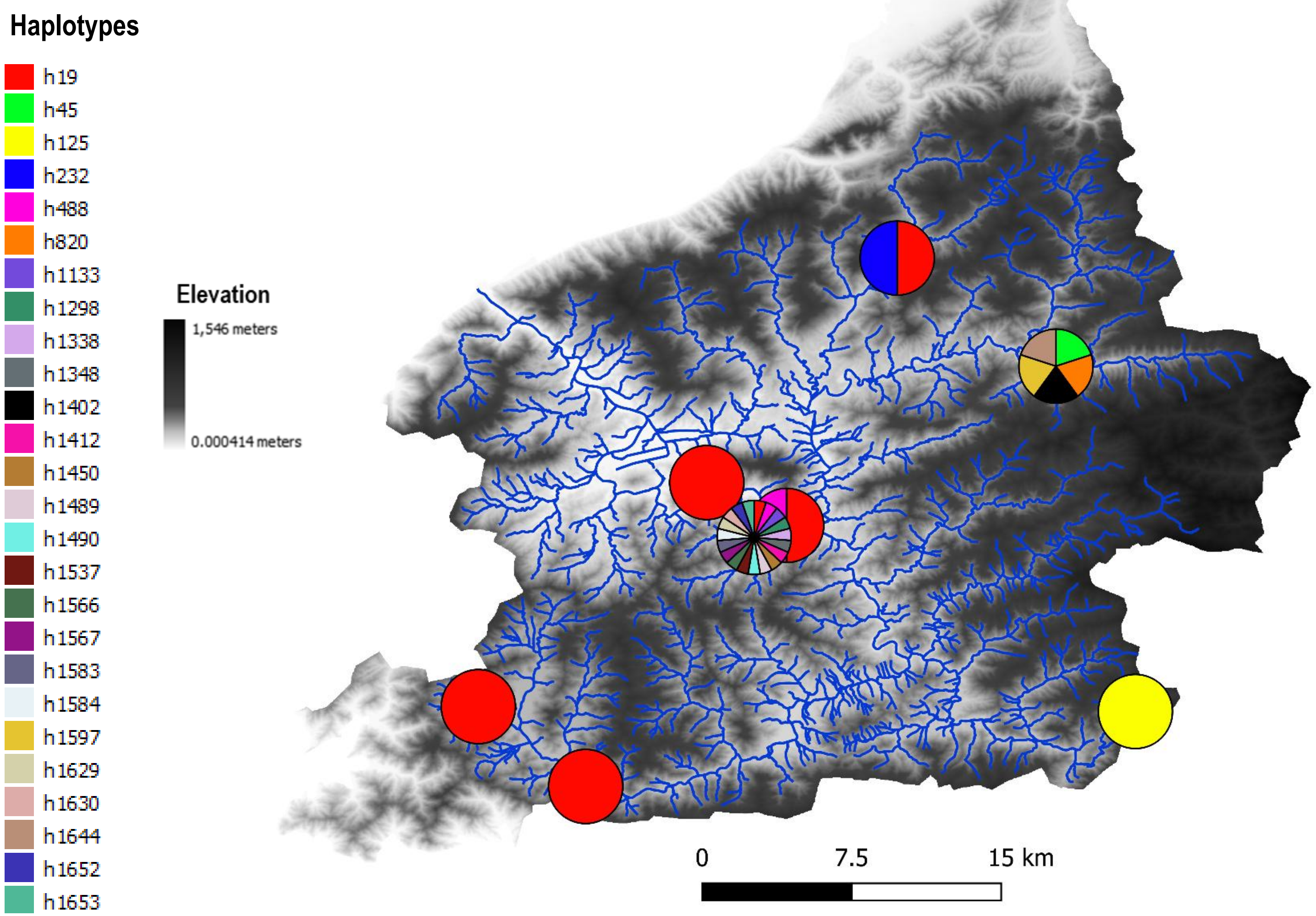


#### Haplotype network

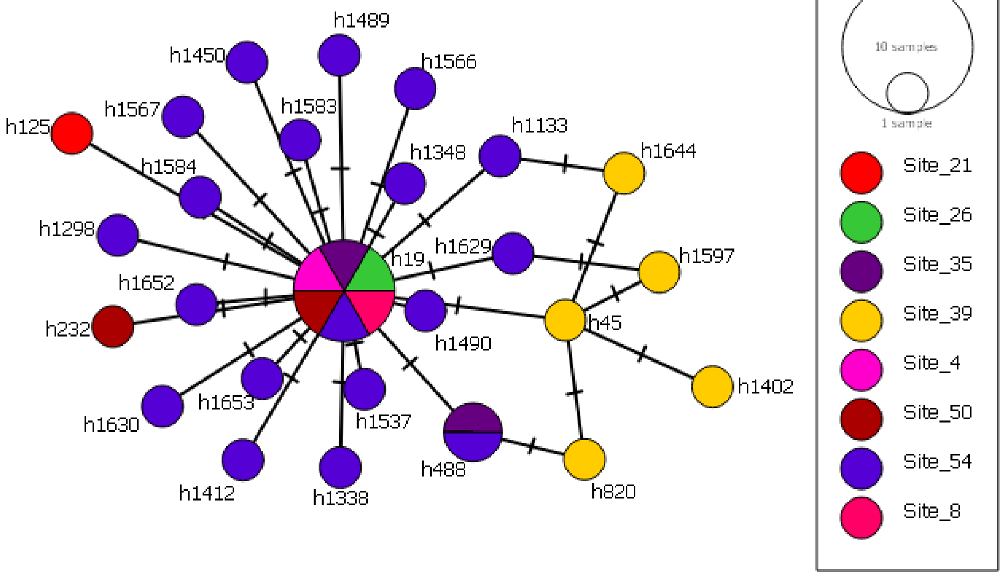

### OTU 42

#### Phylogeography

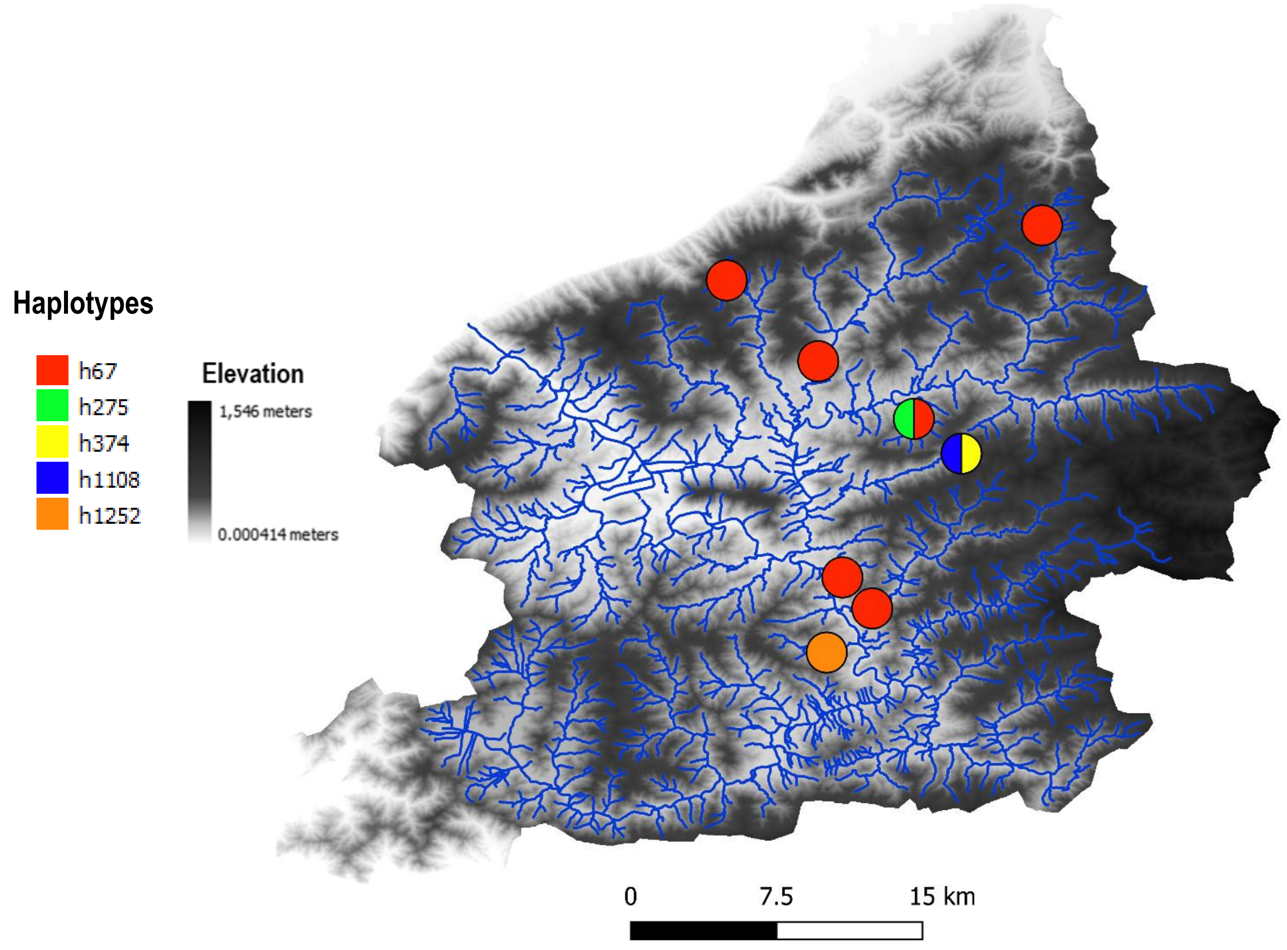


#### Haplotype network

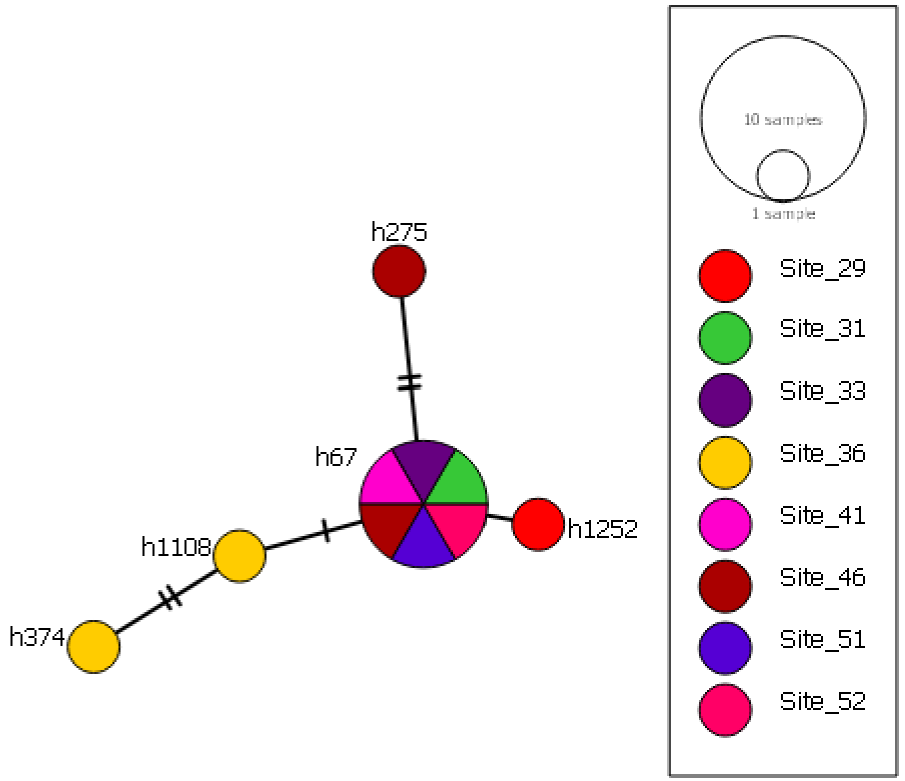

### OTU 55

#### Phylogeography

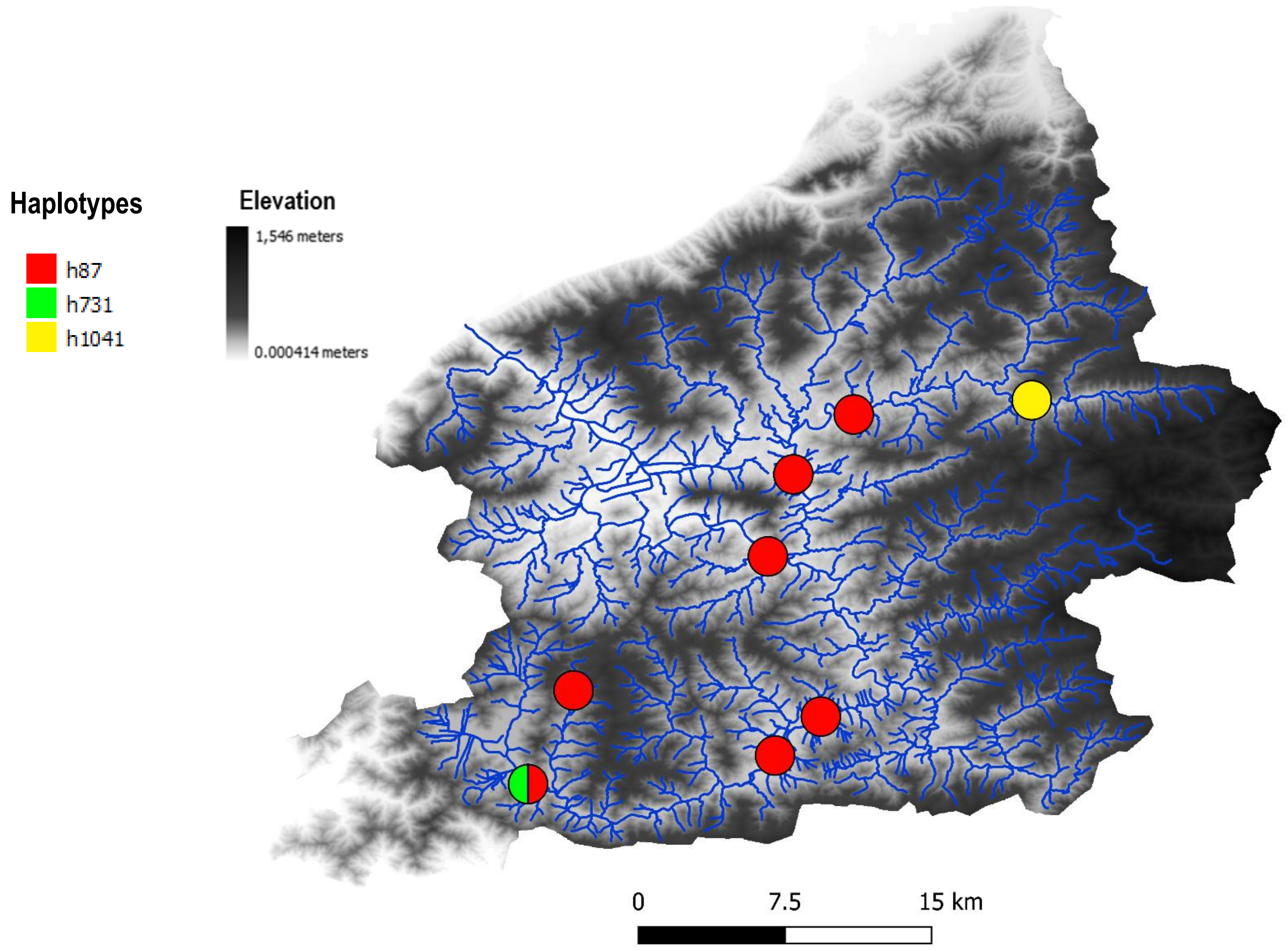


#### Haplotype network

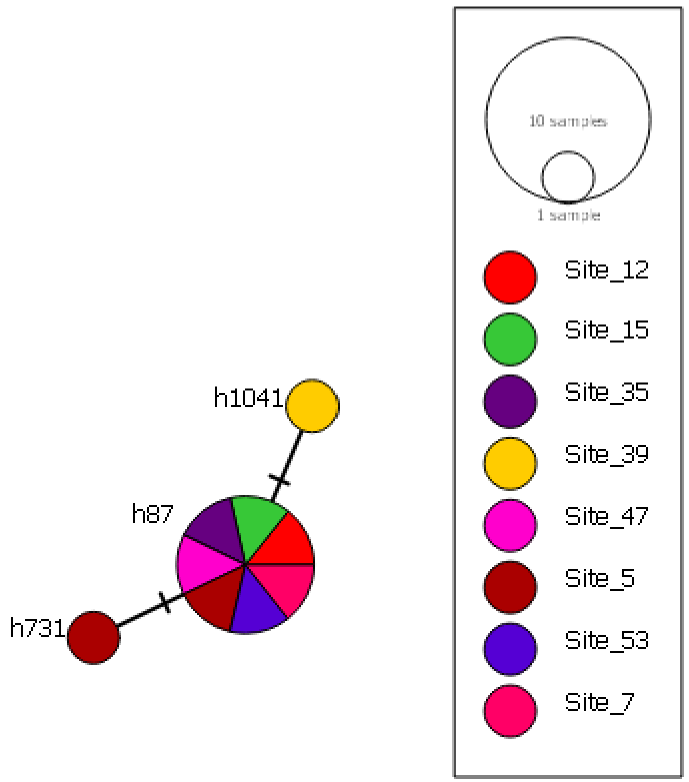

### OTU 56

#### Phylogeography

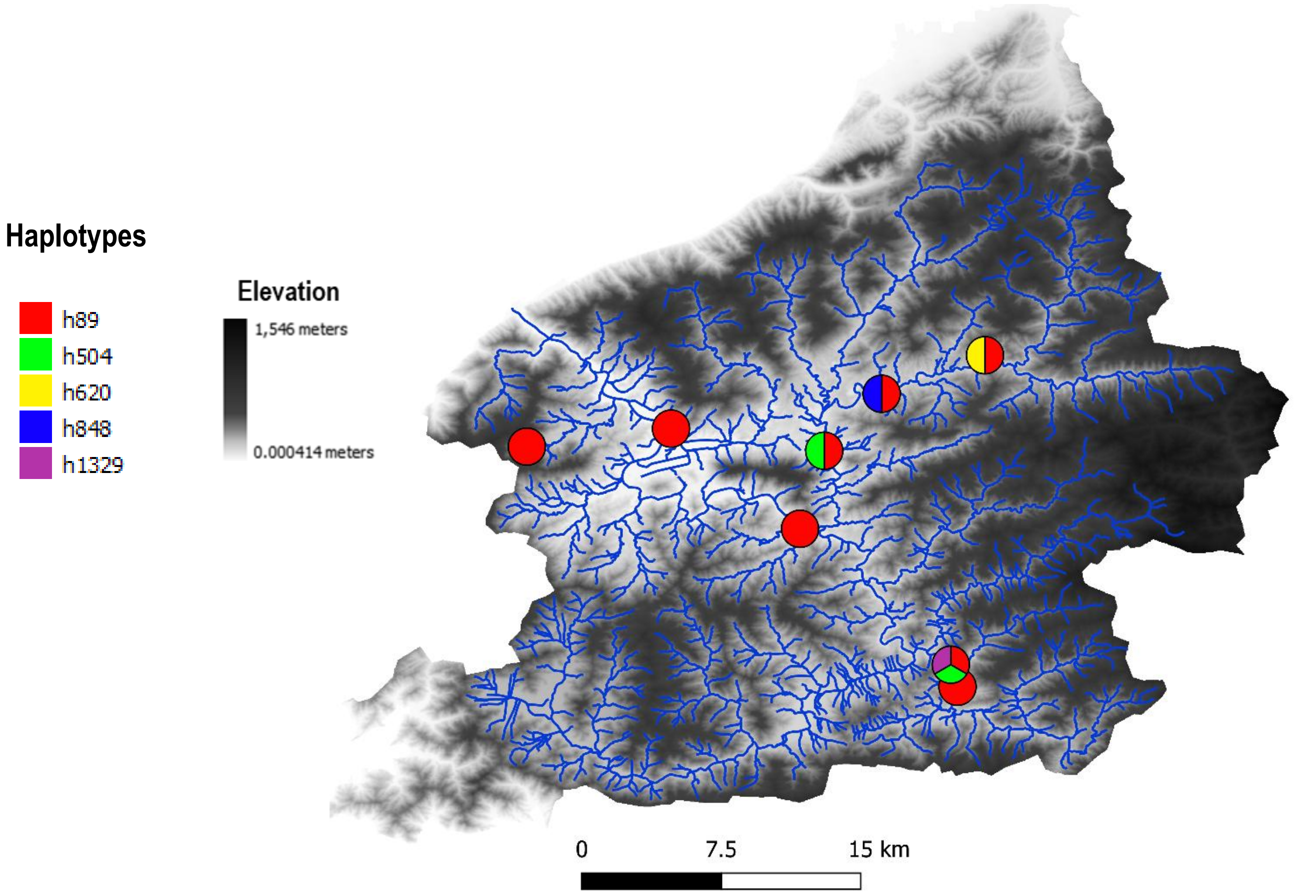


#### Haplotype network

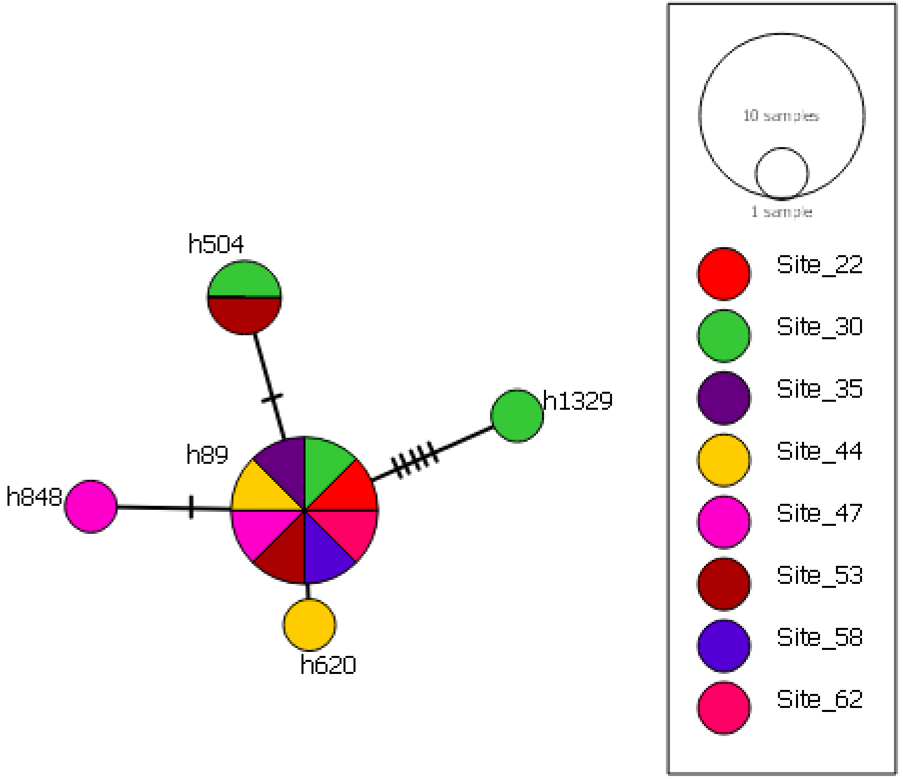

OTU 118

Phylogeography

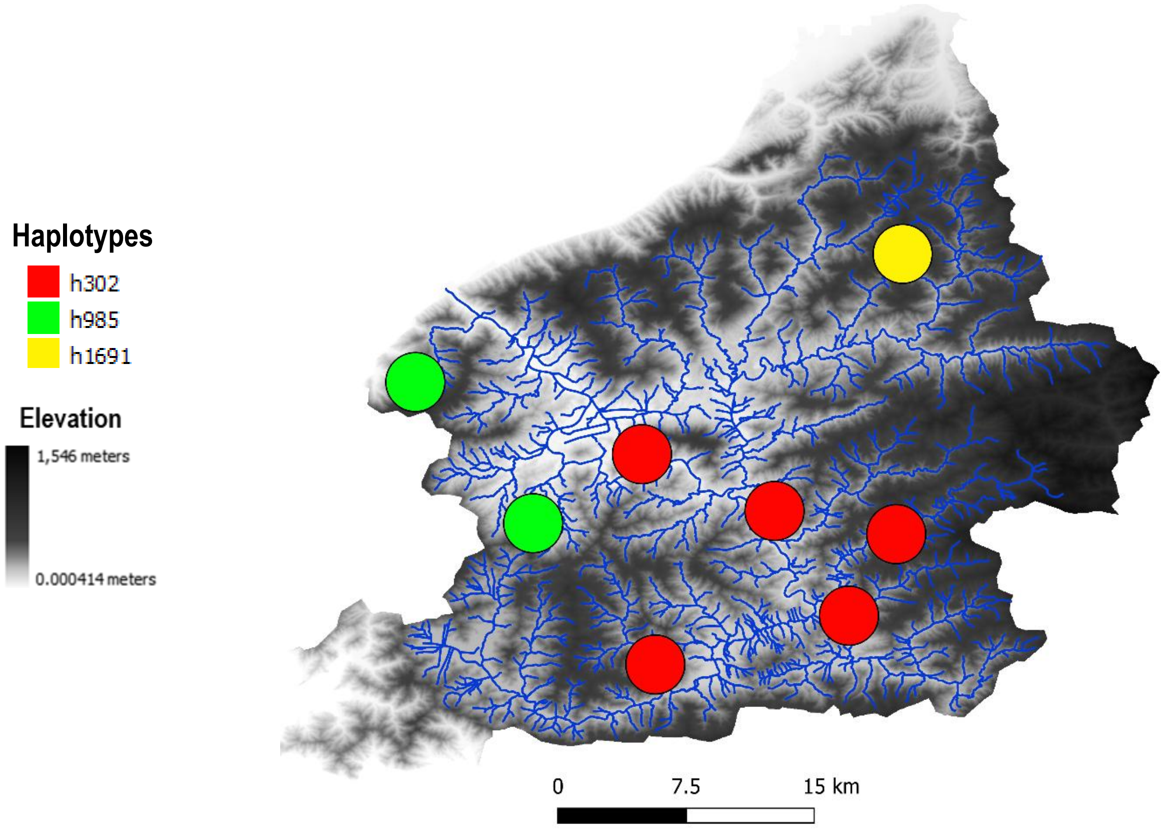


Haplotype network

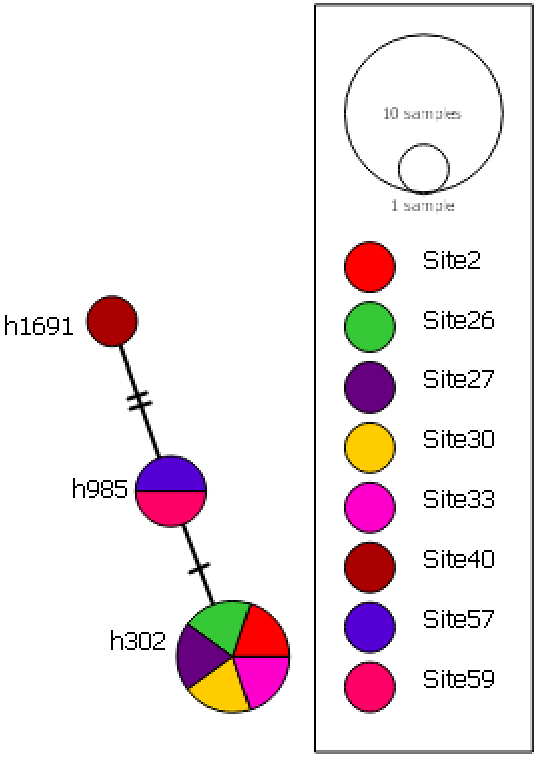

### OTU 53

#### Phylogeography

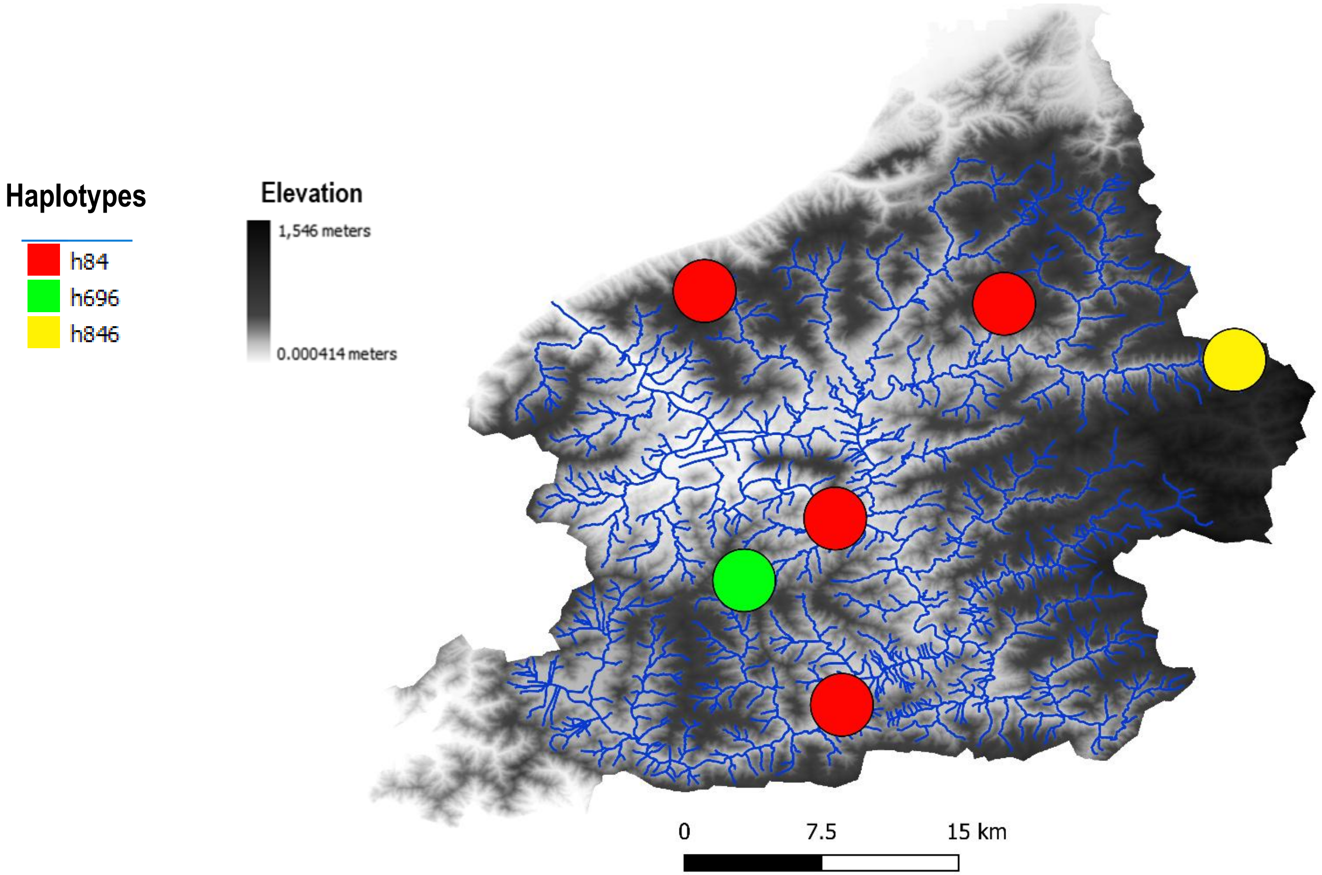


#### Haplotype network

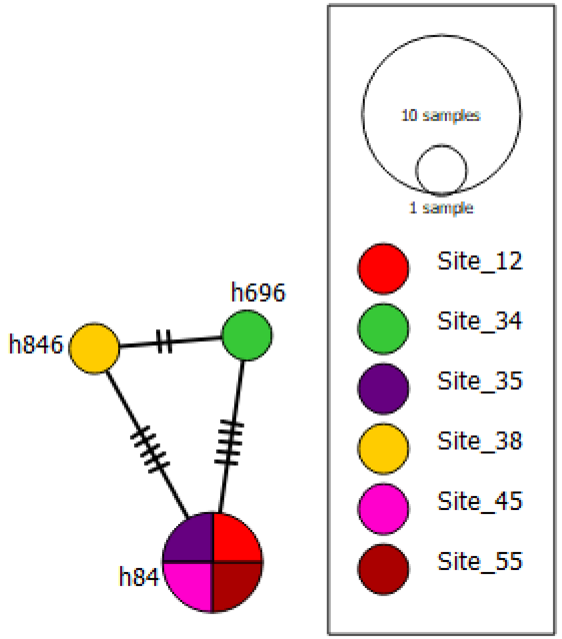

### OTU 16

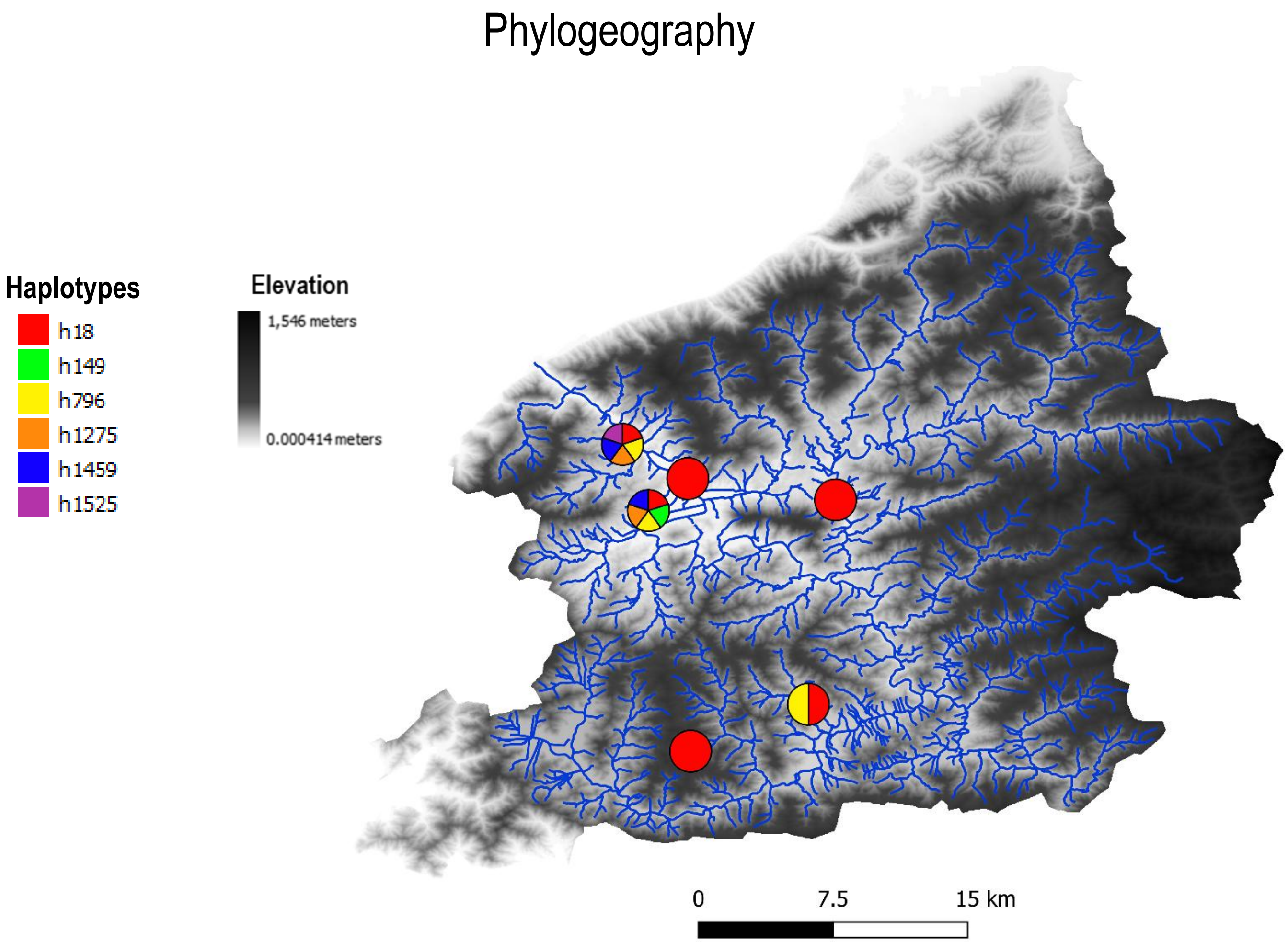
Phylogeography
Haplotypes
h18
h149
h796
h1275
h1459
h1525
Elevation
1,546 meters
0.000414 meters
0
7.5
15 km

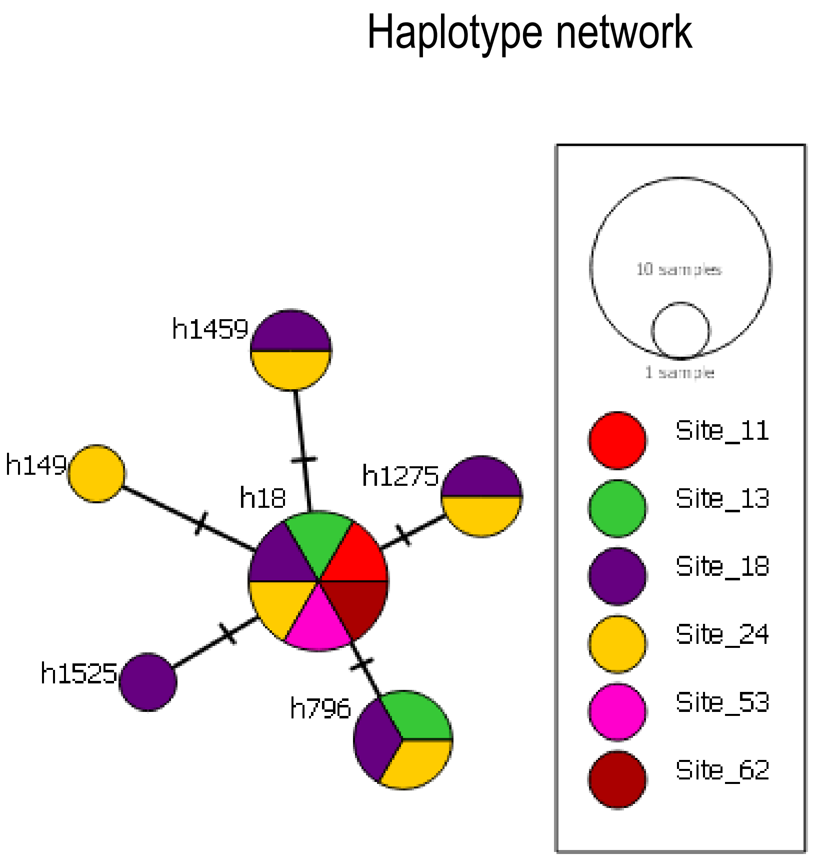
Haplotype network
h1459
h149
h18
h1275
h1525
h796
10 samples
1 sample
Site_11
Site_13
Site_18
Site_24
Site_53
Site_62

### OTU 65

#### Phylogeography

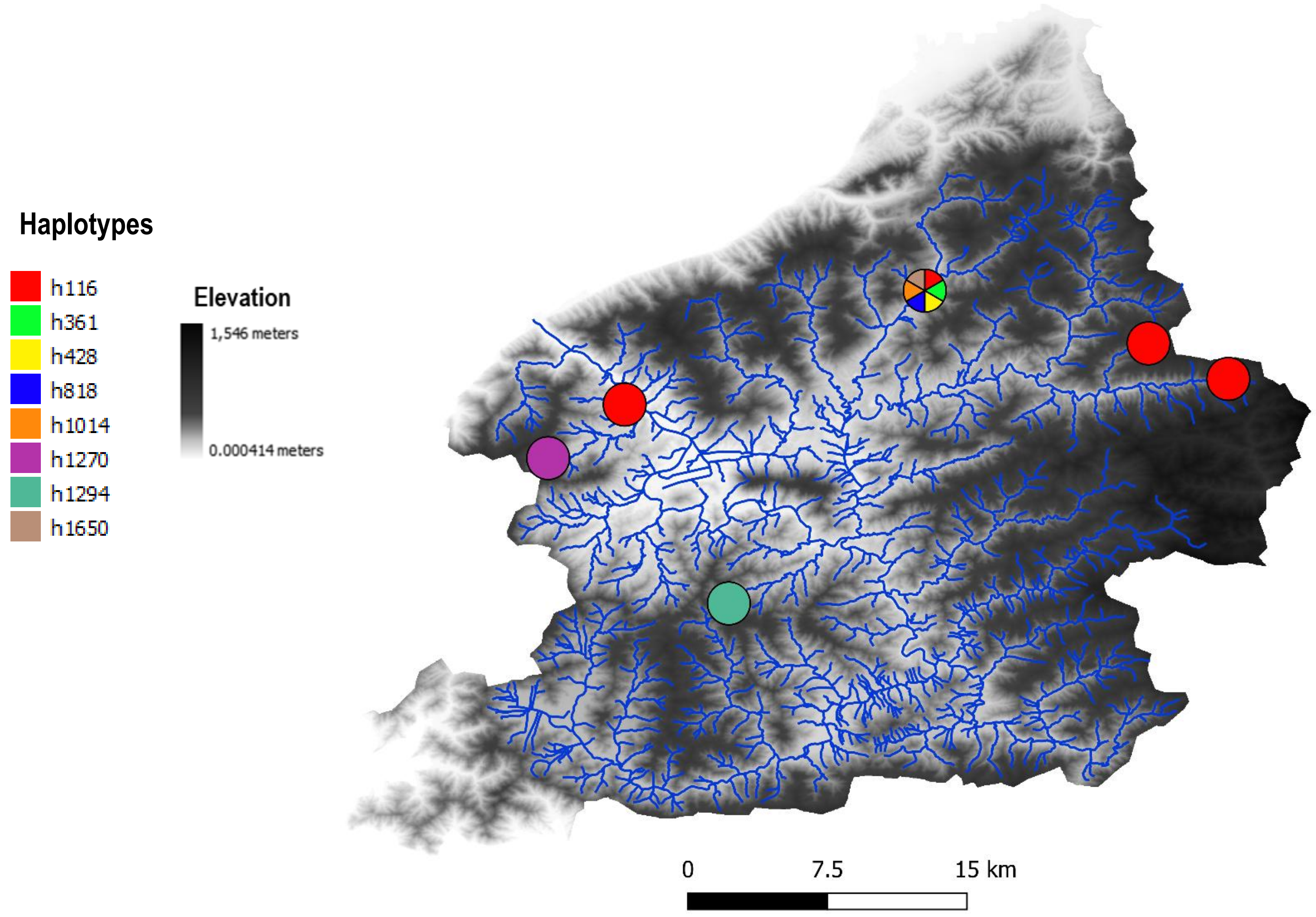


#### Haplotype network

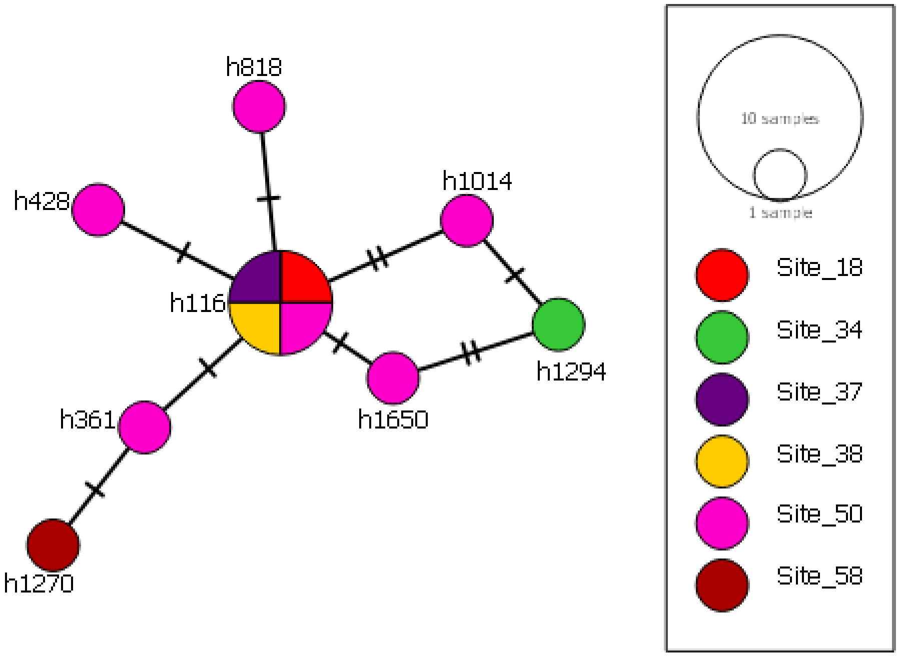

### OTU 87

#### Phylogeography

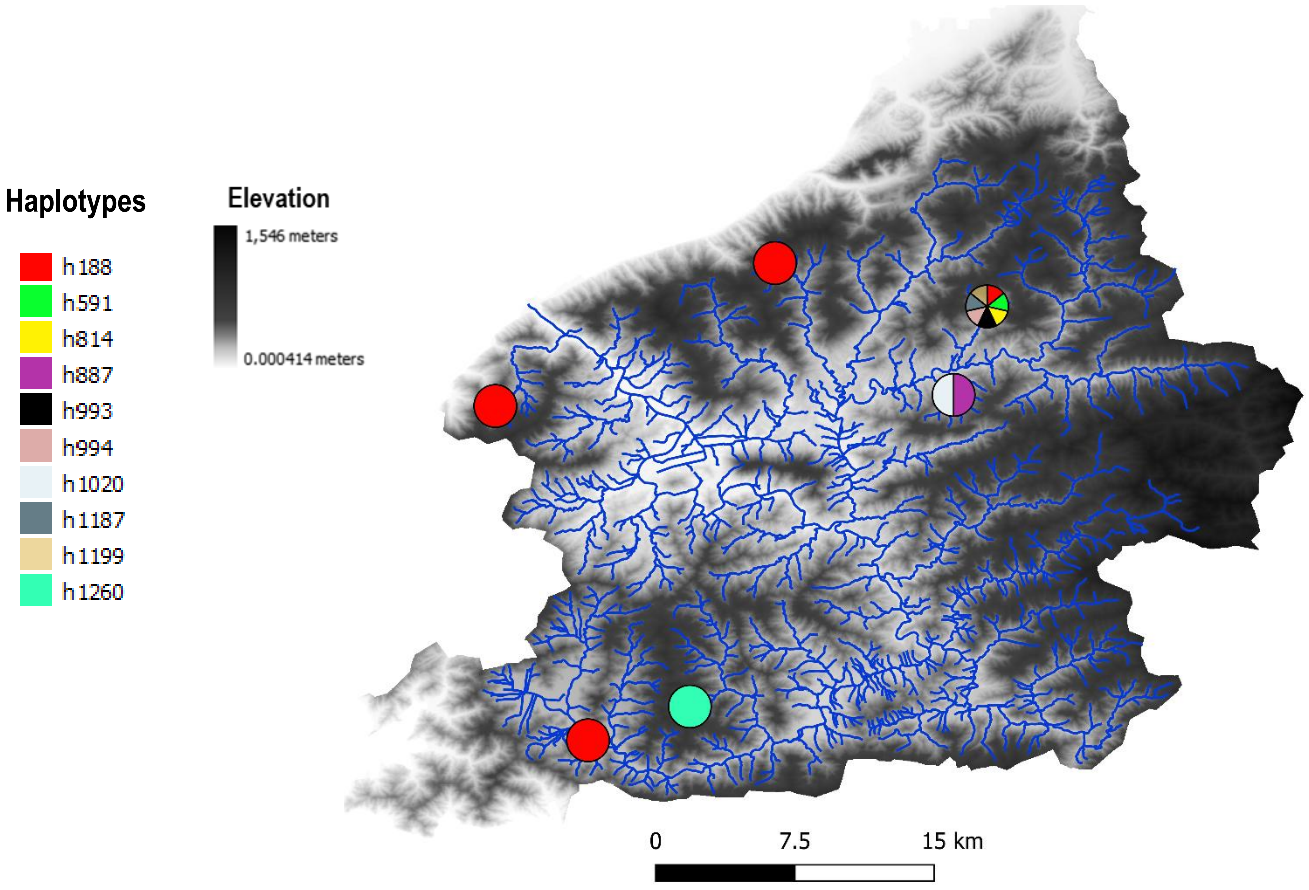


#### Haplotype network

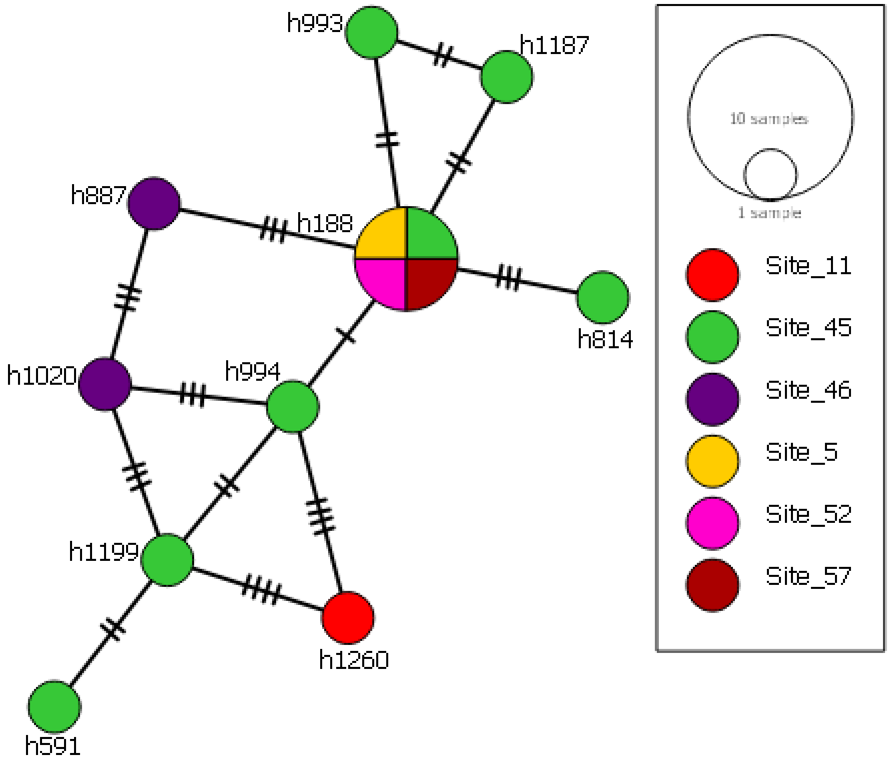

### OUT 79

#### Phylogeography

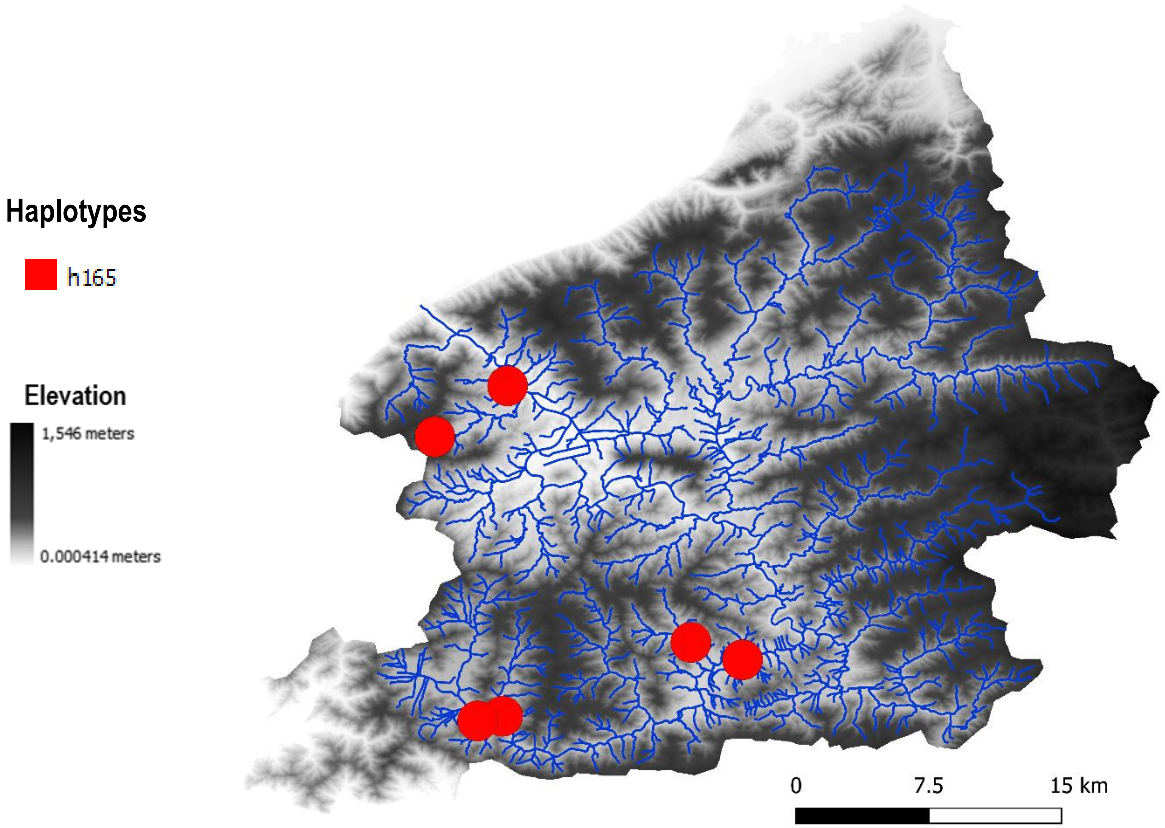


#### Haplotype network

OTU90

#### Phylogeography

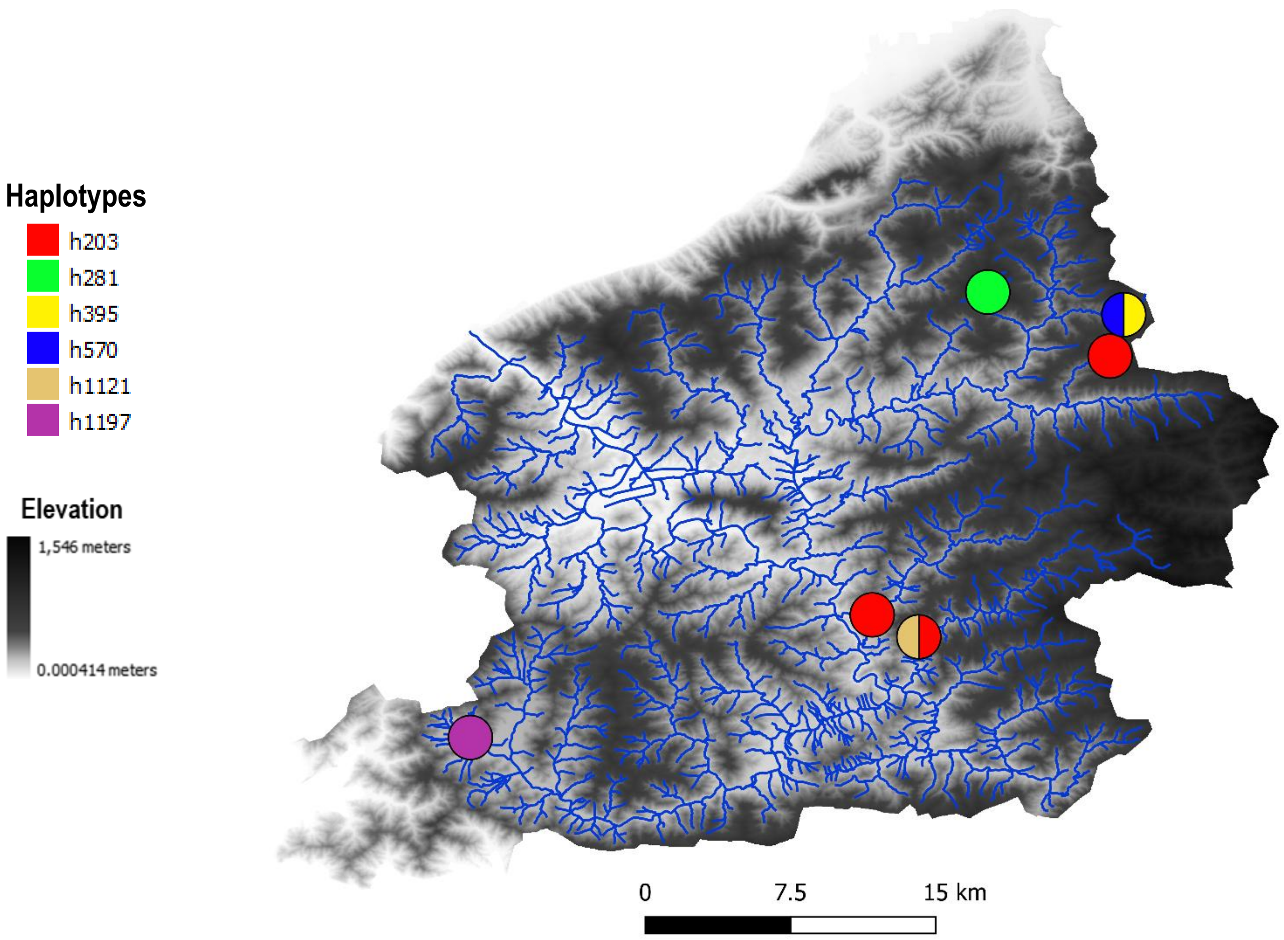


#### Haplotype network

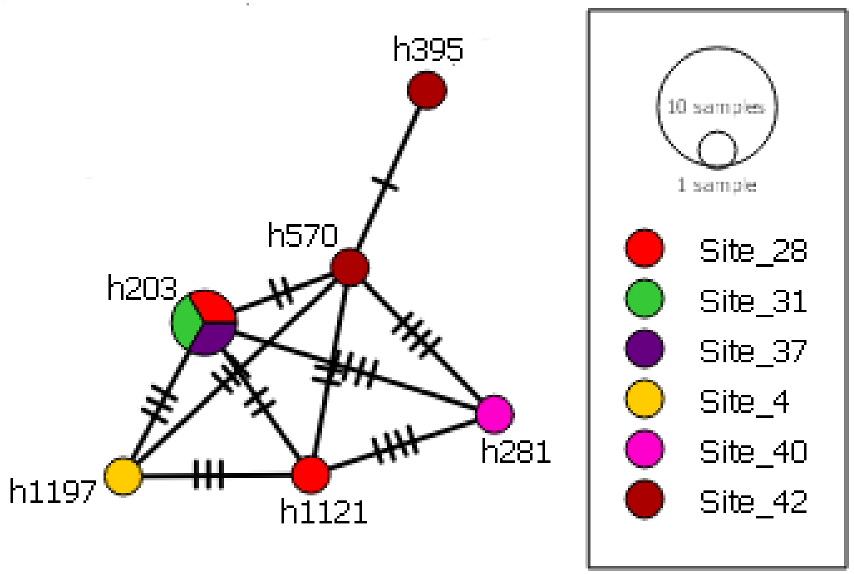

OTU113

#### Phylogeography

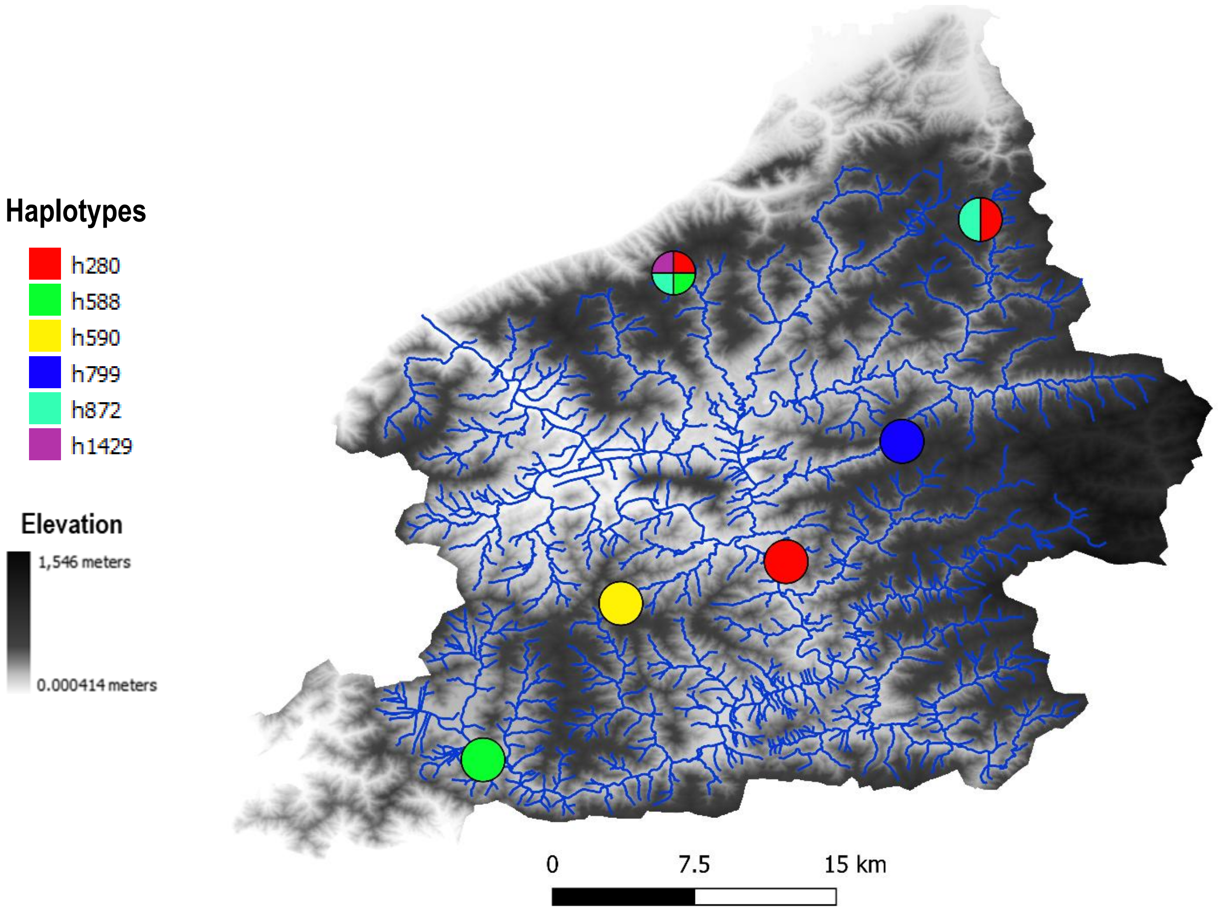


#### Haplotype network

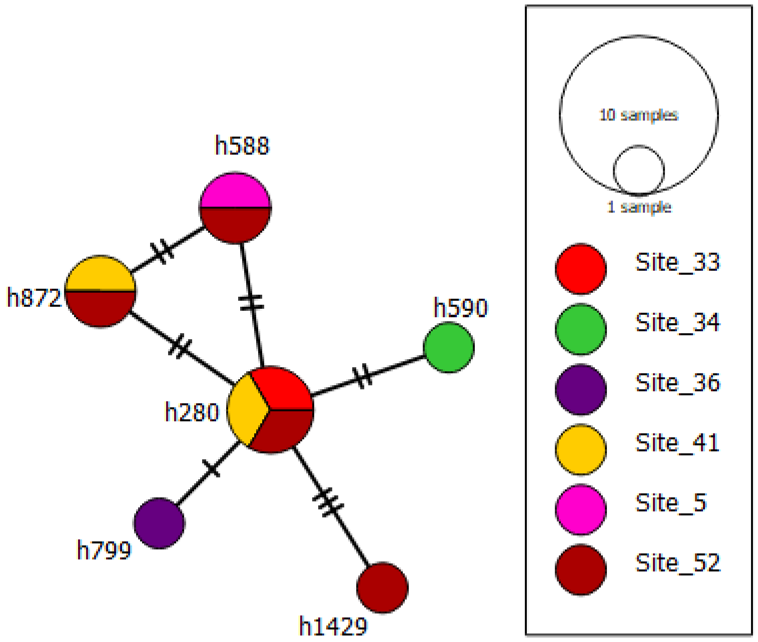

### OTU 198

#### Phylogeography

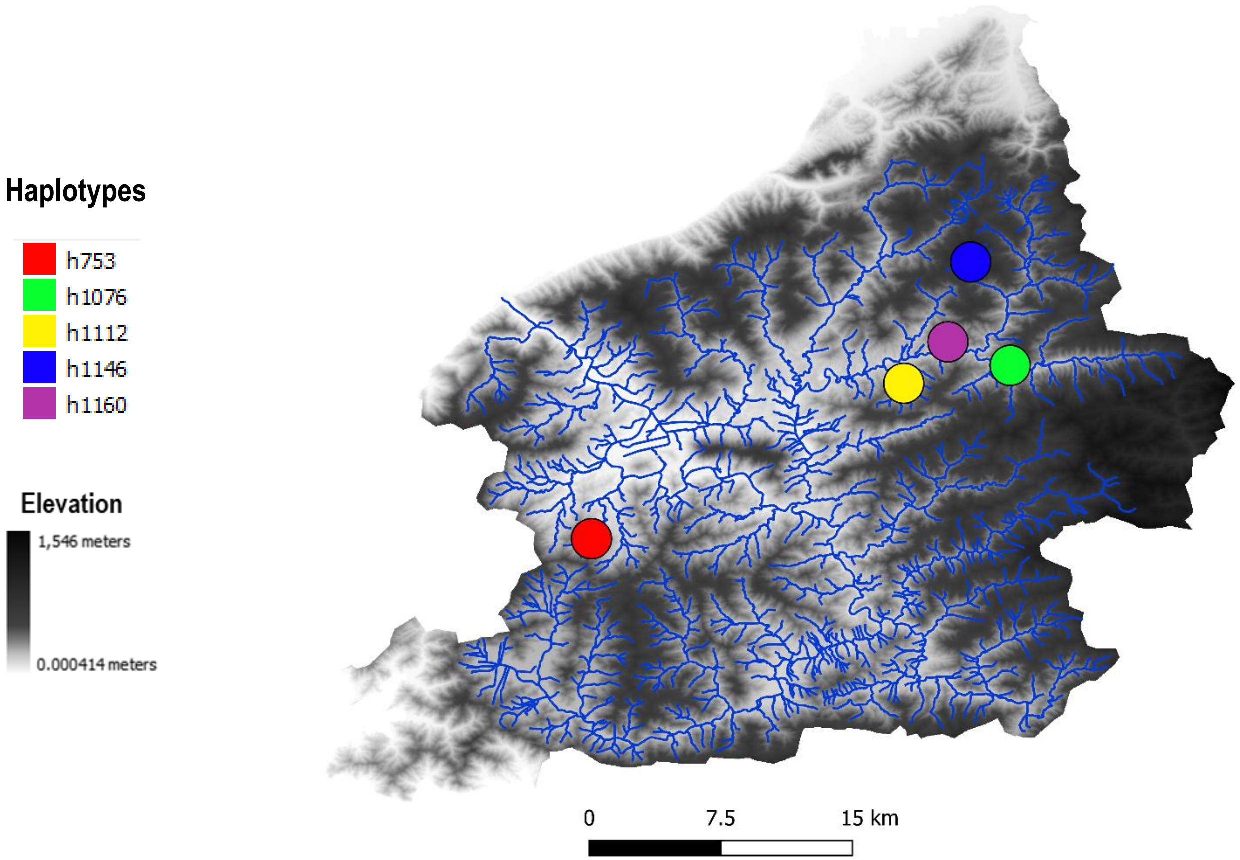


#### Haplotype network

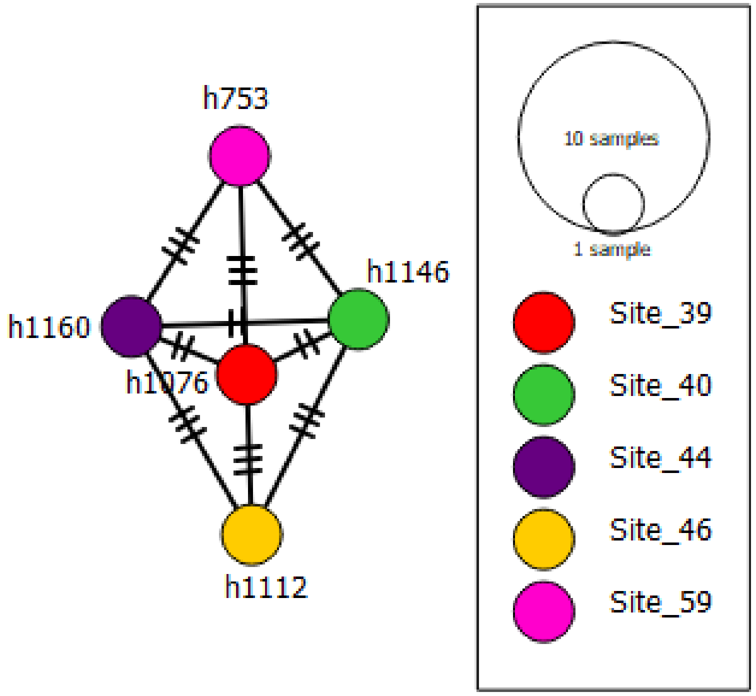

### OTU 52

#### Phylogeography

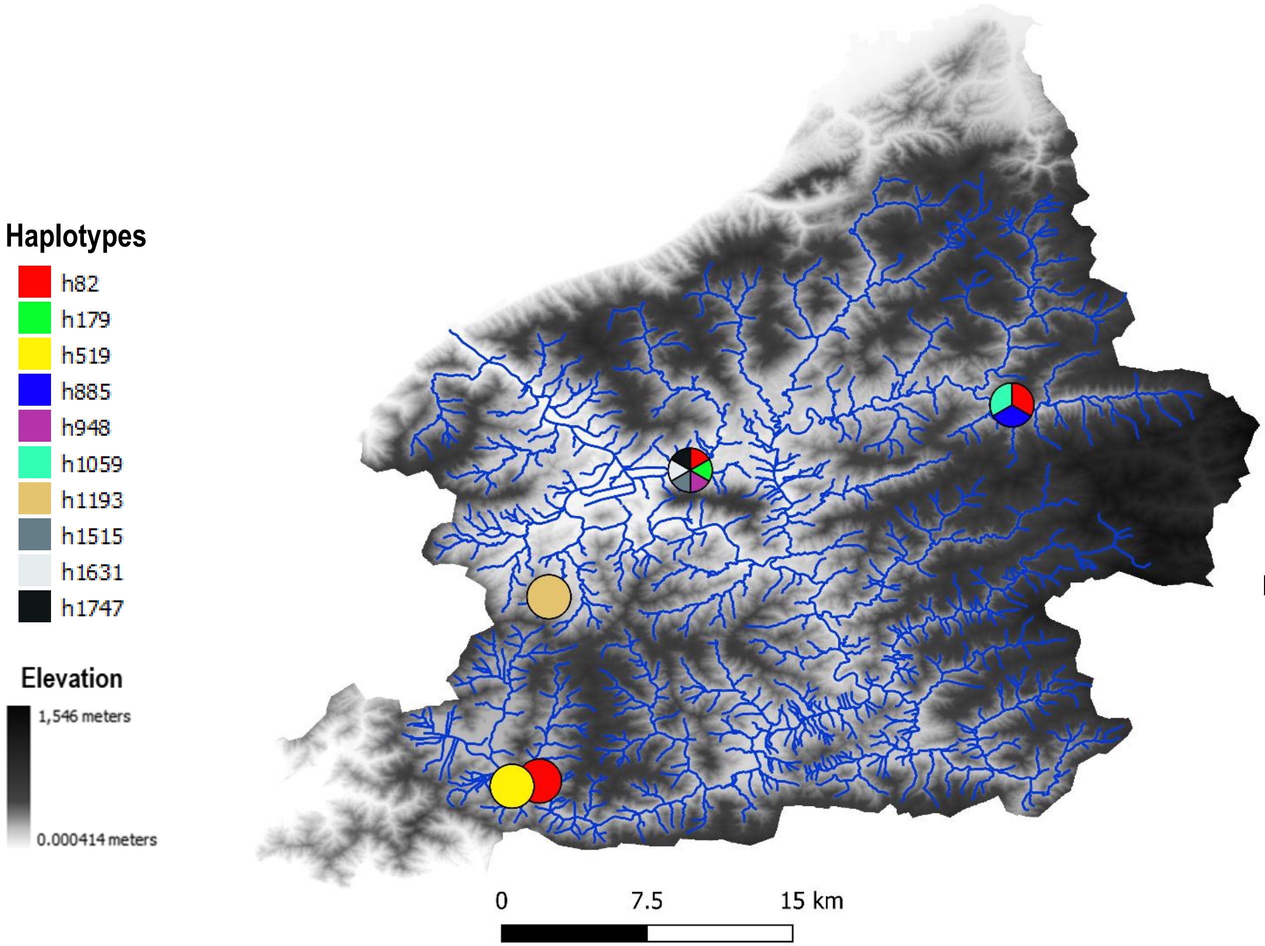


#### Haplotype network

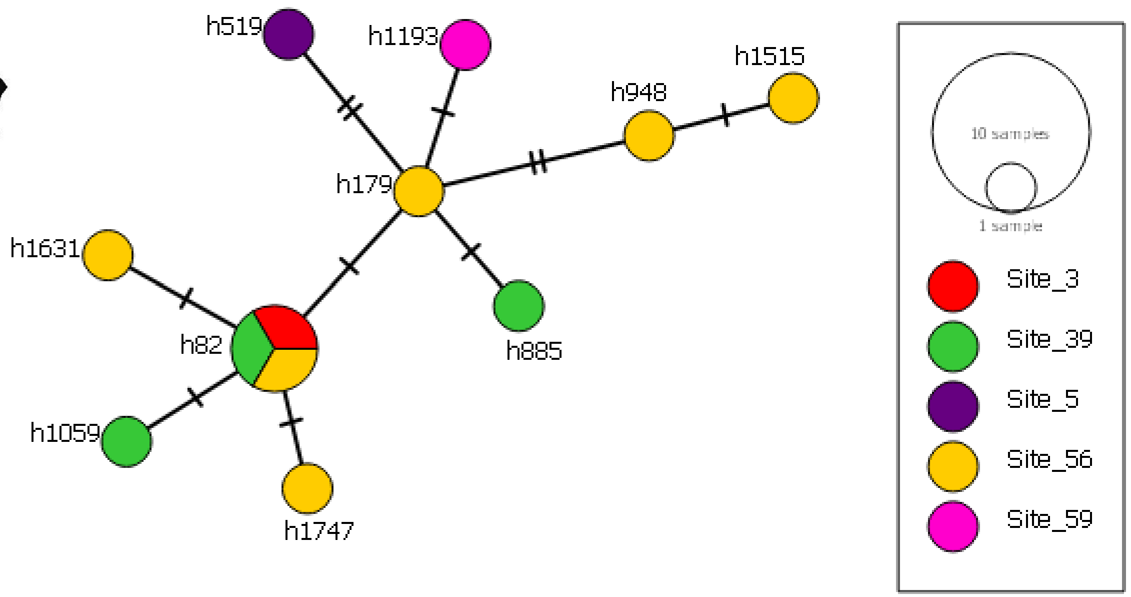

### OTU 54

#### Phylogeography

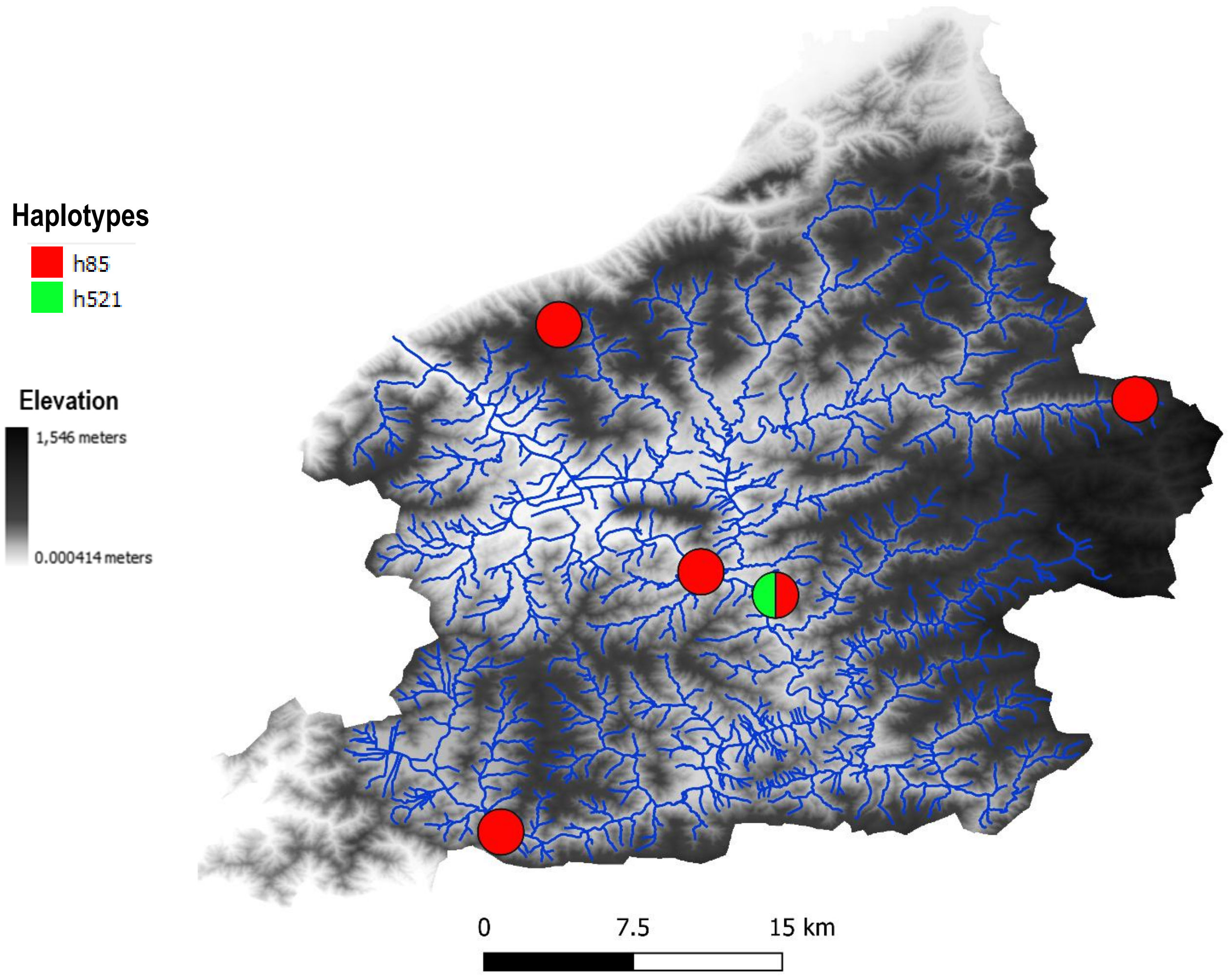


#### Haplotype network

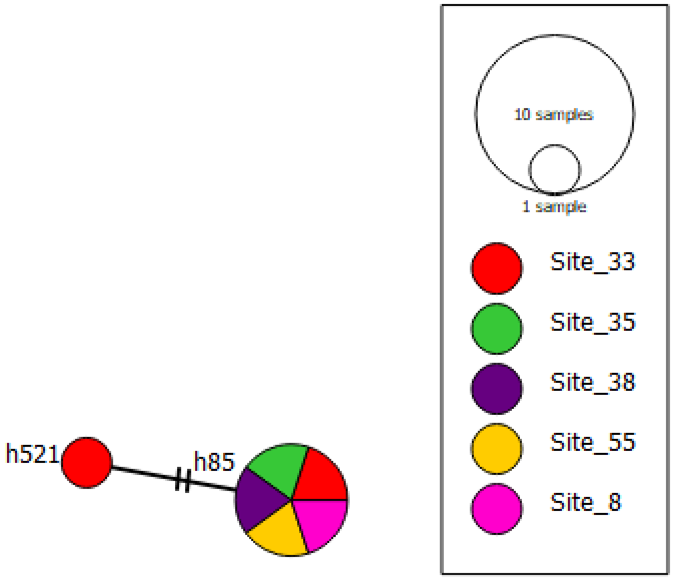

### OTU 70

#### Phylogeography

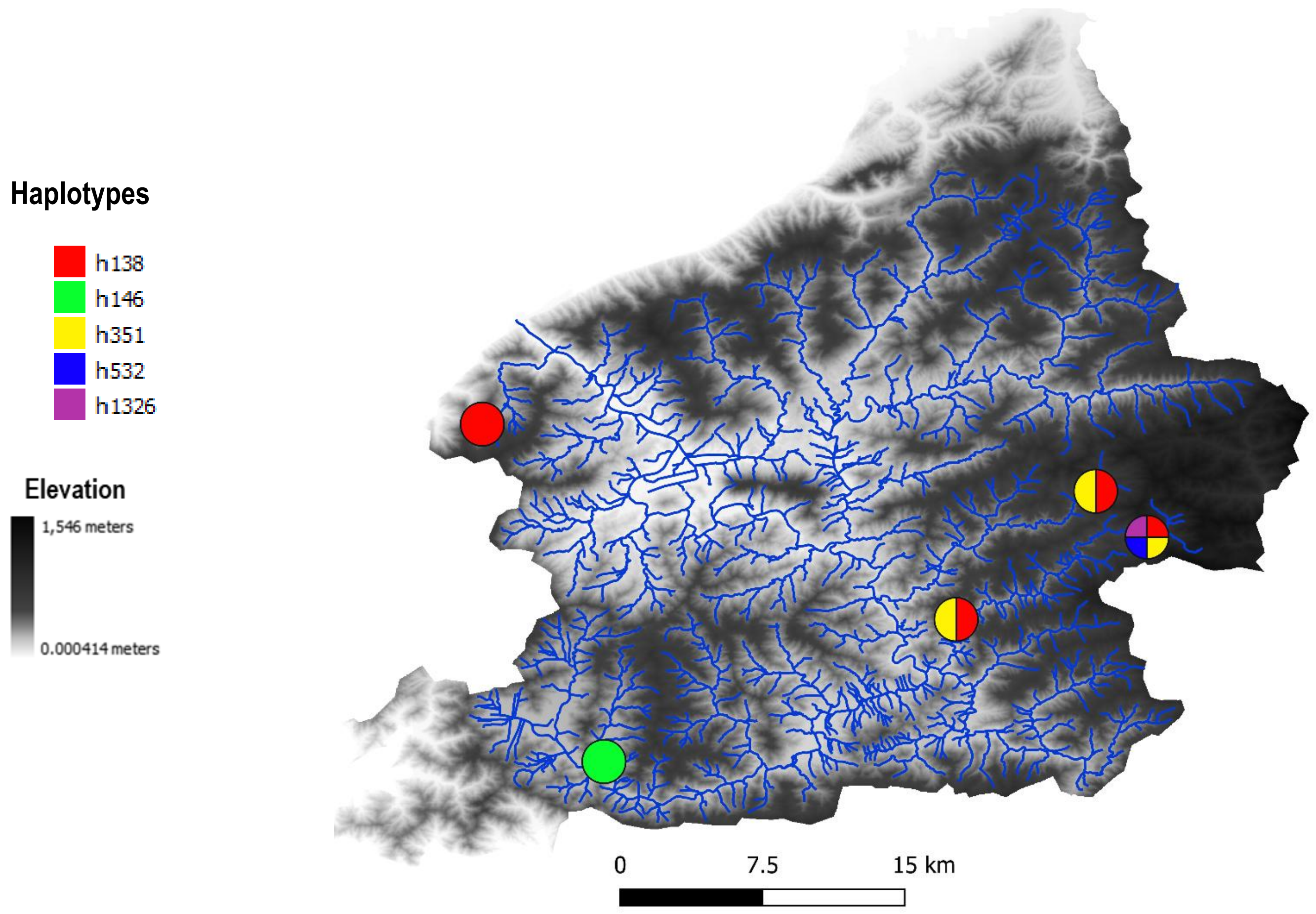


#### Haplotype network

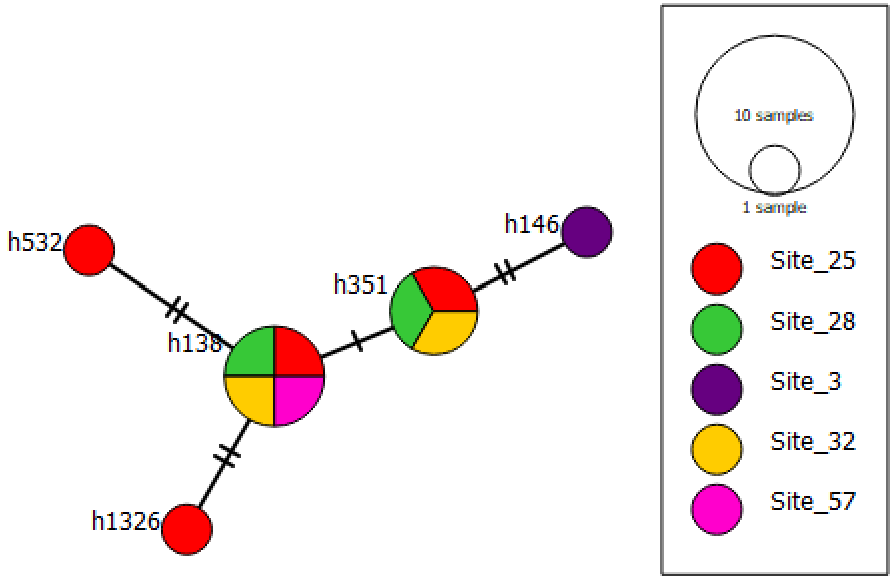

OTU 83

#### Phylogeography

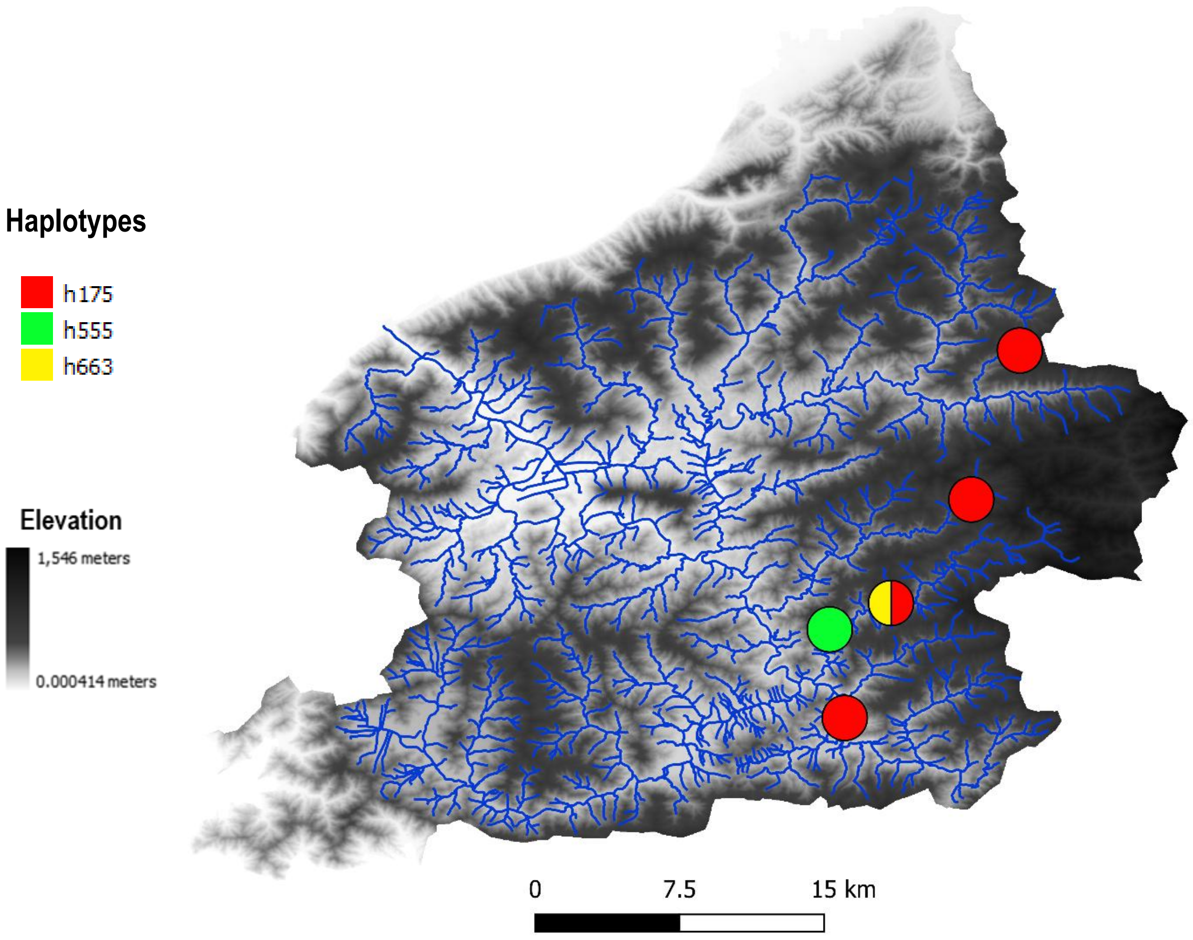


#### Haplotype network

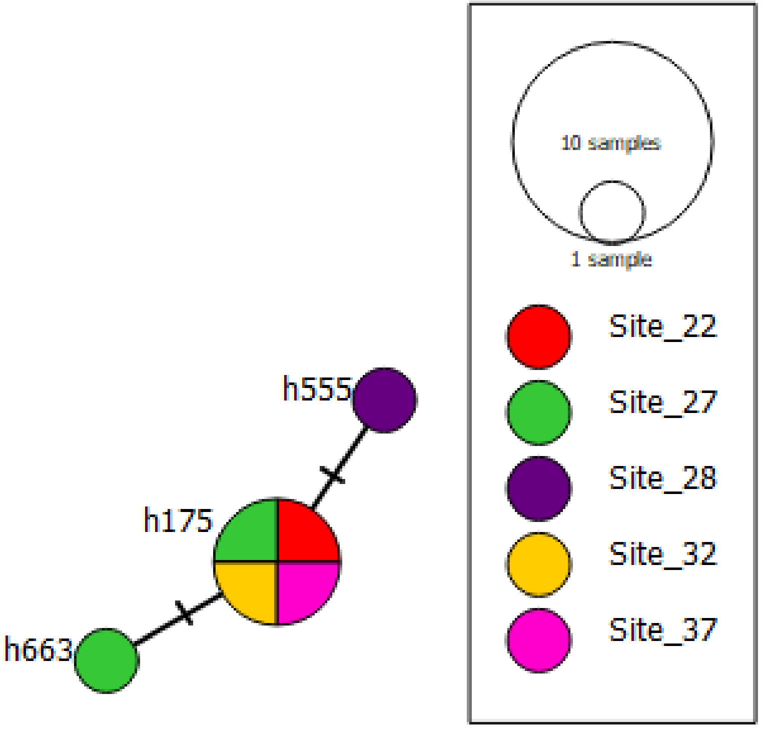

### OTU106

#### Phylogeography

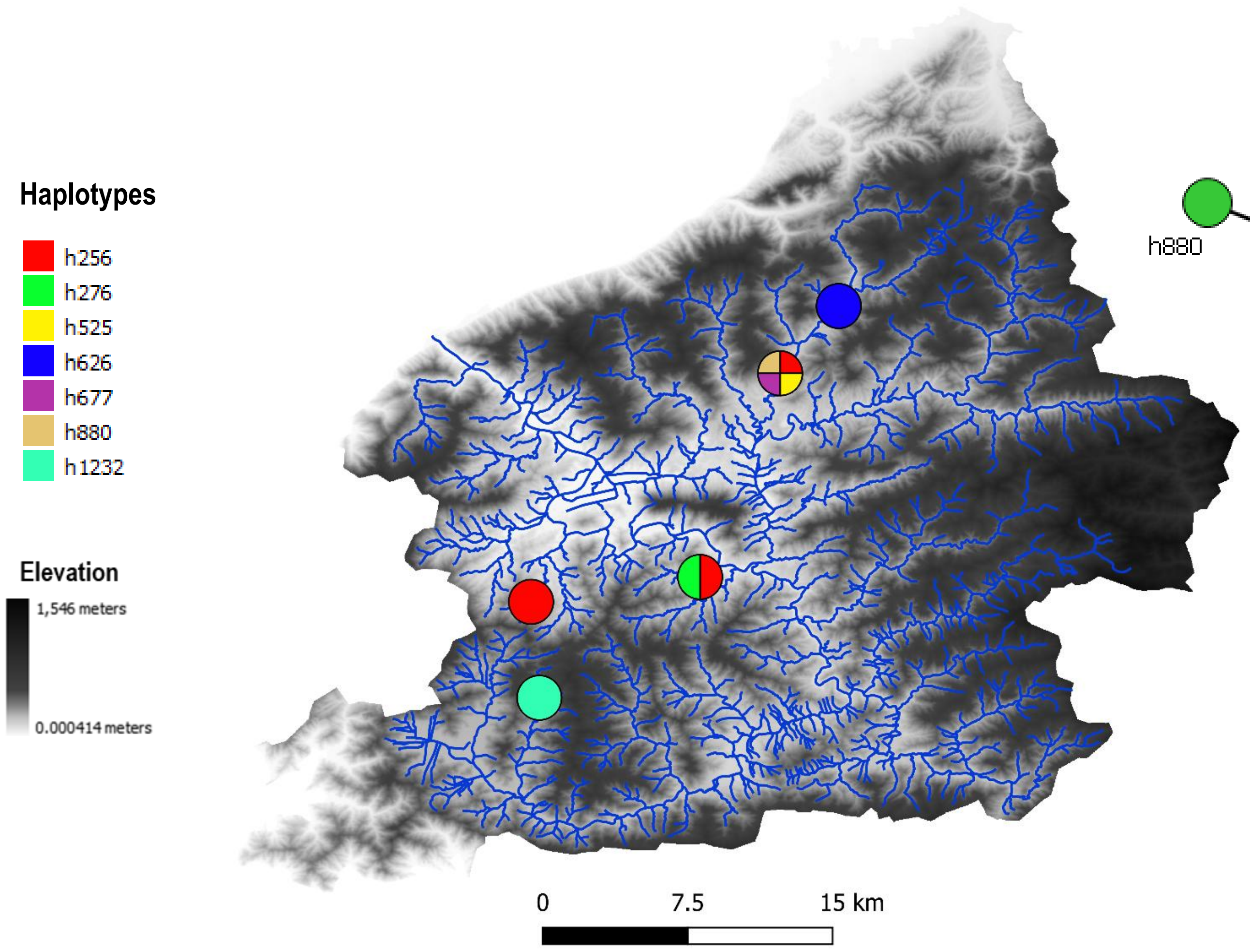


#### Haplotype network

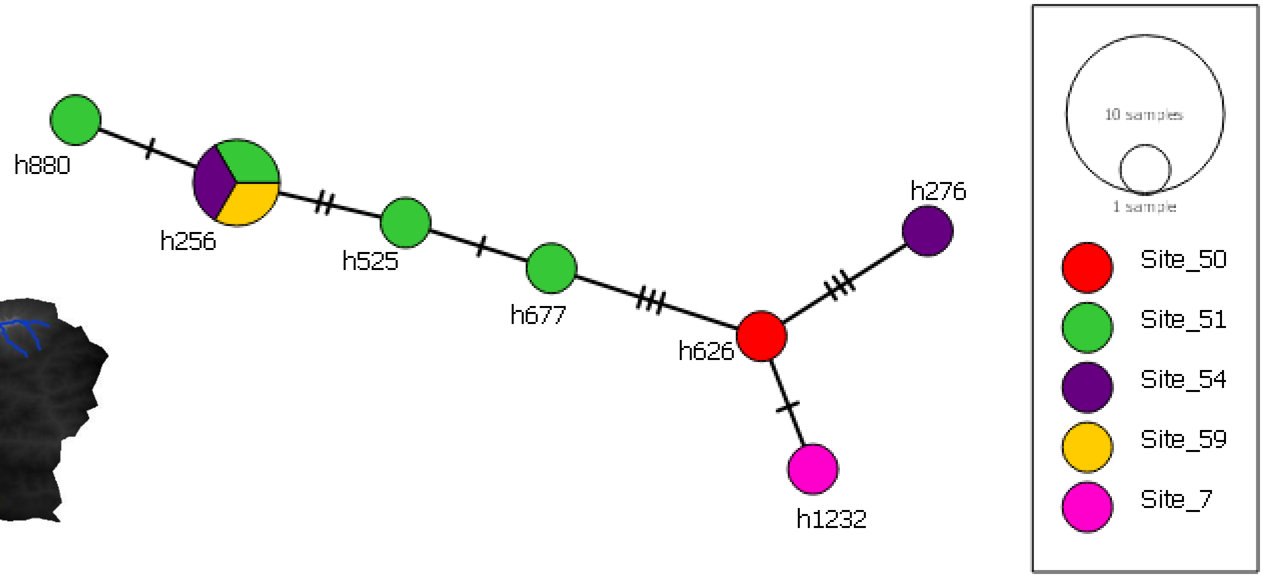

### OTU134

#### Phylogeography

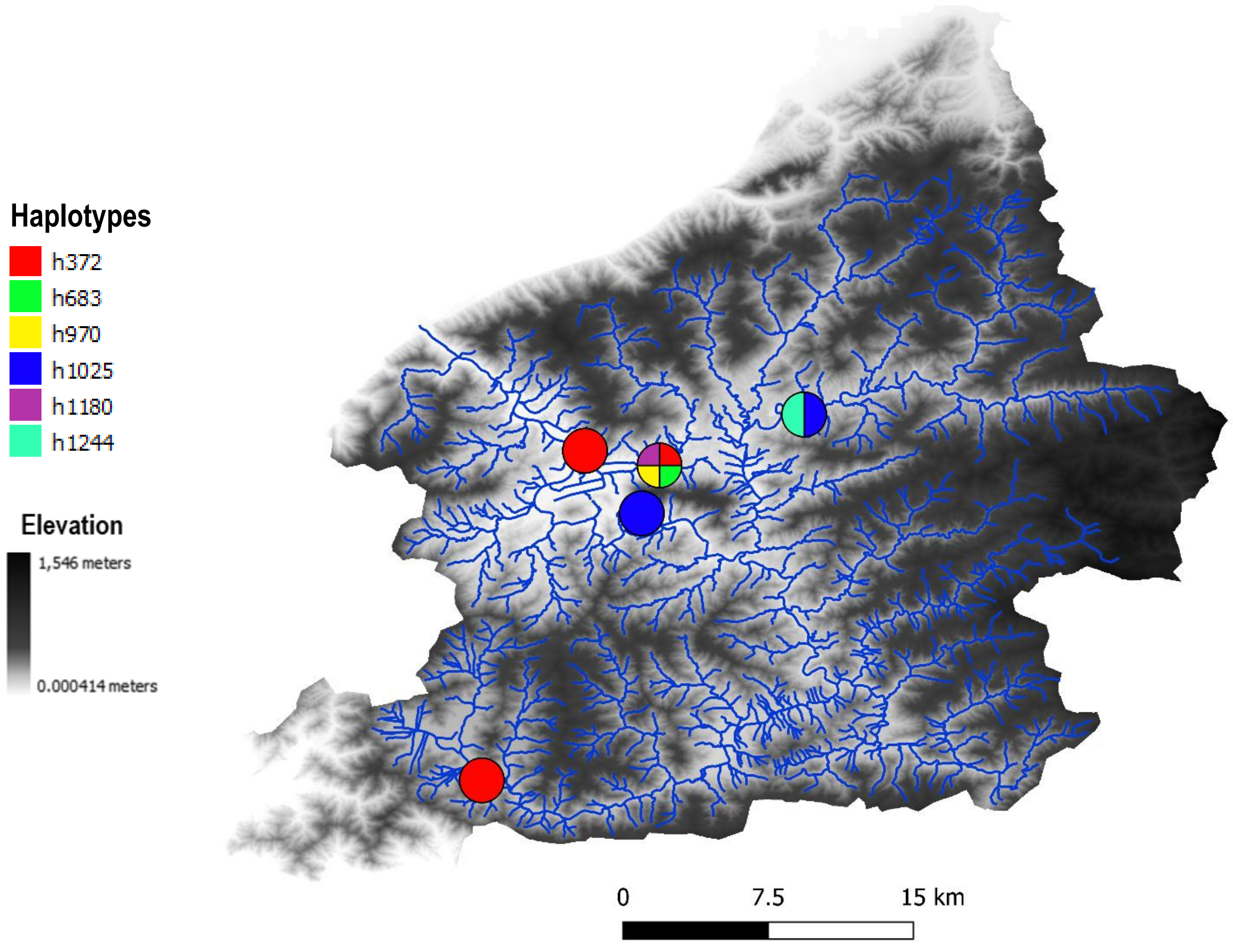


#### Haplotype network

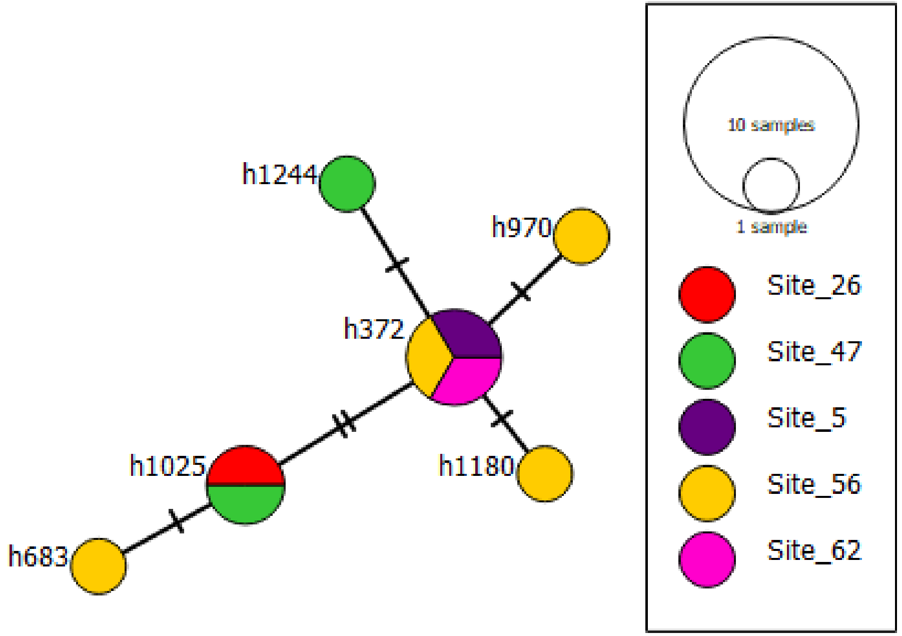

### OTU165

#### Phylogeography

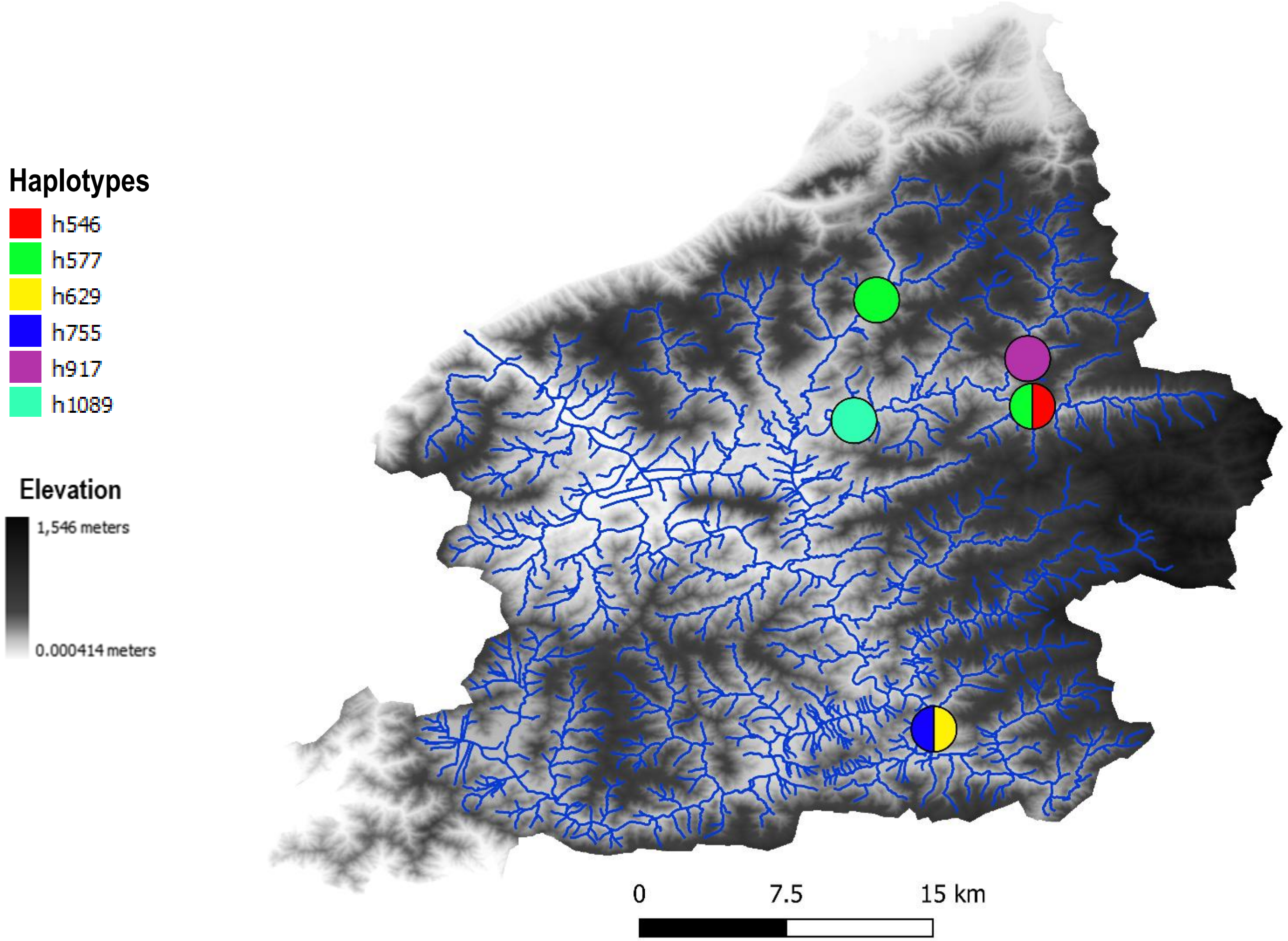


#### Haplotype network

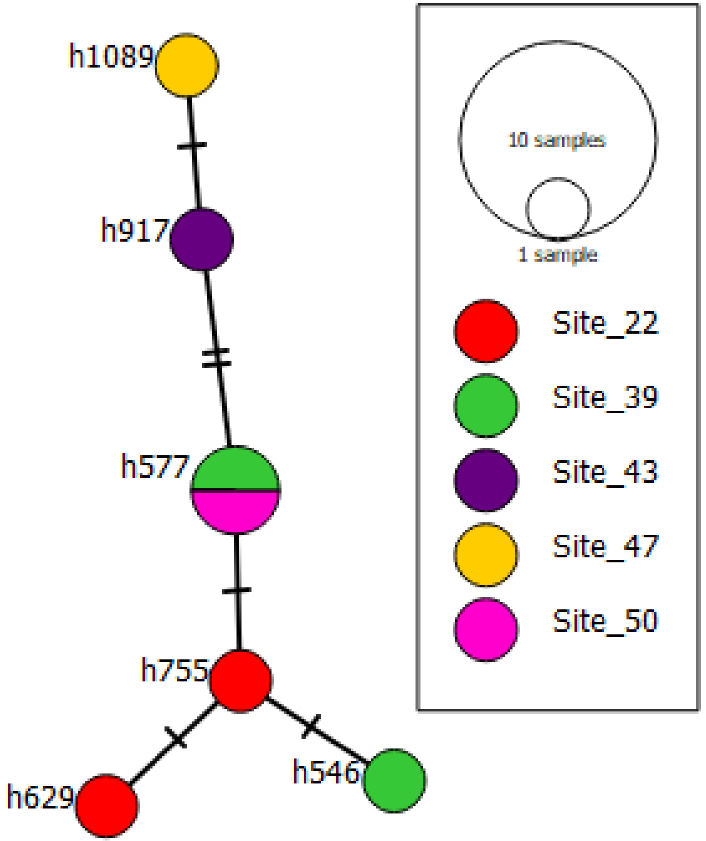

OTU 157

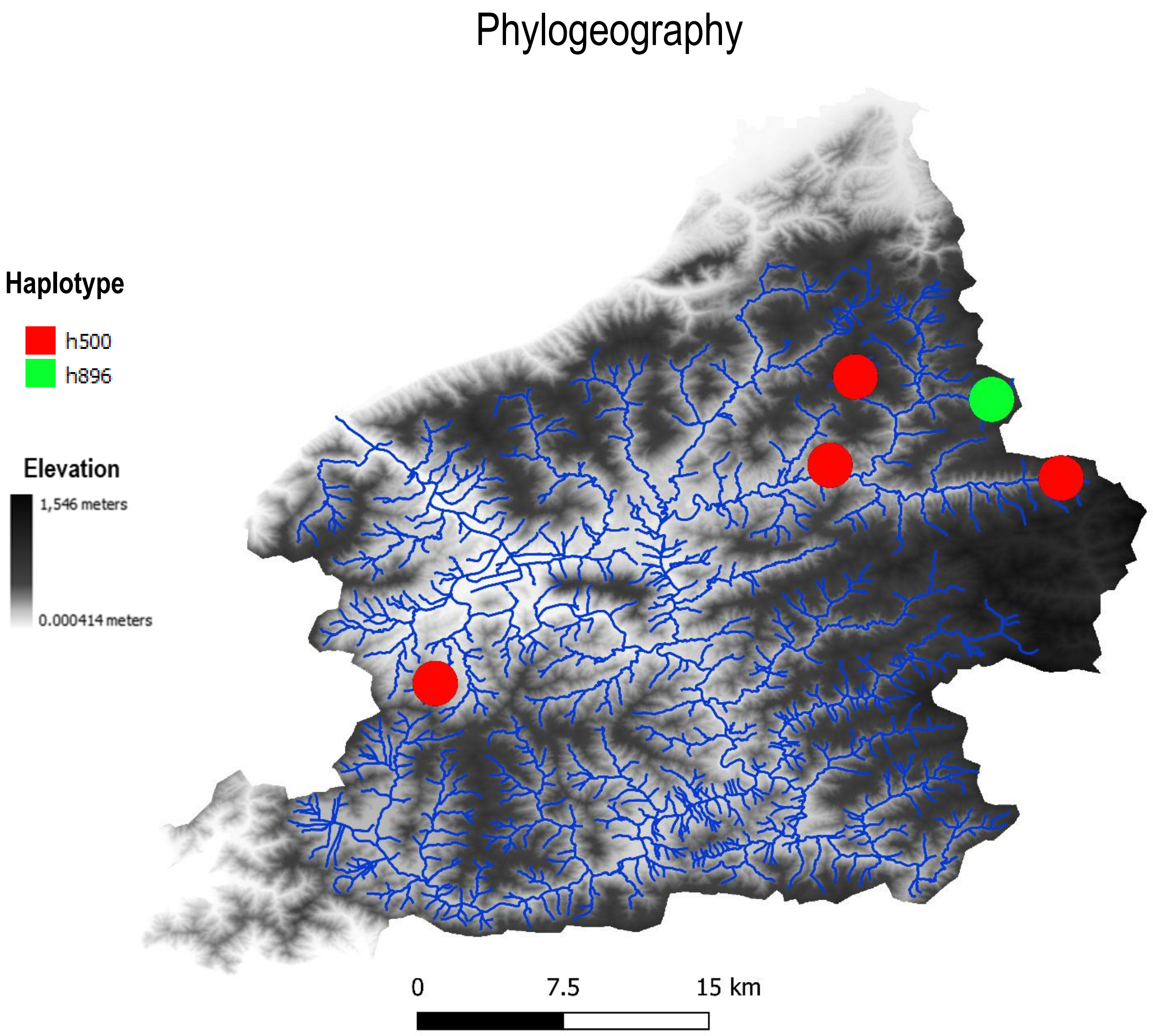

Phylogeography
Haplotype
h500
h896
Elevation
1,546 meters
0.000414 meters
0
7.5
15 km


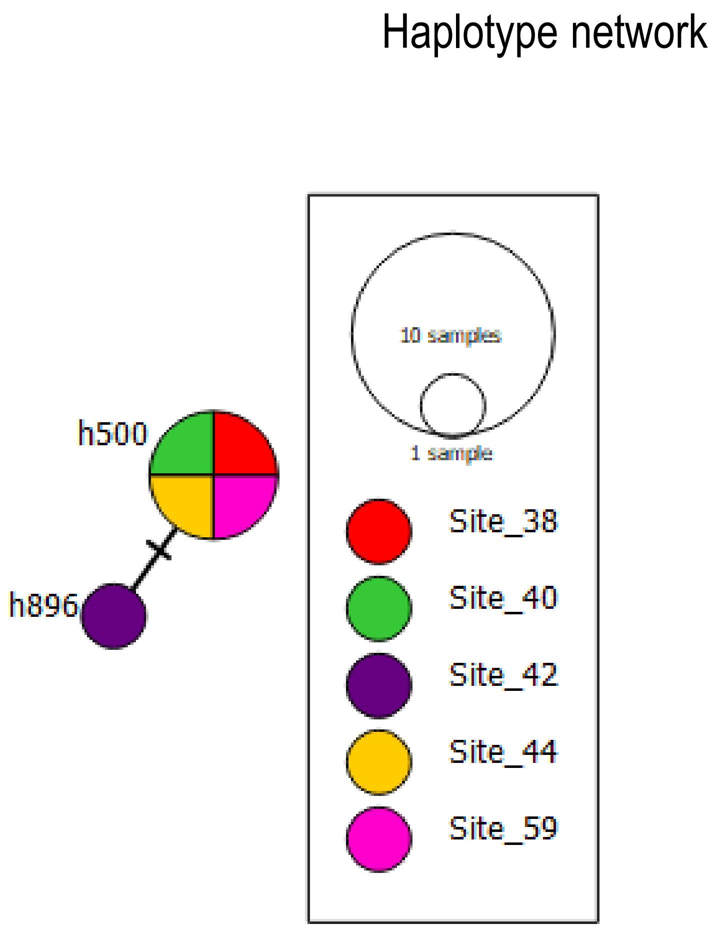

Haplotype network
h500
h896
10 samples
1 sample
Site_38
Site_40
Site_42
Site_44
Site_59

### OTU 94

#### Phylogeography

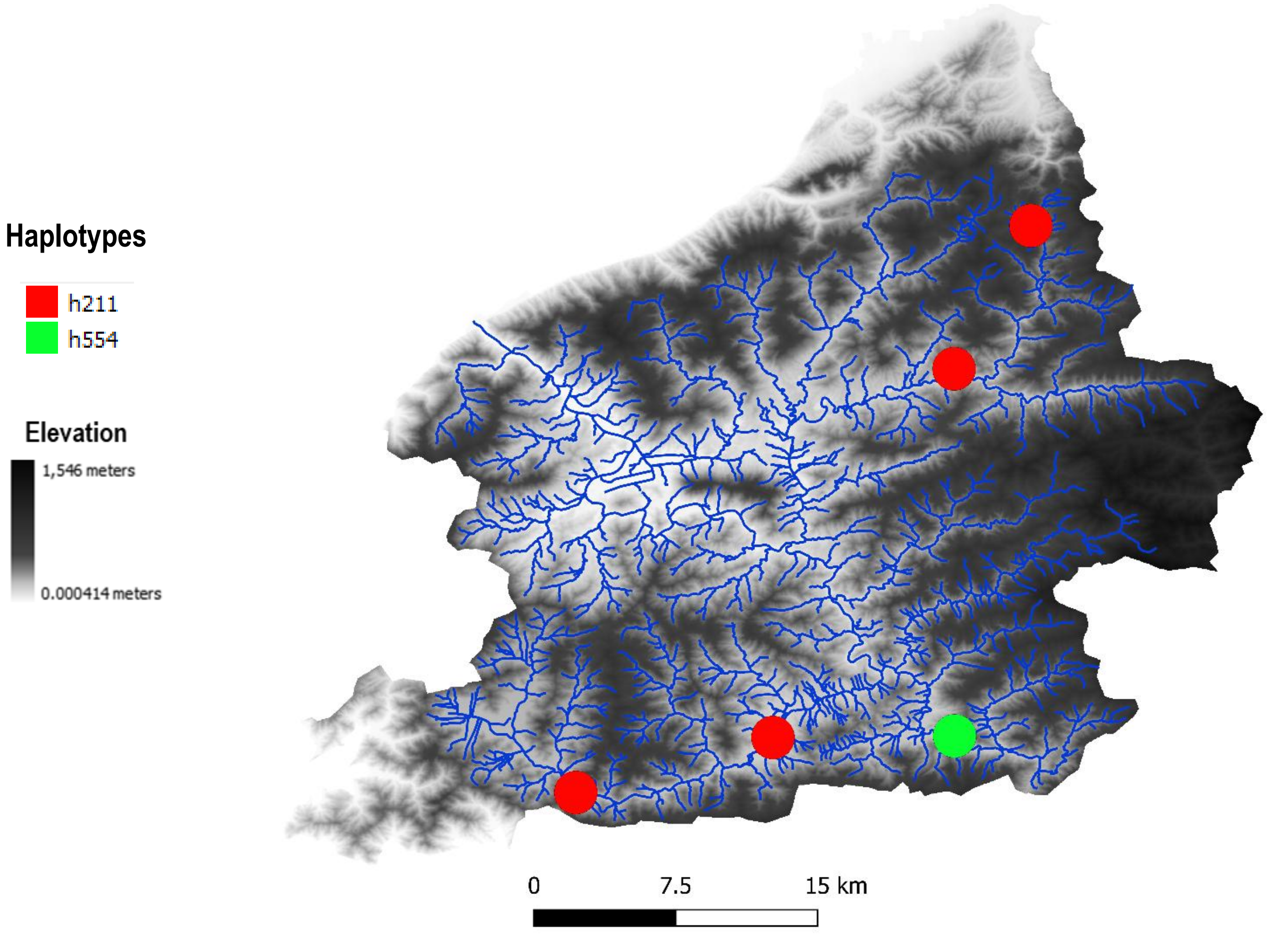


#### Haplotype network

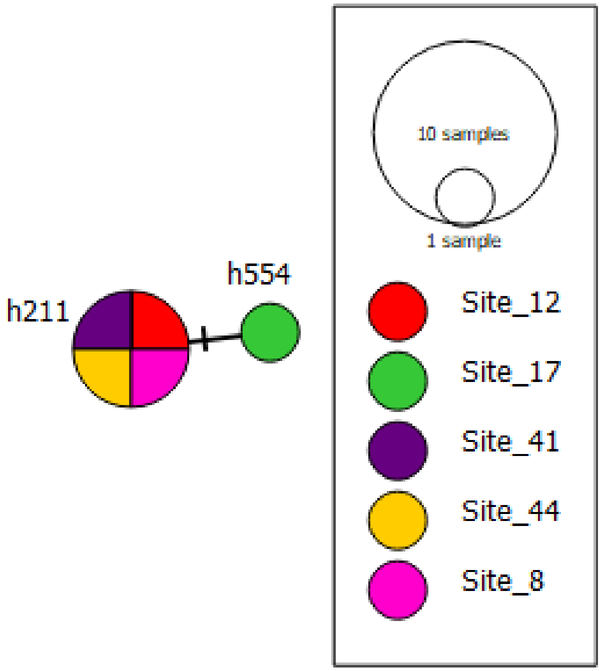

### OTU 105

#### Phylogeography

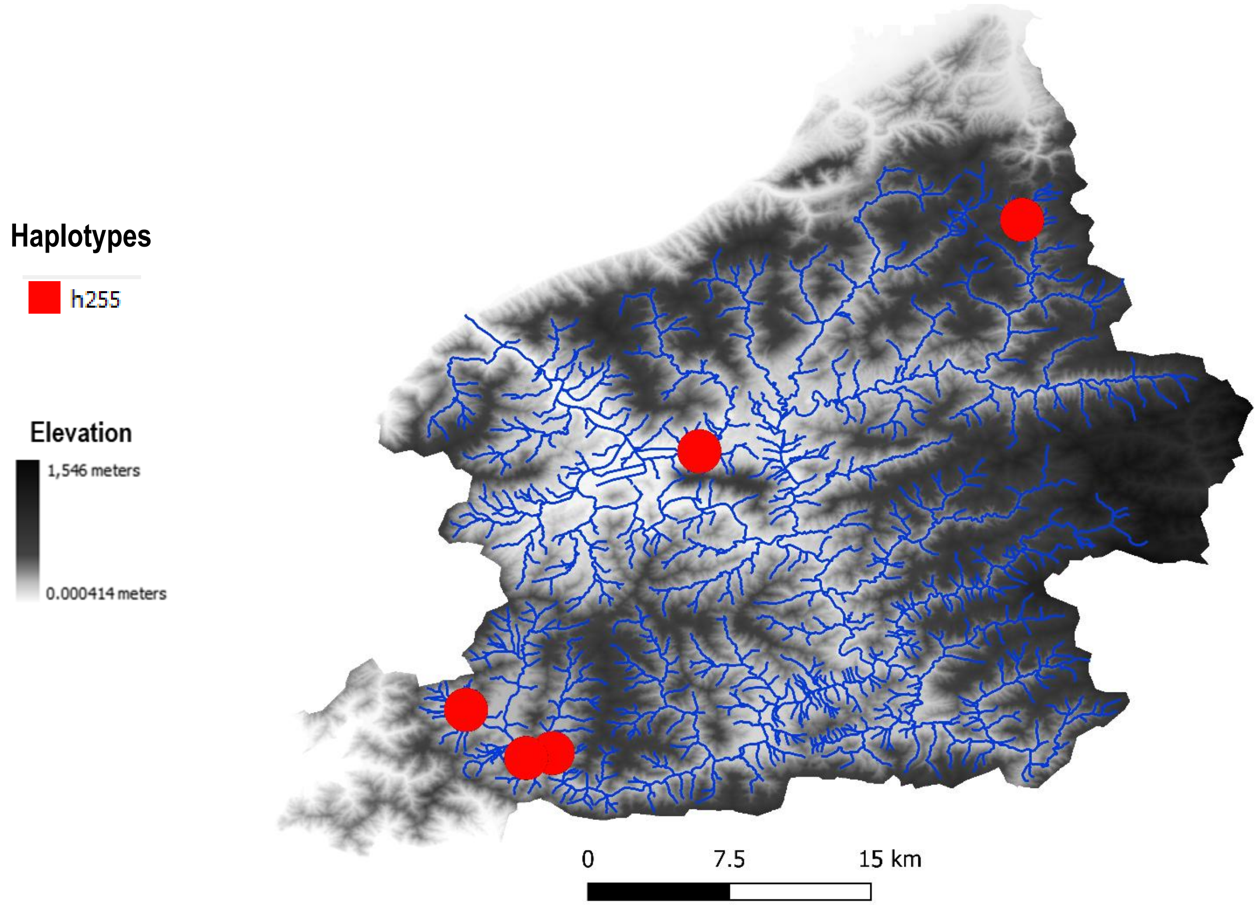


#### Haplotype network

### OTU 132

#### Phylogeography

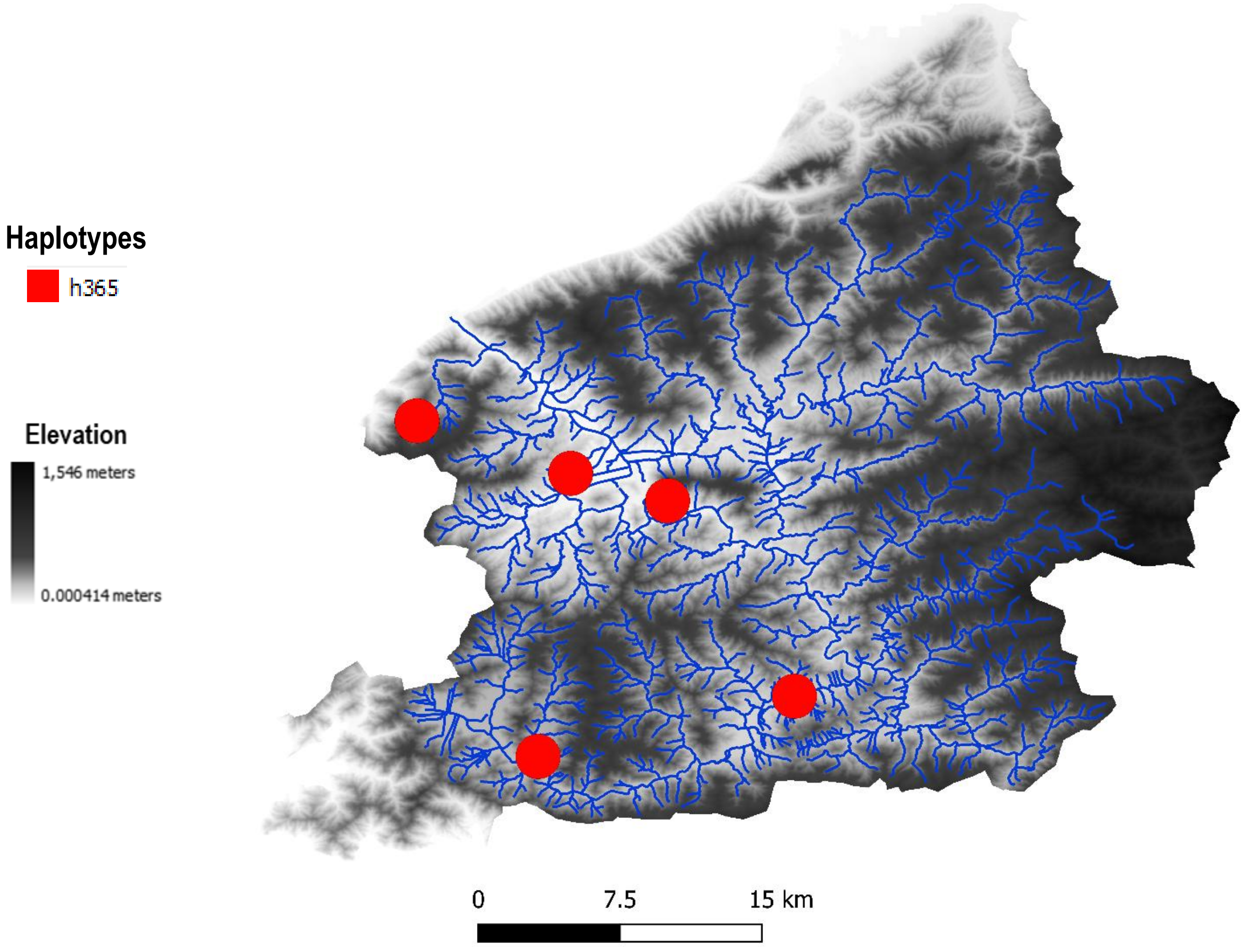


#### Haplotype network
