## supplementary fig. S1 for "Haplotype-level metabarcoding revealed the relationship between species occurrence and population genetic structure at the catchment scale in aquatic insects"

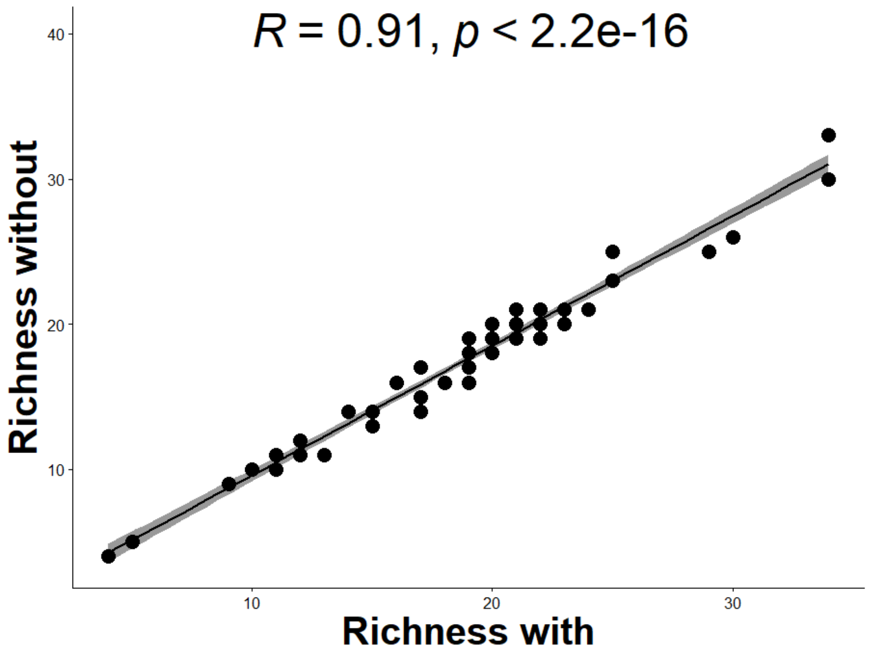


**Supplementray Fig. S1.** Correlation between OTU richness of data set including the six mixed OTUs (Richness with) and without the mixed six OTUs (Richness without)

(a)

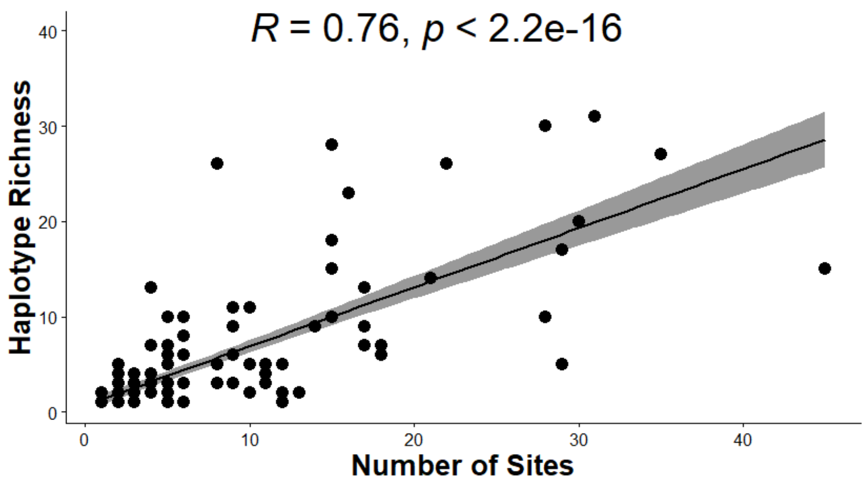


(b)

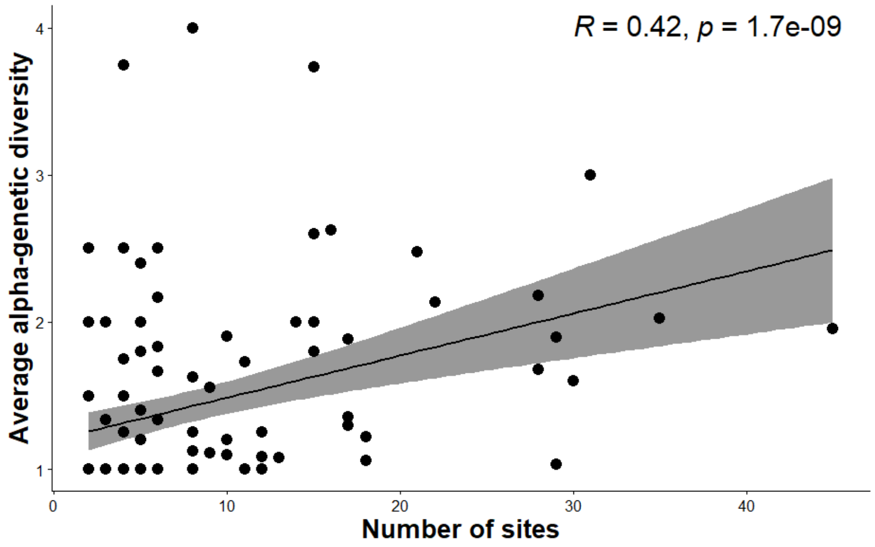


**Supplementary Figure S2**

(a) Correlation between number of sites and haplotype richness in all 194 OTUs
(b) correlation of number of sites and average α-genetic diversity in 117 OTUs present in at least two sites

**(a)**

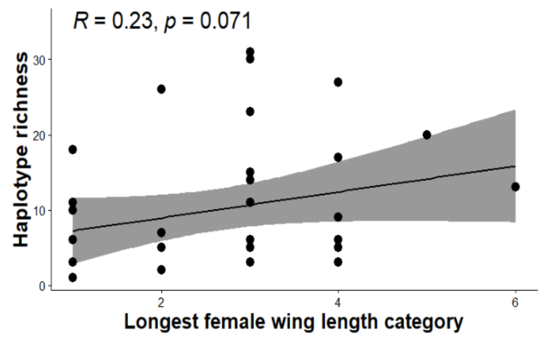


**(b)**

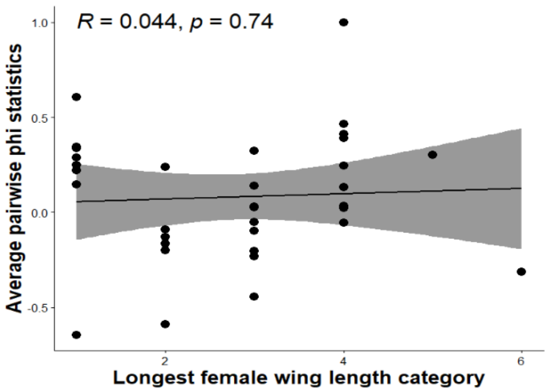


**(c)**

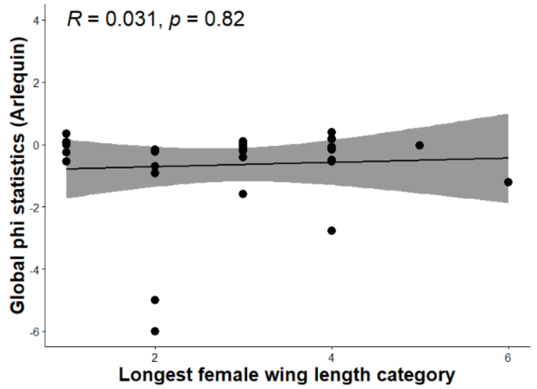


**Supplementary Figure S4**

**(a)** Correlation between female wing length and haplotype richness
**(b)** Correlation between female wing length and average pairwise ϕ-statistics
**(c)** Correlation between female wing length and global ϕ-statistics

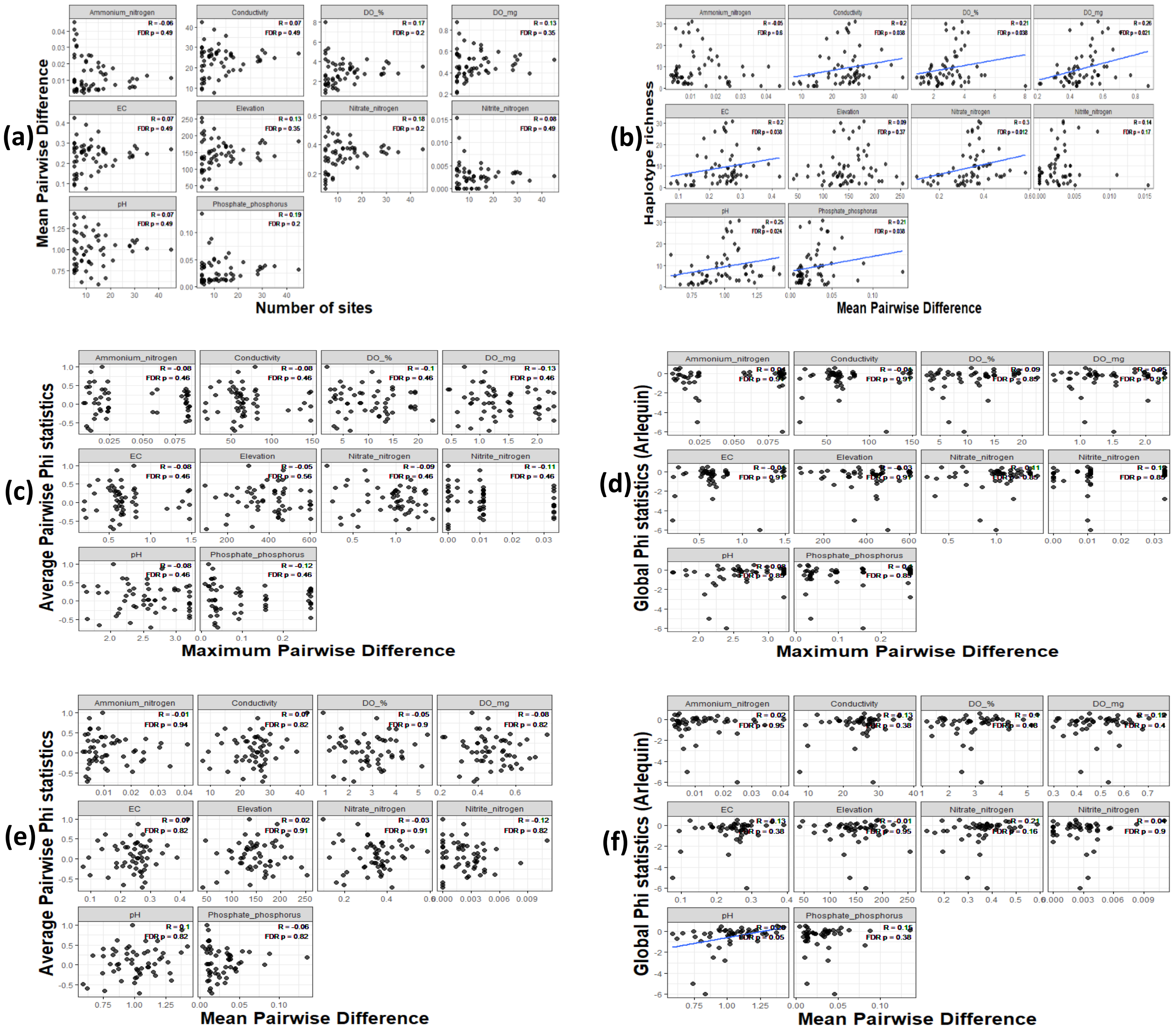


**Supplementary Figure S5**

(a) Correlation between number of sites and mean pairwise difference in all environmental parameters using 59 OTUs
(b) Correlation between haplotype richness and mean pairwise difference in all environmental parameters using 59 OTUs
(c) Correlation between average pairwise ϕ-statistics with maximum pairwise difference in all environmental parameters using 59 OTUs
(d) Correlation between global ϕ-statistics with maximum pairwise difference in all environmental parameters using 59 OTUs
(e) Correlation between average pairwise ϕ-statistics with mean pairwise difference in all environmental parameters using 59 OTUs
(f) Correlation between global ϕ-statistics with mean pairwise difference in all environmental parameters using 59 OTUs
