## supplementary protocol 1 for "Haplotype-level metabarcoding revealed the relationship between species occurrence and population genetic structure at the catchment scale in aquatic insects"

### Phenol-Chloroform Isoamyl (PCI) DNA Extraction using Bulk Community samples

Briefly, 300 µL of Phosphate Buffered Saline (PBS), 180 µL of ATL buffer (Qiagen), 20 µL of Proteinase K (Qiagen), and a sterile bead-beating ball were added to ethanol-dried macroinvertebrate bulk community samples. The mixture was homogenized at 1,500 RPM for 15 minutes using Shake Master Neo cell destructive equipment (BioMedical Sciences) and incubated overnight at 56°C. Afterward, 500 µL of PCI at a ratio of 25:24:1 was added and centrifuged at 1,200 RPM for 10 minutes. The supernatant was then treated with 100 µL of chloroform and centrifuged under the same conditions. DNA was precipitated by adding 30 µL of 3M sodium acetate (pH 5.2) and 660 µL of 99.5% ethanol and stored at -20°C for 2-3 hours. We centrifuged the sample at 12,000 RPM for 15 minutes to collect the DNA and washed it using 70% ethanol. After drying, the pellet was reconstituted with 50 µL of TE buffer and stored at -20°C until use.

**Supplementary Protocol 2**

**Two-step library preparation method (Elbrecht and Steinke, 2018) with modifications**

**Elbrecht, V., & Steinke, D. (2019). Scaling up DNA metabarcoding for freshwater macrozoobenthos monitoring. Freshwater Biology, 64(2), 380-387**

In the first round of PCR, we amplified the target gene from the three samples collected per site. The reaction mixture consisted of 3 µL of 5X Phusion HF buffer (New England Biolabs), 0.6 µL of 2.5 mM dNTPs, 0.45 µL of 50 mM MgCl2, 0.45 µL of 100% DMSO, 0.75 µL of 10 µM each of BF2 and BR2 primers, 0.15 µL of Phusion HF DNA polymerase (2000 units/mL) (New England Biolabs), and 0.5 to 1 µL of DNA (10 to 20 ng/µL). We added molecular-grade water to achieve a total volume of 15 uL. Initial denaturation was performed at 94°C for 3 minutes, followed by 26 to 31 cycles of 94°C for 30 seconds, 58°C for 30 seconds, and 72°C for 60 seconds, with a final extension at 72°C for 10 minutes.

The second round of PCR used fusion primers consisting of BF2/BR2, inline tags, and Illumina sequencing tails. The 15 µL reaction mixture included 3 µL of 5X Phusion HF buffer, 0.6 µL of 2.5 mM dNTPs, 0.6 µL of 50 mM MgCl2, 0.45 µL of 100% DMSO, 1.5 µL each of 5 µM forward and reverse primers, 0.25 µL of Phusion HF DNA polymerase (2000 units/mL), 0.5 µL of the first PCR product (pooled from 3 samples per site), and 6.6 µL of molecular grade water. The thermal cycling conditions were: initial denaturation at 94°C for 2 minutes, followed by 14 cycles of 94°C for 30 seconds, 68°C for 30 seconds, and 72°C for 1 minute, with a final extension at 72°C for 10 minutes.

**Supplementary protocol 3**

**Poisson Tree Process (PTP) analysis**

1. mtDNA COI sequences of closely related species of OTU suspected for being mixed species (> 0.03 genetic divergence) were downloaded from NCBI GenBank
2. Haplotype sequences of OTU suspected for being mixed species were aligned with the mtDNA COI sequences of closely related species using MUSCLE in MEGA
3. Best-fit evolutionary model was determined using MEGA
4. Neighbor-Joining (NJ) tree was created using the best-fit evolutionary model determined by MEGA
5. Posson Tree Process was conducted according to Zhang et al. (2013) (https://species.h-its.org/ptp/)
